# Functional connectome of arousal and motor brainstem nuclei in living humans by 7 Tesla resting-state fMRI

**DOI:** 10.1101/2021.10.18.464881

**Authors:** Kavita Singh, Simone Cauzzo, María Guadalupe García-Gomar, Matthew Stauder, Nicola Vanello, Claudio Passino, Marta Bianciardi

## Abstract

Brainstem nuclei play a pivotal role in many functions, such as arousal and motor control. Nevertheless, the connectivity of arousal and motor brainstem nuclei is understudied in living humans due to the limited sensitivity and spatial resolution of conventional imaging, and to the lack of atlases of these deep tiny regions of the brain. For a holistic comprehension of sleep, arousal and associated motor processes, we investigated in 20 healthy subjects the resting-state functional connectivity of 18 arousal and motor brainstem nuclei in living humans. To do so, we used high spatial-resolution 7 Tesla resting-state fMRI, as well as a recently developed in-vivo probabilistic atlas of these nuclei in stereotactic space. Further, we verified the translatability of our brainstem connectome approach to conventional (e.g. 3 Tesla) fMRI. Arousal brainstem nuclei displayed high interconnectivity, as well as connectivity to the thalamus, hypothalamus, basal forebrain and frontal cortex, in line with animal studies and as expected for arousal regions. Motor brainstem nuclei showed expected connectivity to the cerebellum, basal ganglia and motor cortex, as well as high interconnectivity. Comparison of 3 Tesla to 7 Tesla connectivity results indicated good translatability of our brainstem connectome approach to conventional fMRI, especially for cortical and subcortical (non-brainstem) targets and to a lesser extent for brainstem targets. The functional connectome of 18 arousal and motor brainstem nuclei with the rest of the brain might provide a better understanding of arousal, sleep and accompanying motor function in living humans in health and disease.

## 1. Introduction

Substantial research over several decades, based on lesion/stimulation studies in animals, as well as sleep and coma studies in humans and animals, has identified distinct neural populations in the brainstem promoting wakeful arousal, and/or associated motor function (Table 1). These include nuclei of median raphe (MnR), paramedian raphe (PMnR) and dorsal raphe (DR) substantia nigra-subregion1 (SN1, compatible with pars reticulata), substantia nigra-subregion2 (SN2, compatible with pars compacta), (Datta et al., 1998; Lima et al., 2008, 2007; Olszewski and Baxter, 1954) caudal-rostral linear raphe (CLi-RLi), periaqueductal gray (PAG), mesencephalic reticular formation (mRt), cuneiform nucleus (CnF), isthmic reticular formation (isRt), and pontine reticular formation (PnO-PnC) (Moruzzi and Magoun, 1949; Parvizi and Damasio, 2001), locus coeruleus (LC) (Parvizi and Damasio, 2001; Saper et al., 2010, 2001) pedunculotegmental (PTg) and laterodorsal tegmental nucleus-central gray of the rhombencephalon (LDTg-CGPn) (Parvizi and Damasio, 2001; Saper et al., 2010, 2001), red nucleus-subregion1 (RN1), Red nucleus-subregion2 (RN2), subcoeruleus nucleus (SubC), inferior olivary nucleus (ION) (Merel et al., 2019).

With the goal of elucidating how these nuclei individually and collectively act to promote and maintain wakefulness and various arousal states, animal studies have mapped the connections among these nuclei (Oh et al., 2014; Olszewski and Baxter, 1954; Saper et al., 2010, 2001) and other arousal regions such as the thalamus, hypothalamus and basal forebrain. Based on these studies, it is now increasingly clear that arousal is controlled by a large, distributed networks spanning multiple subcortical and cortical regions, many of which are closely associated with somato-motor and autonomic regulation (Liu and Dan, 2019).

To fully elucidate wakeful arousal and associated motor processes, a comprehensive investigation of functional connections of the nuclei involved is needed. Moreover, with the ever-increasing evidence of overlapping functions of several regions (Elvsåshagen et al., 2020; Eser et al., 2018), it is becoming imperative to investigate affected networks rather than single regions. However, the connectivity of arousal brainstem nuclei is understudied in living humans (Bär et al., 2016; Beliveau et al., 2015; Bianciardi et al., 2016; Coulombe et al., 2016), due to the limited sensitivity and spatial resolution of conventional imaging and to the lack of atlases of these deep tiny regions of the brain.

To this end, we investigated the resting-state functional connectivity of 18 arousal and motor brainstem nuclei in living humans by the use of high-sensitivity and high spatial-resolution 7 Tesla resting-state fMRI, as well as a recently developed *in-vivo* probabilistic atlas of arousal and motor brainstem nuclei in stereotactic (Montreal-Neurological-Institute —MNI) space (Bianciardi et al., 2018, 2015; Garcia-Gomar MG et al., 2021; Singh et al., 2021). For this, we used brainstem nuclei (stated above) as seeds and cortical and sub-cortical regions encompassing the whole grey matter, as targets (Li et al., 2015; Rosen and Halgren, 2021). We also used as targets brainstem nuclei involved in other functions such as autonomic and sensory, available in the literature (Bianciardi et al., 2016, 2018; García-Gomar et al., 2019a; Garcia-Gomar et al., 2021): Superior Colliculus (SC) (Lee and Groh, 2012), Inferior Colliculus (IC) (Nieuwenhuys et al., 2008a), Ventral Tegmental Area-Parabrachial Pigmented Nucleus (VTA-PBP) (Halliday et al., 2012), Microcellular Tegmental Nucleus-Prabigeminal nucleus (MiTg-PBG), Lateral Parabrachial Nucleus (LPB), Medial Parabrachial Nucleus (MPB) (Kaur et al., 2013a), Vestibular nuclei complex (Ve) (Goldberg et al., 2012), Parvocellular Reticular nucleus Alpha (PCRtA) (Chai et al., 1988), Superior Olivary Complex (SOC) (Illing et al., 2000), Superior Medullary Reticular formation (sMRt), Viscero-Sensory Motor nuclei complex (VSM) (Mutolo, 2017), Inferior Medullary Reticular formation (iMRt), Raphe Magnus (RMg), Raphe Obscurus (ROb) and Raphe Pallidus (RPa) (Hornung, 2003)).

Further, we verified the translatability of our brainstem connectome approach to conventional (e.g. 3 Tesla) fMRI. Translatability to clinical studies is crucial to allow the investigation of brainstem arousal mechanisms in disease, including sleep disorders, disorders of consciousness, and psychiatric disorders involving perturbations of arousal (Bryant et al., 2000) for example, hyperarousal in schizophrenia, addiction, and generalized anxiety disorder (GAD) (Carlsson, 1995), or hypoarousal in aggressiveness and attention deficit hyperactivity disorder (ADHD) (Haller et al., 2005; Miano et al., 2006). This could further benefit the investigation of motor mechanisms in Parkinson’s disease, Lewy body dementia, spinocerebellar ataxia, and supranuclear palsy.

## 2. Methods

### 2.1 : Data Acquisition

20 right-handed healthy subjects (10 males and 10 females; age 29.5±1.1 years) participated in two MRI sessions, a 7 Tesla MRI session and 3 Tesla MRI session (Siemens Healthcare, Erlangen, Germany). The session order was randomized across subjects. The study protocol was approved by the Massachusetts General Hospital Institutional Review Board; written informed consent was obtained from participants. During the MRI image acquisition sessions, subjects were asked to lie down in the scanner and be as still as possible. Foam pads were placed beneath subject’s neck covering the space between the subject and the MRI detector array to minimize head movements. A custom-built 32-channel receive coil and volume transmit coil was used at 7 Tesla. We used a custom-built 64-channel receive coil and volume transmit coil at 3 Tesla (Keil et al., 2010). For both 7 Tesla and 3 Tesla acquisition, to account for physiological related signal fluctuations, during fMRI acquisition, we also recorded the timing of cardiac and respiratory cycles by the use by a piezoelectric finger pulse sensor (ADInstruments, Colorado Springs, CO, USA) and a piezoelectric respiratory bellow (UFI, Morro Bay, CA, USA) positioned around the chest, respectively. The recording was performed using a PowerLab data acquisition system (PowerLab 4/SP, ADInstruments, New Zealand) and LabChart (LabChart, ADInstruments, New Zealand), with a sampling rate of 1000 Hz. The same acquisition system also recorded the MR scanner trigger pulses synchronized with the acquisition of each image volume.

During the study, we asked participants to stay awake with their eyes closed during resting-state fMRI acquisitions. We checked their sleep state by asking buzzer response question before and after each run.

#### 2.1a: 7 Tesla MRI session

For each subject, on a 7 Tesla MRI scanner (Magnetom, Siemens Healthineers, Erlangen, Germany) we acquired three runs of gradient-echo echo-planar-images (EPIs) during resting state with the eyes closed, with the following parameters: isotropic voxel-size = 1.1 mm, matrix-size = 180×240, GRAPPA factor = 3, nominal echo-spacing = 0.82 ms, bandwidth = 1488 Hz/Px, N. slices = 123, slice orientation = sagittal, slice-acquisition order = interleaved , echo-time (TE) = 32 ms, repetition-time (TR) = 2.5 s , flip angle (FA) = 75°, simultaneous-multi-slice factor = 3, N. repetitions = 210, phase-encoding direction = anterior-posterior, acquisition-time per-run = 10’07”. To correct for geometric distortion, a 2.0 mm isotropic resolution fieldmap was acquired with the following parameters: matrix-size = 116x132, bandwidth = 1515 Hz/Px, number of slices = 80, slice orientation = sagittal, slice-acquisition order = interleaved, TE1/TE2 = 3.00 ms/4.02 ms, TR = 570.0 ms, FA = 36°, simultaneous-multi-slice factor = 3, phase-encoding direction = anterior-posterior.

#### 2.1 b: 3 Tesla MRI session

To access clinical translatability of our 7 Tesla functional connectome, we scanned the same subjects at 3 Tesla (‘Connectom’ scanner, Siemens Healthineers) for multi-echo MPRAGE (MEMPRAGE), resting state fMRI and fieldmap image acquisition. Of note, we used conventional sequences for 3 Tesla MRI acquisition on a non-conventional (‘Connectom’) 3 Tesla scanner. We acquired an anatomical T_1_-weighted MEMPRAGE image with isotropic voxel-size = 1 mm, TR = 2.53 s, TEs = 1.69,3.5,5.3,7.2 ms, inversion-time = 1.5 s, FA = 1.5 s, FOV = 7°, bandwidth = 650Hz/pixel, GRAPPA-factor = 3, slice orientation = sagittal, slice-acquisition order = anterior-posterior, acquisition time = 4′28”.

We also acquired a single run of functional gradient-echo EPIs with isotropic voxel-size = 2.5 mm, matrix-size = 100×215, GRAPPA factor = 2, nominal echo-spacing = 0.5 ms, readout bandwidth = 2420 Hz/Px, N. slices = 64, slice orientation = transversal, slice-acquisition order = interleaved, TE = 30 ms, TR = 3.5 s, FA = 85°, GRAPPA factor = 2, Number of repetitions = 150, phase-encoding direction = anterior-posterior, acquisition-time = 9’06”.

To correct for geometric distortions, we acquired a fieldmap with isotropic voxel-size = 2.5 mm, FOV = 215x215, bandwidth = 300 Hz/Px, N. slices = 128, slice orientation = sagittal, slice-acquisition order = interleaved, TE1/TE2 = 4.92 ms/4.02 ms, TR = 849.0 ms, FA = 85°, phase-encoding direction = anterior-posterior, acquisition time = 2’19’’.

### 2.2 : Data processing

#### 2.2a: MEMPRAGE and Fieldmap Processing

For each subject, we computed the root-mean-square of the MEMPRAGE image across echo times. We then rotated it to standard orientation (“RPI”), bias field corrected (SPM) (Frackowiak et al., 2004), brain extracted and cropped the lower slices (*FSL 5.0.7 tools-FMRIB Software Library, FSL, Oxford, UK*). The preprocessed MEMPRAGE was parcellated with Freesurfer (Destrieux et al., 2010) to generate cortical and sub-cortical targets. For registration purposes, the MEMPRAGE images were averaged across 20 subjects to build an optimal template using the Advanced Normalization Tool (ANTs, Philadelphia, PA, United States). The optimal template was then registered to the MNI152_1mm template (Grabner et al., 2006) through an affine transformation and a nonlinear warp. The transformation matrices from individual subjects’ MEMPRAGE space to optimal template space and from optimal template space to MNI space were concatenated to obtain the full coregistration warp from single-subject MEMPRAGE to MNI space.

For each subject, the magnitude image of the fieldmap was reoriented, bias field corrected (SPM), and brain extracted [*FSL 5.0.7 tools-FMRIB Software Library, FSL, Oxford, UK*]. The brain mask generated after brain extraction was precisely inspected and manually edited to optimally include the oral pons, medulla, and to remove any unwanted non-brain voxel. Similarly, the phase image of the field map was reoriented to standard space, brain extracted (using the edited magnitude image mask), converted to radians per second, and smoothed with a 3D Gaussian kernel (4 mm sigma). The magnitude image was then warped to match the phase image (FSL-Fugue) and used to linearly register the phase image to the resting state fMRI image (FSL-FLIRT). The phase image registered to the fMRI was finally used to compute the warp for distortion correction using FSL program Fugue.

#### 2.2b: 7 Tesla fMRI Preprocessing

Physiological noise correction was done in each resting state fMRI run using custom-built Matlab function of RETROICOR (Glover et al., 2000) adapted to our slice acquisition sequence. For each subject, EPIs were slice-timing corrected and reoriented to standard space. (FSL 5.0.7 tools-FMRIB Software Library, FSL, Oxford, UK). fMRIs were motion corrected to the first volume of the first fMRI run (using Freesurfer, which also yielded six nuisance regressors to be used as covariates later). The time averaged fMRI of the first run was bias field corrected (SPM), distortion corrected, and used as a reference fMRI volume for coregistration to the MEMPRAGE image. This EPI was coregistered to the MEMPRAGE using a two-step process including an affine coregistration (AFNI program 3dAllineate) and a nonlinear one (AFNI 3dQwarp) (AFNI, Bethesda, MD, USA [Cox, 1996]). The nonlinear approach was iterative with progressively finer refinement levels and edges enhancement. For 17 subjects out of 20, 3 refinement levels were used, while 4 were used for the remaining 3 subjects for satisfactory results. To reduce the number of spatial interpolations and the subsequent loss of spatial definition, the affine and nonlinear coregistrations of the fMRIs to the MEMPRAGE image were applied in a single step together with motion and distortion correction (AFNI-3dNwarpApply). For the latter we used the warps computed from the fieldmap (described above).

Further, from the fMRI time-course we regressed out six rigid-body motion time-series nuisance regressors (computed during Freesurfer motion-correction), a regressor describing respiratory volume per unit time convolved with a respiration response function (Birn et al., 2008) (computed using Matlab), a regressor describing heart rate convolved with a cardiac response function (Chang et al., 2009) (computed using Matlab) and five regressors modelling the signal in cerebrospinal fluid (CSF). These CSF regressors were extracted from a principal-components analysis performed on a mask of the CSF (5.3 × 6.5 × 7.5 cm cuboid) neighboring the brainstem, defined by thresholding the EPIs standard deviation over time (please report the threshold here). Preciously, this CSF mask included CSF-filled spaces neighboring the brainstem, mainly the inferior portion of the third ventricle, the cerebral aqueduct and the fourth ventricle. Previous studies (Satpute et al., 2013) showed that removal of the signal of the cerebral aqueduct and ventricles adjacent to the brainstem is important for the functional connectivity analysis of brainstem nuclei lining the CSF, such as the periaqueductal gray, the dorsal raphe, and keeping in mind the nuclei in current study vestibular nuclei and inferior olivary nuclei. The BOLD signal fluctuations were converted to percentage signal changes with respect to the mean by dividing the data by temporal signal mean and multiplying by 100. High and low bandpass filtering was applied (between 0.01 Hz and 0.1 Hz) (AFNI-3dBandpass). Seminal studies showed functional connectivity signals of interest typically range between 0.01 − 0.1 Hz (Biswal et al., 1995; Cordes et al., 2001; Demirci et al., 2009; Salvador et al., 2005). Scanner drifts are mainly present at frequencies below 0.01 Hz (Bianciardi et al., 2009) and physiological noise derived from respiratory and cardiac function can be also found at comparatively higher frequencies (> 0.1 Hz) (Cordes et al., 2001; Lowe et al., 1998; Thomas et al., 2002). Filter specification and implementation employed in fMRI connectivity literature varies, however majority of studies demonstrate that the 0.01–0.1 Hz frequency range had the greatest power in revealing the underlying connectivity (Auer, 2008; Uddin et al., 2009; Zhong et al., 2009). Based on these evidences, we used most standard bandpass filter with cutoff frequencies in the range of 0.01-0.1. Finally, any residual temporal mean was removed (i.e. we subtracted the overall signal mean over time in each voxel), and the three resting state fMRI runs were concatenated. Note that we did not perform global signal regression because it alters the distribution of regional signal correlations in the brain and can induce artefactual anticorrelations (Aquino et al., 2020; Murphy et al., 2009).

##### 2.2 b (i) Defining seed and target regions for 2D connectome generation

As seed regions, we used the labels of 18 brainstem nuclei (13 bilateral and 5 unilateral, for a total of 31 seed regions) involved in arousal and motor function (developed by this group (Bianciardi et al., 2018, 2015; Garcia-Gomar MG et al., 2021; Singh et al., 2021), see also Table 1:

• serotonergic nuclei of median raphe (MnR), paramedian raphe (PMnR) and dorsal raphe (DR) (Parvizi and Damasio, 2001; Saper et al., 2010, 2001);
• mesolimbic dopaminergic nuclei of Substantia Nigra-subregion1 (SN1, compatible with pars reticulata), Substantia Nigra-subregion2 (SN2, compatible with pars compacta), (Datta et al., 1991; Lima et al., 2008, 2007; Olszewski and Baxter, 1954), Caudal-Rostral Linear Raphe (CLi-RLi), Periaqueductal gray (PAG);
• meso-pontine reticular formation nuclei of (Moruzzi and Magoun, 1949; Parvizi and Damasio, 2001) mesencephalic reticular formation (mRt), cuneiform nucleus (CnF), isthmic reticular formation (isRt), and pontine reticular formation (PnO-PnC);
• noradrenergic nuclei of locus coeruleus (LC) (Parvizi and Damasio, 2001; Saper et al., 2010, 2001);
• cholinergic nuclei of pedunculotegmental (PTg) and Laterodorsal Tegmental Nucleus-Central Gray of the rhombencephalon (LDTg-CGPn) (Parvizi and Damasio, 2001; Saper et al., 2010, 2001)] and
• motor function nuclei of Red nucleus-subregion1 (RN1), Red Nucleus-subregion2 (RN2), Subcoeruleus nucleus (SubC), Inferior Olivary Nucleus (ION) (Merel et al., 2019).

As targets, we used the 31 seeds as well as other 15 brainstem nuclei labels (12 bilateral and 3 unilateral, for a total of 27 target regions, developed by this group (Bianciardi et al., 2018, 2016; Garcia-Gomar MG et al., 2021; García-Gomar et al., 2019b; Singh et al., 2021, 2019), namely: Superior Colliculus (SC), Inferior Colliculus (IC), Ventral Tegmental Area-Parabrachial Pigmented Nucleus (VTA-PBP), Microcellular Tegmental Nucleus-Prabigeminal nucleus (MiTg-PBG), Lateral Parabrachial Nucleus (LPB), Medial Parabrachial Nucleus (MPB), Vestibular nuclei complex (Ve), Parvocellular Reticular nucleus Alpha (PCRtA), Superior Olivary Complex (SOC), Superior Medullary Reticular formation (sMRt), Viscero-Sensory Motor nuclei complex (VSM), Inferior Medullary Reticular formation (iMRt), Raphe Magnus (RMg), Raphe Obscurus (ROb) and Raphe Pallidus (RPa). Some of the target nuclei were included in arousal and motor network based on animal studies literature viz LPB, MPB (Kaur et al., 2013b), VTA-PBP (Solt et al., 2014), PCRtA, sMRt, iMRt (Pfaff et al., 2012). For both seeds and targets, probabilistic atlas labels were binarized at a 35 % threshold. For an example dataset, in Supplementary Figure 1, we show the alignment of all brainstem seeds (red) on the T_1_–weighted image, 3 Tesla and 7 Tesla pre-processed fMRI. Also, these seeds and targets are shown on MNI template as Supplementary Figure 2. Additionally, we used as targets, 74 FreeSurfer bilateral cortical (Destrieux et al., 2010) and 10 bilateral single-subject subcortical parcellations (described above in 2.2) including cerebellar cortex, caudate, putamen, pallidum, accumbens area, amydgala, hippocampus, thalamus, sub-thalamic subregion1, and 2 (Bianciardi et al., 2015), and the hypothalamus (Pauli et al., 2018), mapped from MNI space to single-subject space, for a total of additional 169 target regions. The total number of target regions was 227.

We also investigated arousal and motor networks as circuits where we defined nodes involved in these two circuits based on existing literature. These circuits have some overlapping nodes however they have been investigated as two separate networks in the current work. Arousal circuit comprised of frontal cortex, thalamus (Redinbaugh et al., 2020), accumbens (Luo et al., 2018), hypothalamus (Szymusiak and McGinty, 2008), MnR, mRt, SN1, SN2, PAG, CLi-RLi, CnF, isRT, PTg, DR, parabrachial nuclei, LC, PnO-PnC, PMnR and VTA-PBP (Solt et al., 2014). The motor circuit included motor cortex, caudate, putamen, pallidum, RN1, RN2, mRt, isRT, CnF, cerebellar cortex, PTg, intermediate reticular formation-iMRt (Pfaff et al., 2012), Superior Medullary Reticular formation-sMRt, ION, Parvocellular Reticular nucleus Alpha-PCRtA, SubC, PnO-PnC, SN1 and SN2 (Middleton and Strick, 2000; C.J. Stoodley and Schmahmann, 2010) (Matelli et al., 2004). This is also shown in Table 1.

##### 2.2b (ii): Region-based correlation analysis

For each subject, we averaged the time-courses within seed and target regions (or “nodes”) and computed the *Pearson correlation coefficient* across them, generating a 31 × 169 correlation matrix (with 31 × (31 − 1)/2 + 31 × (227 − 31) = 6541 correlation values above the diagonal). We applied on the correlation values a Fisher transform to have null mean and unitary standard deviation (Z-value). To test for statistical significance, a two-sided one sample Student’s t-test was performed on Fisher transformed Z-values across subjects with the null hypothesis of zero correlation, followed by Bonferroni correction for multiple comparisons (p = 0.05) across the whole matrix. For each seed region, a connectome of significant Z-values (which we call “connections” or “links”) at the group level was displayed using a circular 2D representation (Matlab v. 8.4, Natick, MA), as used in our earlier studies (Bianciardi et al., 2016) and referred to as ‘2D circular connectome’ in this work.

We provided graph analysis metrics for the full connectome: in particular we employed the GRETNA (GRaph thEoreTical Network Analysis) toolbox (http://www.nitrc.org/projects/gretna/) (Wang et al., 2015) in Matlab to compute global graph measures (assortativity, rich club, synchronization, hierarchy, clustering coefficient, characteristic path length, gamma, lambda, sigma, global efficiency and local efficiency) and, for each seed, nodal ones (betweenness centrality, degree centrality, local efficiency, clustering coefficient, shortest path length, normalized participant coefficient). We ran the graph analysis on the same matrix that we used to build the connectomes, i.e. the matrix of -log10(p-values) obtained from the group-level analysis, thresholded at p < 0.0005 Bonferroni corrected for multiple comparisons. Finally, we computed for each bilateral seed a laterality index in Matlab defined as the difference between binary links of left and right mirrored seeds, divided by the number of active links, scoring thus 0% for perfectly symmetric nuclei and 100% for perfectly asymmetric ones. The degree centrality, reflecting information communication ability of a given node in the functional network, was used to define nodes as hubs.

##### 2.2b (iii): Voxel-based analysis

For voxel-based analysis, we used as regressors the average timecourse within seeds, computed on unsmoothed data. The single-subject regression analysis (AFNI) was then performed on smoothed fMRI data in MNI space with a Gaussian kernel with 1.25mm standard deviation. In the regression step, to mask out noisy regions filled with CSF, we used a binary mask of the MNI template that included only gray and white matter tissues. The mask is distributed with the FSL software. Further, we computed the group-level statistics (two-sided one sample t-test, AFNI) on the first level beta coefficients. For display purposes, a per-voxel p-value between 10^-5^ and 10^-4^ was chosen to threshold the activation maps. The false-discovery rate at cluster level was controlled using a corrected p-value between 0.01 and 0.05 obtained by imposing a minimum cluster-size as computed using AFNI program 3dClustsim from the estimated autocorrelation function of the residuals). These results showed activations at the subcortical level. For cortical activations, the thresholded data was projected on the pial surface using FreeSurfer.

#### 2.2 c: 3 Tesla fMRI Preprocessing

The resting state fMRI data acquired at 3 Tesla were preprocessed using similar pipeline as used for 7 Tesla resting state fMRI data. The only difference in processing pipeline was in the coregistration of two subjects where the coregistration strategy (also used at 7 Tesla) did not give significant quality. In these two subjects only affine coregistration was used because non linear coregistration introduced small yet visible inaccuracies at the level of the anterior pons, most likely due to neighboring large vessels, and the lack of spatial resolution/contrast to discriminate between the two tissue types. Also, 3 Tesla data underwent only region-based analysis, as the goal was to assess translatability by comparing the region-based connectivity matrices.

### 2.3 : Diagram generation

We generated a schematic diagram of the arousal functional circuit and of the motor functional circuit, using as nodes the brainstem nuclei involved in arousal (Saper et al., 2010, 2001) and motor (Merel et al., 2019) function based on human and animal literature. We then used the connectivity values of our human connectome to build a circuit diagram. Specifically, we averaged out the connectivity strength of node subregions to yield a single connectivity value among to nodes and varied the line thickness of each link based on the connectivity strength.

## 3. Results

### 3.1 : Functional connectome of our arousal-motor nuclei

Our results showed arousal-motor functional network connectome spanning major brainstem and brain regions. The connectivity matrix and 2D circular connectome of all arousal and motor nuclei are shown in Figure 1; global graph measures and laterality index of these nuclei are shown in Figure 2. With regard to the global graph measures, the brainstem-to-brainstem connections showed higher small-world measures and efficiency compared to cortical-to-cortical connections, and lower assortativity and hierarchy measures. For all the seeds, the laterality index displayed high (mirrored) symmetry of the connectivity maps of left and right nuclei. Individual left sided nuclei involved in arousal-motor network showed extensive, yet specific connectivity as shown in Figures 3-11 Similar (mirrored) connectomes were seen for right-sided nuclei as well, thus, were not displayed (see also laterality index in Figure 2). The voxel based functional activations are displayed as surface and volume display that match the results obtained in the region-based 2D circular connectomes. In Supplementary Figure 3, we show 3 Tesla connectome of two representative nuclei (PAG, from the arousal system, and RN1, from the motor system).

**Figure 1.**
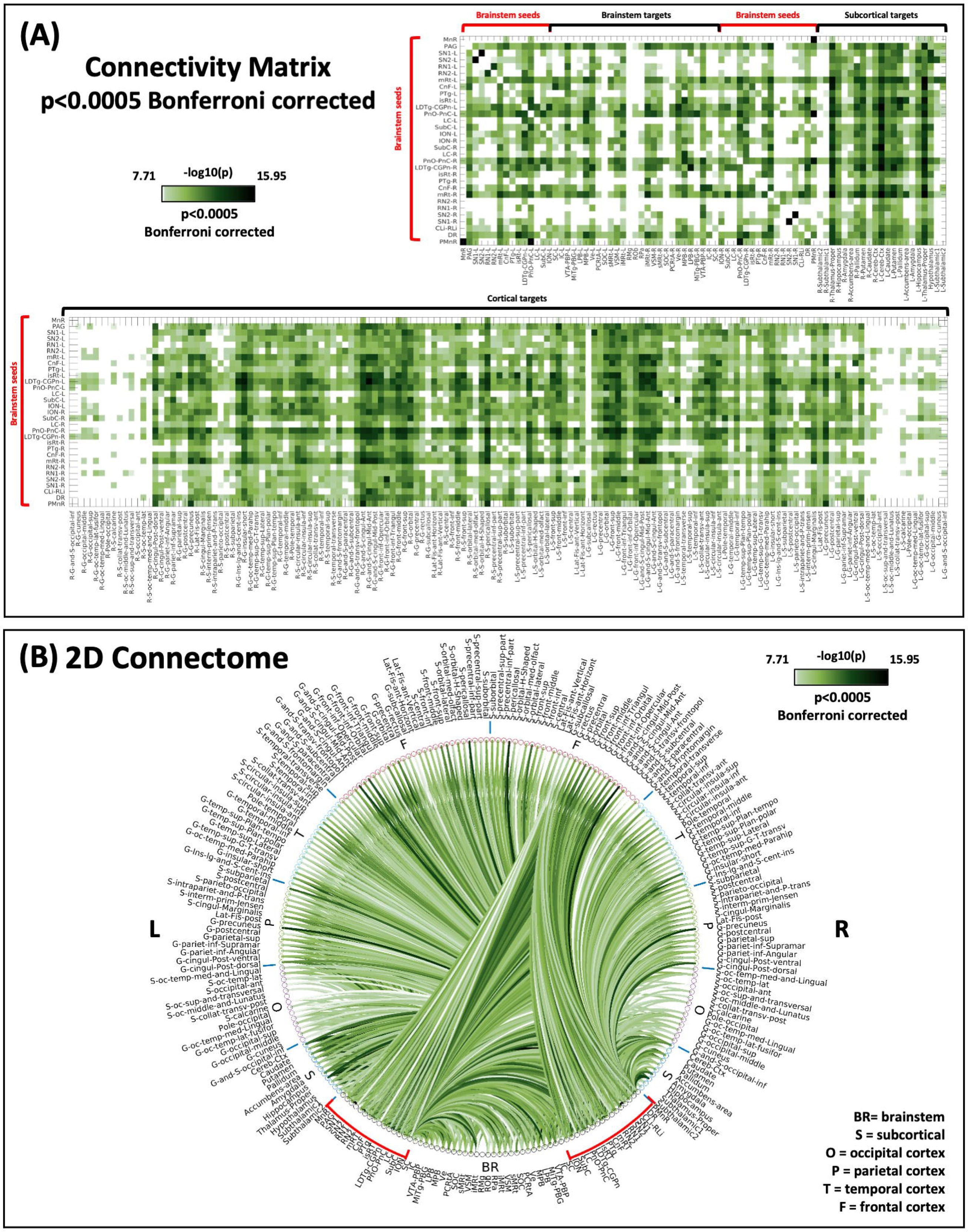
**(A). Connectivity matrix, of all arousal and motor nuclei**. We show the 2D connectivity matrix of the arousal-motor brainstem nuclei at p < 0.0005 Bonferrroni corrected threshold. (**B). 2D circular connectome of all arousal and motor nuclei**. We show the region-based 2D-functional connectome at the group level of the arousal-motor brainstem nuclei at p < 0.0005 Bonferrroni corrected threshold. **List of abbreviations: (brainstem nuclei used as seeds are marked with red brackets:** Median Raphe nucleus (MnR), Periaqueductal Gray (PAG), Substantia Nigra-subregion1 (SN1), Substantia Nigra-subregion2 (SN2), Red nucleus-subregion1 (RN1), Red Nucleus-subregion2 (RN2), Mesencephalic Reticular formation (mRt), Cuneiform (CnF), Pedunculotegmental nuclei (PTg), Isthmic Reticular formation (isRt), Laterodorsal Tegmental Nucleus-Central Gray of the rhombencephalon (LDTg-CGPn), Pontine Reticular Nucleus, Oral Part-Pontine Reticular Nucleus Caudal Part (PnO-PnC), Locus Coeruleus (LC), Subcoeruleus nucleus (SubC), Inferior Olivary Nucleus (ION), Caudal-rostral Linear Raphe (CLi-RLi), Dorsal raphe (DR), and Paramedian Raphe nucleus (PMnR). **Additional brainstem nuclei used as targets:** Superior Colliculus (SC), Inferior Colliculus (IC), Ventral Tegmental Area-Parabrachial Pigmented Nucleus (VTA-PBP), Microcellular Tegmental Nucleus-Prabigeminal nucleus (MiTg-PBG), Lateral Parabrachial Nucleus (LPB), Medial Parabrachial Nucleus (MPB), Vestibular nuclei complex (Ve), Parvocellular Reticular nucleus Alpha (PCRtA), Superior Olivary Complex (SOC), Superior Medullary Reticular formation (sMRt), Viscero-Sensory Motor nuclei complex (VSM), Inferior Medullary Reticular formation (iMRt), Raphe Magnus (RMg), Raphe Obscurus (ROb) and Raphe Pallidus (RPa). **Abbreviations used:** ASS:Assortativity, RC 21, RC 108:, SYN:Synchornization, HIER: Hiererichy, CLUST: clustering cofficient, cPL: Characteristic Path length, gEFF:global efficiency, lEFF: local efficiency.

**Figure 2.**
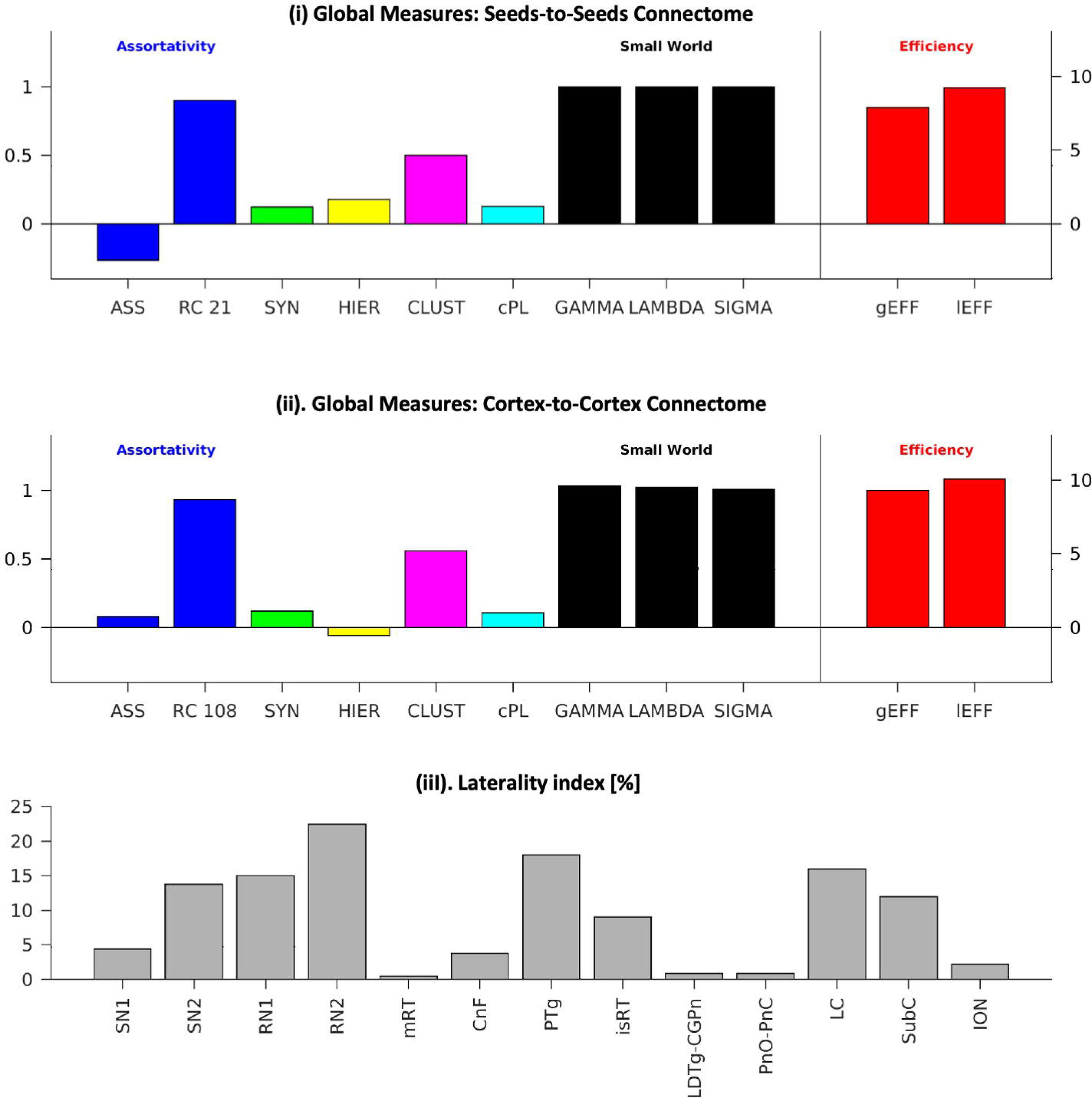
**Global graph measures and laterality index of all arousal and motor nuclei**. We display the global graph measures separately for (i) brainstem to brainstem connections, and for (ii) cortical to cortical connections; in (iii) we display the brainstem nuclei laterality index. **List of brainstem nuclei abbreviations:** Median Raphe nucleus (MnR), Periaqueductal Gray (PAG), Substantia Nigra-subregion1 (SN1), Substantia Nigra-subregion2 (SN2), Red nucleus-subregion1 (RN1), Red Nucleus-subregion2 (RN2), Mesencephalic Reticular formation (mRt), Cuneiform (CnF), Pedunculotegmental nuclei (PTg), Isthmic Reticular formation (isRt), Laterodorsal Tegmental Nucleus-Central Gray of the rhombencephalon (LDTg-CGPn), Pontine Reticular Nucleus, Oral Part-Pontine Reticular Nucleus Caudal Part (PnO-PnC), Locus Coeruleus (LC), Subcoeruleus nucleus (SubC), Inferior Olivary Nucleus (ION), Caudal-rostral Linear Raphe (CLi-RLi), Dorsal raphe (DR), and Paramedian Raphe nucleus (PMnR). **Abbreviations used:** ASS:Assortativity, RC 21, RC 108:, SYN:Synchornization, HIER: Hiererichy, CLUST: clustering cofficient, cPL: Characteristic Path length, gEFF:global efficiency, lEFF: local efficiency.

**Figure 3.**
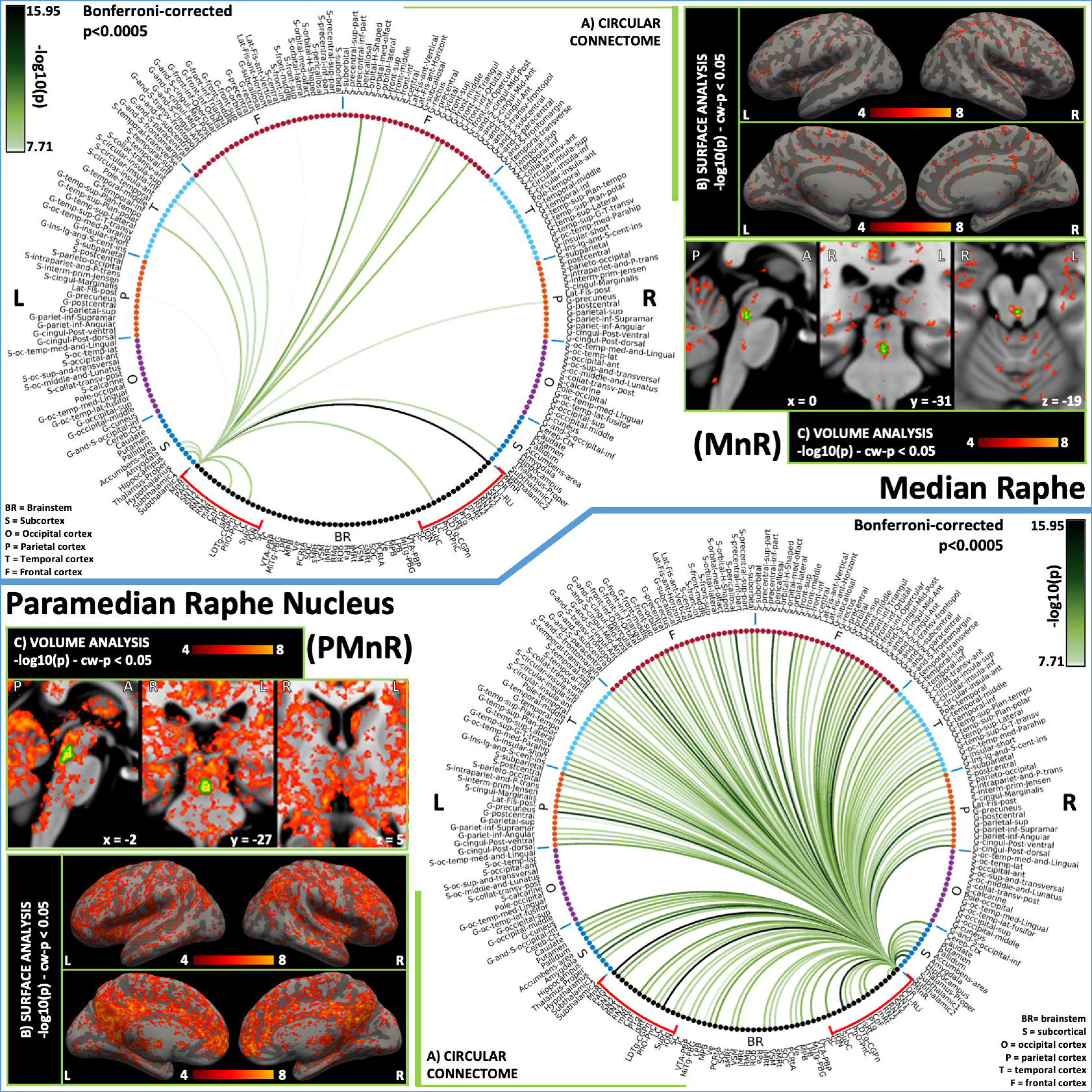
2D circular connectome (A), voxel-based connectivity maps in the cortex (B) and subcortex (C) of (top) MnR and (bottom) PMnR nucleus. MnR showed significant connections with arousal network regions such as hypothalamus, thalamus, along with arousal brainstem nuclei (PMnR, CnF and PnO-PnC). Similarly, PMnR showed significant connectivity with several arousal regions including the cortex, forebrain, hypothalamus and thalamus, along with all major brainstem arousal nuclei (MnR, PAG, SN1, DR, CLi-RLi, mRt, CnF, isRt, VTA, LPB-MPB, LDTG-CGPn, PnO-PnC), except for LC, SN2, PTg. **List of abbreviations: (brainstem nuclei used as seeds are marked with red brackets:** Median Raphe nucleus (MnR), Periaqueductal Gray (PAG), Substantia Nigra-subregion1 (SN1), Substantia Nigra-subregion2 (SN2), Red nucleus-subregion1 (RN1), Red Nucleus-subregion2 (RN2), Mesencephalic Reticular formation (mRt), Cuneiform (CnF), Pedunculotegmental nuclei (PTg), Isthmic Reticular formation (isRt), Laterodorsal Tegmental Nucleus-Central Gray of the rhombencephalon (LDTg-CGPn), Pontine Reticular Nucleus, Oral Part-Pontine Reticular Nucleus Caudal Part (PnO-PnC), Locus Coeruleus (LC), Subcoeruleus nucleus (SubC), Inferior Olivary Nucleus (ION), Caudal-rostral Linear Raphe (CLi-RLi), Dorsal raphe (DR), and Paramedian Raphe nucleus (PMnR). **Additional brainstem nuclei used as targets:** Superior Colliculus (SC), Inferior Colliculus (IC), Ventral Tegmental Area-Parabrachial Pigmented Nucleus (VTA-PBP), Microcellular Tegmental Nucleus-Prabigeminal nucleus (MiTg-PBG), Lateral Parabrachial Nucleus (LPB), Medial Parabrachial Nucleus (MPB), Vestibular nuclei complex (Ve), Parvocellular Reticular nucleus Alpha (PCRtA), Superior Olivary Complex (SOC), Superior Medullary Reticular formation (sMRt), Viscero-Sensory Motor nuclei complex (VSM), Inferior Medullary Reticular formation (iMRt), Raphe Magnus (RMg), Raphe Obscurus (ROb) and Raphe Pallidus (RPa).

**Figure 4.**
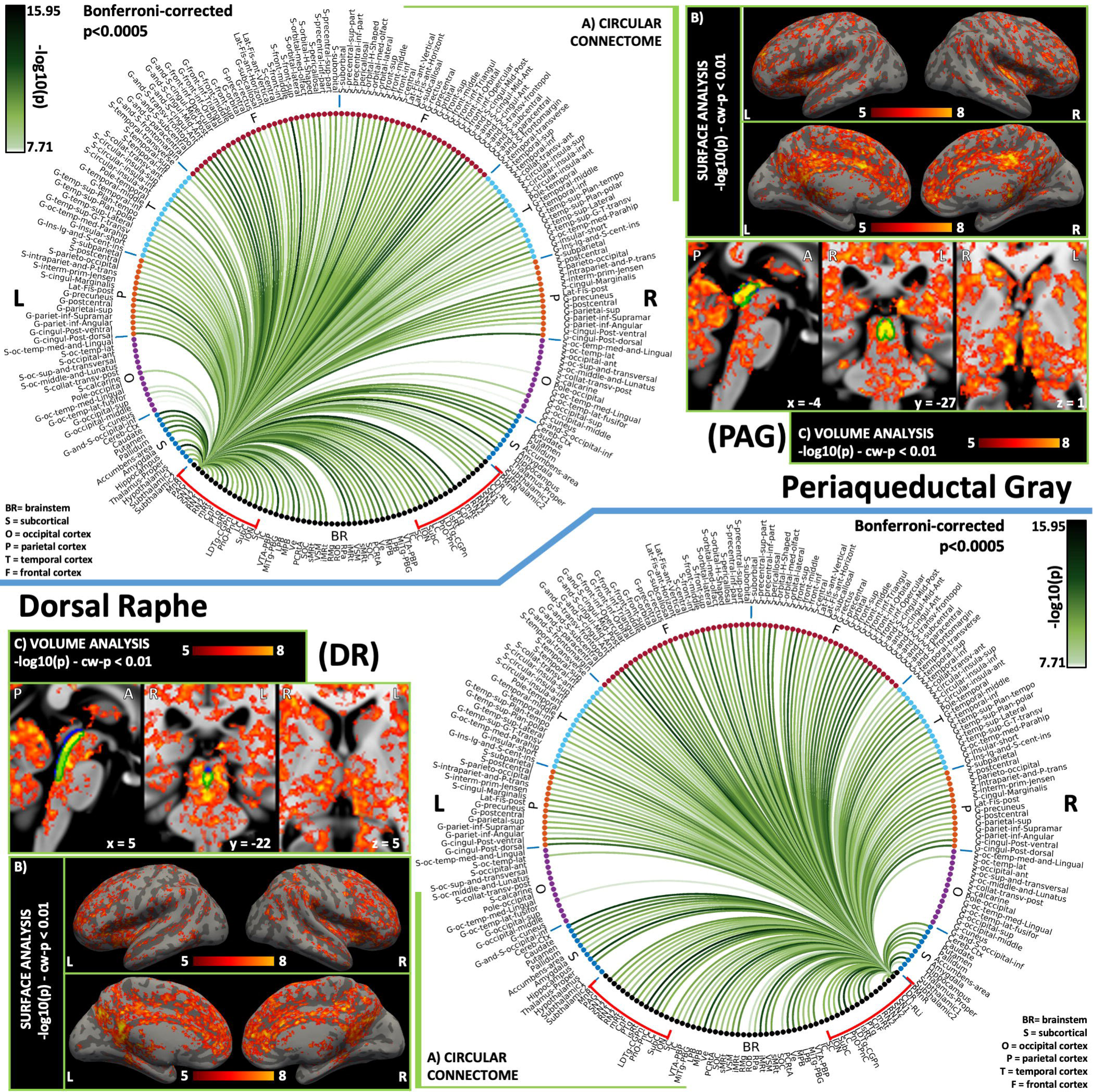
**2D circular connectome (A), voxel based connectivity maps in the cortex (B) and subcortex (C) of (top) PAG and (bottom) DR.** PAG showed strong connectivity with arousal regions (basal forebrain, hypothalamus, thalamus, VTA-PBP, LC, CnF, LDTg-CGPn, and mRt), REM-sleep brainstem nuclei (SubC, sMRt, iMRt), autonomic and limbic regions (hypothalamus, prefrontal cortex, insular cortex, cingulate cortex, LPB, MPB, VSM, and RPa), yet it was not connected with the amygdala. **DR** showed high connectivity with all cortical regions except occipital regions. It showed expected connectivity with other arousal network nuclei except MnR although it showed connectivity with PMnR. **List of abbreviations: (brainstem nuclei used as seeds are marked with red brackets:** Median Raphe nucleus (MnR), Periaqueductal Gray (PAG), Substantia Nigra-subregion1 (SN1), Substantia Nigra-subregion2 (SN2), Red nucleus-subregion1 (RN1), Red Nucleus-subregion2 (RN2), Mesencephalic Reticular formation (mRt), Cuneiform (CnF), Pedunculotegmental nuclei (PTg), Isthmic Reticular formation (isRt), Laterodorsal Tegmental Nucleus-Central Gray of the rhombencephalon (LDTg-CGPn), Pontine Reticular Nucleus, Oral Part-Pontine Reticular Nucleus Caudal Part (PnO-PnC), Locus Coeruleus (LC), Subcoeruleus nucleus (SubC), Inferior Olivary Nucleus (ION), Caudal-rostral Linear Raphe (CLi-RLi), Dorsal raphe (DR), and Paramedian Raphe nucleus (PMnR). **Additional brainstem nuclei used as targets:** Superior Colliculus (SC), Inferior Colliculus (IC), Ventral Tegmental Area-Parabrachial Pigmented Nucleus (VTA-PBP), Microcellular Tegmental Nucleus-Prabigeminal nucleus (MiTg-PBG), Lateral Parabrachial Nucleus (LPB), Medial Parabrachial Nucleus (MPB), Vestibular nuclei complex (Ve), Parvocellular Reticular nucleus Alpha (PCRtA), Superior Olivary Complex (SOC), Superior Medullary Reticular formation (sMRt), Viscero-Sensory Motor nuclei complex (VSM), Inferior Medullary Reticular formation (iMRt), Raphe Magnus (RMg), Raphe Obscurus (ROb) and Raphe Pallidus (RPa).

**Figure 5.**
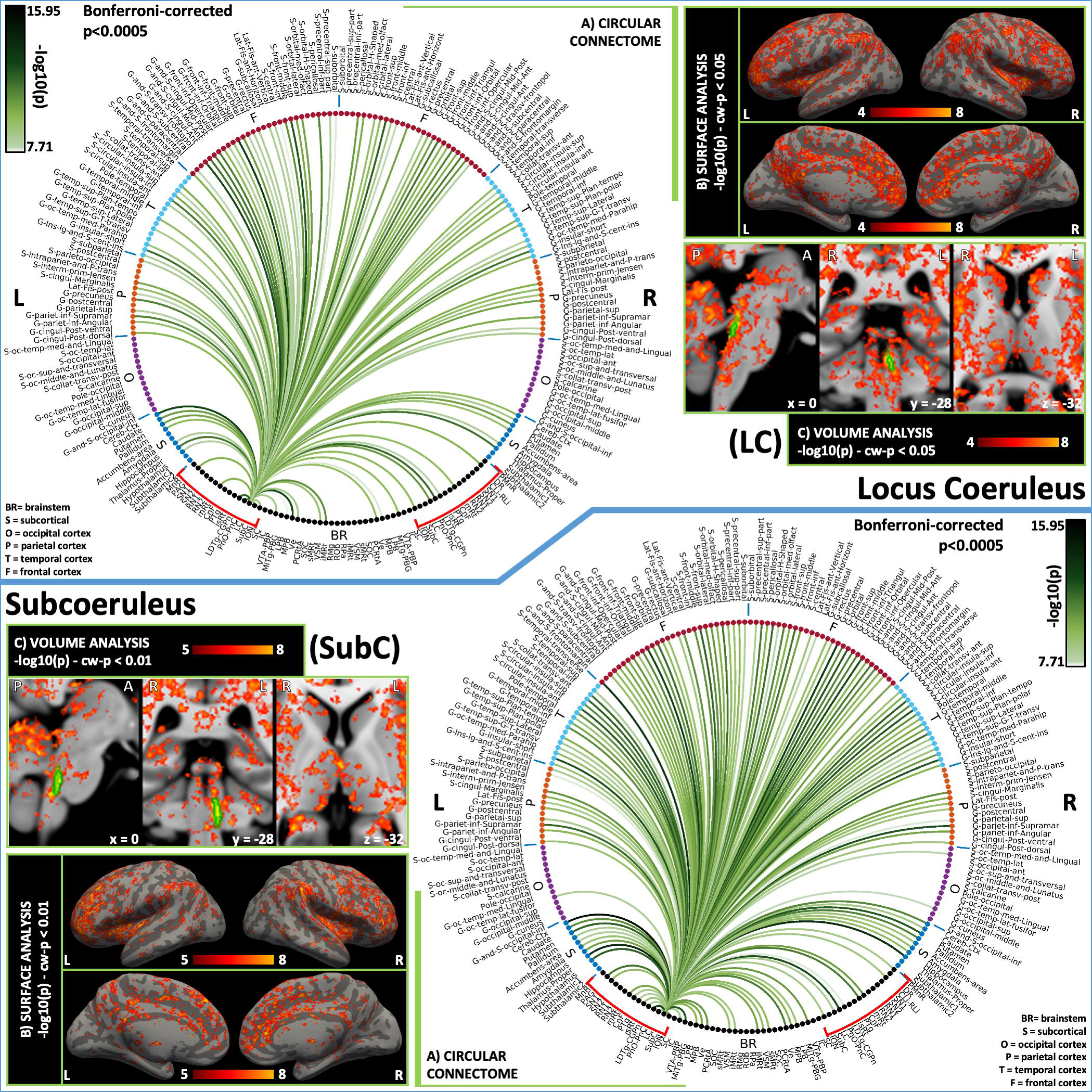
**2D circular connectome (A), voxel based connectivity maps in the cortex (B) and subcortex (C) of (top) LC and (bottom) SubC. LC** showed functional connectivity with the cortex (except occipital regions), thalamus and hypothalamus as expected for an arousal nucleus. Also, it showed connectivity with other arousal-motor network nuclei (PAG, mRt, LDTG-CGPn, PnO-PnC, SubC, ION, DR, VTA-PBP, parabrachial). **SubC**, a motor and REM-sleep nucleus, showed expected connectivity with basal ganglia and cerebellum. It also showed connectivity to brainstem arousal and motor nuclei (mRt, CnF, PTg, isRt, LDTG-CGPn, ION). **List of abbreviations: (brainstem nuclei used as seeds are marked with red brackets:** Median Raphe nucleus (MnR), Periaqueductal Gray (PAG), Substantia Nigra-subregion1 (SN1), Substantia Nigra-subregion2 (SN2), Red nucleus-subregion1 (RN1), Red Nucleus-subregion2 (RN2), Mesencephalic Reticular formation (mRt), Cuneiform (CnF), Pedunculotegmental nuclei (PTg), Isthmic Reticular formation (isRt), Laterodorsal Tegmental Nucleus-Central Gray of the rhombencephalon (LDTg-CGPn), Pontine Reticular Nucleus, Oral Part-Pontine Reticular Nucleus Caudal Part (PnO-PnC), Locus Coeruleus (LC), Subcoeruleus nucleus (SubC), Inferior Olivary Nucleus (ION), Caudal-rostral Linear Raphe (CLi-RLi), Dorsal raphe (DR), and Paramedian Raphe nucleus (PMnR). **Additional brainstem nuclei used as targets:** Superior Colliculus (SC), Inferior Colliculus (IC), Ventral Tegmental Area-Parabrachial Pigmented Nucleus (VTA-PBP), Microcellular Tegmental Nucleus-Prabigeminal nucleus (MiTg-PBG), Lateral Parabrachial Nucleus (LPB), Medial Parabrachial Nucleus (MPB), Vestibular nuclei complex (Ve), Parvocellular Reticular nucleus Alpha (PCRtA), Superior Olivary Complex (SOC), Superior Medullary Reticular formation (sMRt), Viscero-Sensory Motor nuclei complex (VSM), Inferior Medullary Reticular formation (iMRt), Raphe Magnus (RMg), Raphe Obscurus (ROb) and Raphe Pallidus (RPa).

From the GRETNA graph analysis (Figure 12), we found that the betweenness centrality, which is a measure of effect of node on information flow between other nodes, for all nuclei was lower than the network average (35.75) except for PMnR, which had a value of 82. The degree centrality, reflecting information communication ability of a given node in the functional network, was higher (than average) for PAG, RN1, RN2, isRt, LDTg-CGPn and DR, which can then be defined as hubs. Most of the network nuclei showed above average (9.85) local efficiency (a measure of efficient communication of a node among the first neighbors if it is removed), except PAG, mRt and LDTg-CGPn. The clustering coefficient (measuring the likelihood of a node’s neighborhoods connectivity with each other) showed a similar pattern as local efficiency with only PAG, mRt and LDTg-CGPn below network group average (0.54). Except for PAG, mRt, isRt, LDTg-CGPn and DR nuclei, every other nucleus in the network showed above average shortest path length (0.12), indicative of its routing efficiency between itself and all the other nodes in the network. PAG, mRt, isRt, LDTg-CGPn, DR and PMnR showed above average normalized participant coefficient (0.93).

### 3.2 : 7 Tesla versus 3 Tesla results

The association between 7 Tesla and 3 Tesla mean connectivity indices (n=20) in whole brain targets (r-value=0.77), brainstem only targets (r-value=0.76) and cortical/subcortical (other than brainstem) targets (r-value=0.76) was high (Figure 13A). The % of common links between 7 Tesla and 3 Tesla data decreased with increasing the statistical threshold (Figure 13B); for a p-value=0.05 Bonferroni corrected, it was equal to 90 %, 75% and 92% for whole brain targets, brainstem only targets, cortical/other subcortical targets, respectively. These results show translatability of 7 Tesla results in a conventional dataset acquired at 3 Tesla. In Supplementary Figure 3, we also show 3 Tesla versus 7 Tesla connectome of two representative nuclei (PAG, from the arousal system, and RN1, from the motor system). The head motion metrics for 7 Tesla and 3 Tesla data is shown in Supplementary figure 4.

### 3.3 : Circuit Diagram generation

The resulting 2D connectome was used to build a circuit diagram (Figure 14, for 7 Tesla fMRI, and Supplementary Figure 5 for 3 Tesla fMRI) with accompanying connection strengths, thus analyzing the arousal and motor network in literature versus our results. Specifically, we generated a schematic diagram of the arousal and motor functional circuit, using as nodes the brainstem nuclei and cortical regions involved in each function based on human and animal literature (Balaban, 2004; de Lacalle and Saper, 2000; Indovina et al., 2020; Saper and Stornetta, 2015; Yasui et al., 1989), and as links the connectivity values of our human connectome. We averaged out left and right values, and the connectivity strength of subregions belonging to a same node, to yield a single connectivity value among nodes. We varied the line thickness of each link based on the connectivity significance.

## 4. Discussion

In the following section, we present the human functional connectome of arousal and motor nuclei based on 7 Tesla resting-state functional MRI in the context of previous animal and human studies. Specifically, we discuss results of left sided brainstem nuclei shown in Figures 3 to 11. We then summarize the properties of brainstem-brainstem and brainstem-brain human functional connectivity, emerging from basic graph analysis connectivity metrics. Further, we discuss the translatability of the 7 Tesla resting-state fMRI brainstem connectome approach to conventional imaging. Finally, we acknowledge the limitations of this study, and we emphasize the potential of the developed connectome as a preliminary connectivity profile, which might prove useful for future clinical and research work.

### Literature versus our connectome results

#### Median raphe (MnR) and paramedian raphe (PMnR)

The **MnR and PMnR,** defined as a single nuclei complex by some (Behzadi et al., 1990) and as separate nuclei by others (Olszewski and Baxter, 1954; Paxinos G et al., 2012), have been separately delineated (Bianciardi et al., 2015) and their functional connectome detangled in the present study. **MnR** is an arousal nucleus connected to the basal forebrain, VTA, hippocampus, prefrontal cortex, cerebellum, basal ganglia, amygdala, SN, RPa, ROb, nucleus of the solitary tract (VSM), prepositus nucleus (VSM) and interpeduncular nucleus (VTA) (Nieuwenhuys et al., 2008a; Olszewski and Baxter, 1954). Little is known about PMnR connectivity (Olszewski and Baxter, 1954). In this study, we observed sparser fMRI connectivity of MnR (to hippocampus, thalamus, hypothalamus, CnF, PnO-PnC, PMnR, frontal and temporal cortices) as compared to previous literature (Arnsten and Goldman-Rakic, 1984; Behzadi et al., 1990), possibly due to the small size of this nucleus (< 15 mm^3^). Yet, **PMnR** (a ring-shaped label surrounding MnR) showed denser fMRI connectivity pattern, which was in line with the MnR literature (see above). For instance, it showed strong fMRI connectivity with basal forebrain (accumbens and pallidum in our 2D connectomes), hippocampus, basal ganglia, amygdala, as well as with arousal regions such as thalamus, hypothalamus, MnR and PnO-PnC.

#### Periaqueductal gray (PAG) and dorsal raphe (DR)

The **PAG** is a neuromodulatory nucleus involved in arousal, sleep, pain and autonomic function. It receives major afferents from hypothalamus, prefrontal cortex, insular cortex, cingulate cortex, SC, IC (Amunts et al., 2005; Holstege, 1989), which displayed strong functional connectivity in our study. It also showed fMRI connectivity with thalamus, hypothalamus, basal forebrain, VTA-PBP, LC, CnF, LDTg-CGPn, and mRt as expected based on its involvement in arousal and sleep (Saper et al., 2010). The PAG is also a REM-sleep area (Saper et al., 2005; Sapin et al., 2009; Weber et al., 2018), which interestingly showed fMRI connectivity with REM-on areas (SubC) and REM-sleep muscle atonia areas (iMRt and sMRt). Moreover, PAG showed fMRI connectivity with autonomic and limbic nuclei (LPB, MPB, VSM, and RPa), yet it was not connected with the amygdala.

The **DR** is a nucleus involved in sleep and arousal; it has been shown to be connected via efferents to SN2 (compatible with SN pars compacta), caudate, putamen, hypothalamus and thalamus, amygdala, hippocampus, interpeduncular nucleus (IPN-part of VTA-PBP of our nuclei) (Olszewski and Baxter, 1954), LC and parabrachial nuclei (Bobillier et al., 1976), which were present in our connectome as well. DR was also functionally connected to other raphe nuclei, limbic forebrain areas, VTA, PAG, LC, nucleus prepositus (part of VSM nuclei complex in our targets) in line with reports from literature (Jacobs and Azmitia, 1992; Kalén et al., 1985; Poller et al., 2011), yet it lacked connectivity with RMg previously described in rats (Kalén et al., 1985).

#### Locus coeruleus (LC) and Sub-coeruleus (SubC)

The **LC** is a noradrenergic neuromodulator involved in arousal along with vigilance, cognitive processes such as attention, memory and motivation (Sara, 2009). Based on animal studies, LC connects to entire neo-cortex/medial frontal cortex, olfactory bulb, thalamus, amygdala, hippocampus, pallidum, spinal trigeminal nucleus, cochlear nuclei, tectum, cerebellum, spinal cord, PAG, the lateral paragigantocellular nucleus (PGiL), the prepositus nucleus (VSM) (Aston-Jones et al., 1991), DR, solitary nucleus (included in VSM) and ION. With respect to the brainstem, we found similar connections, except for PGiL and the olfactory bulb (not examined as targets in the present study), thus validating our results. We did not find connections of LC to amygdala, which can be further investigated. Regarding the cortical connectivity our results are in line with previous literature reporting widespread connectivity of cortex including frontal, limbic and sensory regions.

The **SubC** is a motor nucleus involved in muscle atonia during REM sleep (Simon et al., 2012) via its connections to gigantocellular nucleus (included in sMRt). Lesions of the SubC lead to the loss of muscle atonia during REM sleep both experimentally in rats, and in humans (Boeve et al., 2007; Karlsson et al., 2005; Xi and Luning, 2009). Crucially, SubC displayed fMRI connectivity to both sMRt and iMRt. Moreover, we found connectivity to motor regions, such as caudate, putamen, pallidum and cerebellar cortex. Further literature suggests SubC is mainly connected to PTg and PnO-PnC (Simon et al., 2012) as can be seen in our connectome. Additional brainstem areas have been reported to have connectivity with SubC, such as PAG, PTg, parabrachial nuclei and Ve (Datta et al., 1998; Olszewski and Baxter, 1954) in line with our current findings. Previous studies have provided anatomical evidence for the ponto-geniculo-occipital wave generating cells involved in the generation of REM sleep from the SubC projecting to occipital cortex (Datta et al., 1998). Our results showed sparse connectivity between SubC and occipital cortex in line with the aforementioned studies (Datta et al., 1998).

#### Laterodorsal tegmental nucleus-central gray of the rhombencephalon (LDTg-CGPn) and pontine reticular nucleus oral part - pontine reticular nucleus caudal cart (PnO-PnC)

The **LDTg-CGPn** is an arousal nucleus involved in modulation of limbic structures (Dobbs and Mark, 2012; Mark et al., 2011). It showed functional connectivity with major brainstem nuclei, cortical and sub-cortical regions involved in arousal. Notably, in line with the literature, it showed functional connectivity with hypothalamus, basal forebrain, VTA (Omelchenko and Sesack, 2005), interpeduncular nuclei (part of VTA in our nuclei delineation), thalamus, medial prefrontal cortex (Satoh and Fibiger, 1986), cingulate and subicular cortices (part of the hippocampus) (Woolf and Butcher, 1986). Previously, connectivity of LDTg to SN has been reported in rats, with stronger connectivity for tracers centered in SN compacta (compatible with SN2 in this study) compared to injections restricted to SN reticulata (compatible with SN1 in this study) (Gould et al., 1989). These findings are in line with our results reporting connectivity of LDTg-CGPn towards SN1 and SN2. Recent studies have shown a direct projection from LDTg to the accumbens (Coimbra et al., 2019), which plays an important role for motivation and positive reinforcement. In line with these findings, in our connectome we observed functional connectivity between LDTg-CGPn and bilateral accumbens. Recent studies have found projections from LDTg to the ventral pallidum, external globus pallidus, amygdala, IC and DR (Dautan et al., 2016), in line with our results as shown in Figure 6. Sparse information exists regarding the connections of CGPn, and further studies are needed with this regard.

**Figure 6.**
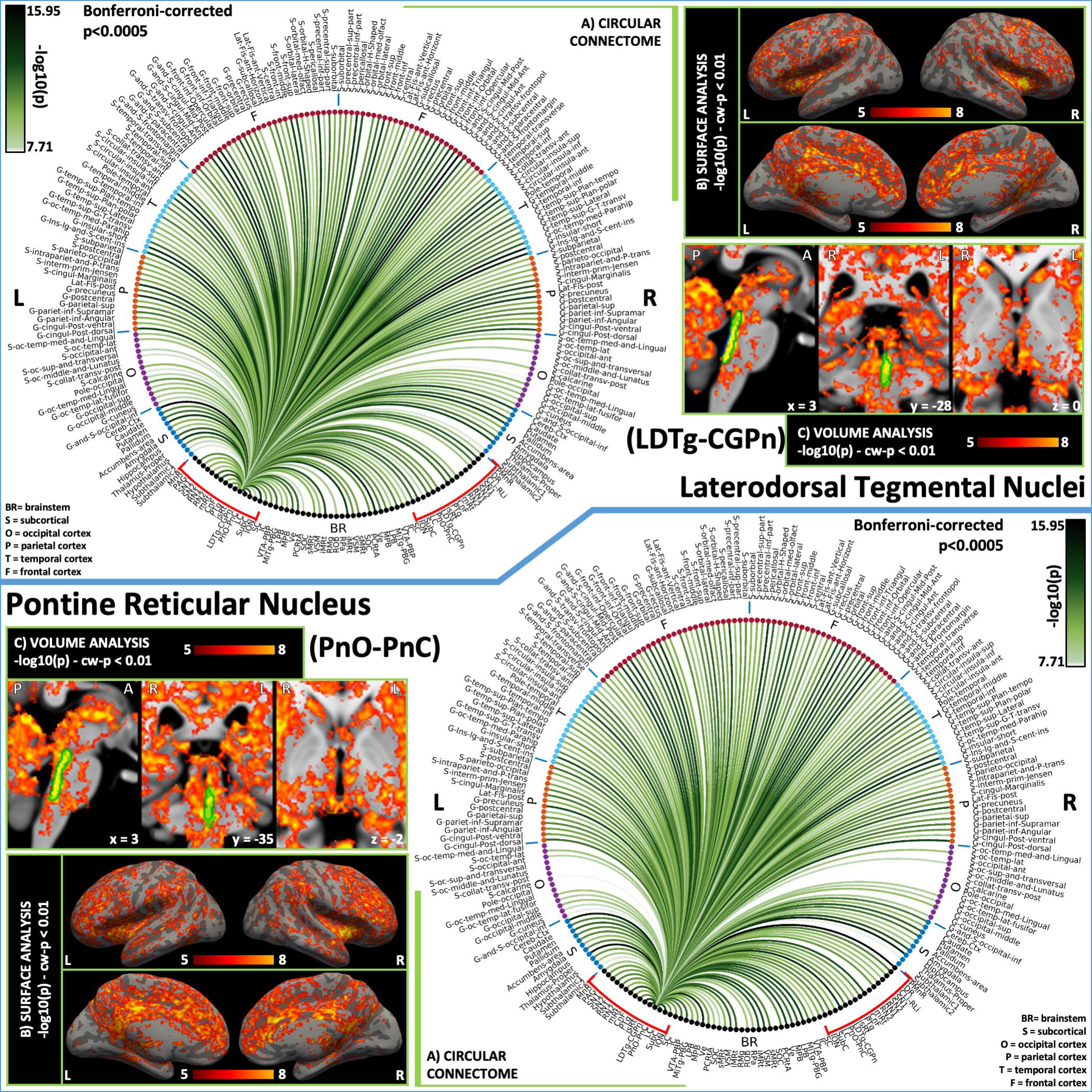
**2D circular connectome (A), voxel based connectivity maps in the cortex (B) and subcortex (C) of (top) LDTg-CGPn and (bottom) PnO-PnC. LDTG-CGPn**, an arousal nucleus, showed widespread functional connectivity with the cortex and subcortex. It showed good connectivity with arousal brainstem nuclei such as MnR, mRt, and arousal regions (hypothalamus, thalamus). **PnO-PnC**, an arousal and motor nucleus, showed strong connectivity with thalamus, hypothalamus, basal forebrain and cortex along with other brainstem nuclei involved in arousal (e.g. raphe nuclei, parabrachial nuclei and VTA-PBP), as well as connectivity with cerebellum, basal ganglia along with motor brainstem nuclei. **List of abbreviations: (brainstem nuclei used as seeds are marked with red brackets:** Median Raphe nucleus (MnR), Periaqueductal Gray (PAG), Substantia Nigra-subregion1 (SN1), Substantia Nigra-subregion2 (SN2), Red nucleus-subregion1 (RN1), Red Nucleus-subregion2 (RN2), Mesencephalic Reticular formation (mRt), Cuneiform (CnF), Pedunculotegmental nuclei (PTg), Isthmic Reticular formation (isRt), Laterodorsal Tegmental Nucleus-Central Gray of the rhombencephalon (LDTg-CGPn), Pontine Reticular Nucleus, Oral Part-Pontine Reticular Nucleus Caudal Part (PnO-PnC), Locus Coeruleus (LC), Subcoeruleus nucleus (SubC), Inferior Olivary Nucleus (ION), Caudal-rostral Linear Raphe (CLi-RLi), Dorsal raphe (DR), and Paramedian Raphe nucleus (PMnR). **Additional brainstem nuclei used as targets:** Superior Colliculus (SC), Inferior Colliculus (IC), Ventral Tegmental Area-Parabrachial Pigmented Nucleus (VTA-PBP), Microcellular Tegmental Nucleus-Prabigeminal nucleus (MiTg-PBG), Lateral Parabrachial Nucleus (LPB), Medial Parabrachial Nucleus (MPB), Vestibular nuclei complex (Ve), Parvocellular Reticular nucleus Alpha (PCRtA), Superior Olivary Complex (SOC), Superior Medullary Reticular formation (sMRt), Viscero-Sensory Motor nuclei complex (VSM), Inferior Medullary Reticular formation (iMRt), Raphe Magnus (RMg), Raphe Obscurus (ROb) and Raphe Pallidus (RPa).

**Figure 7.**
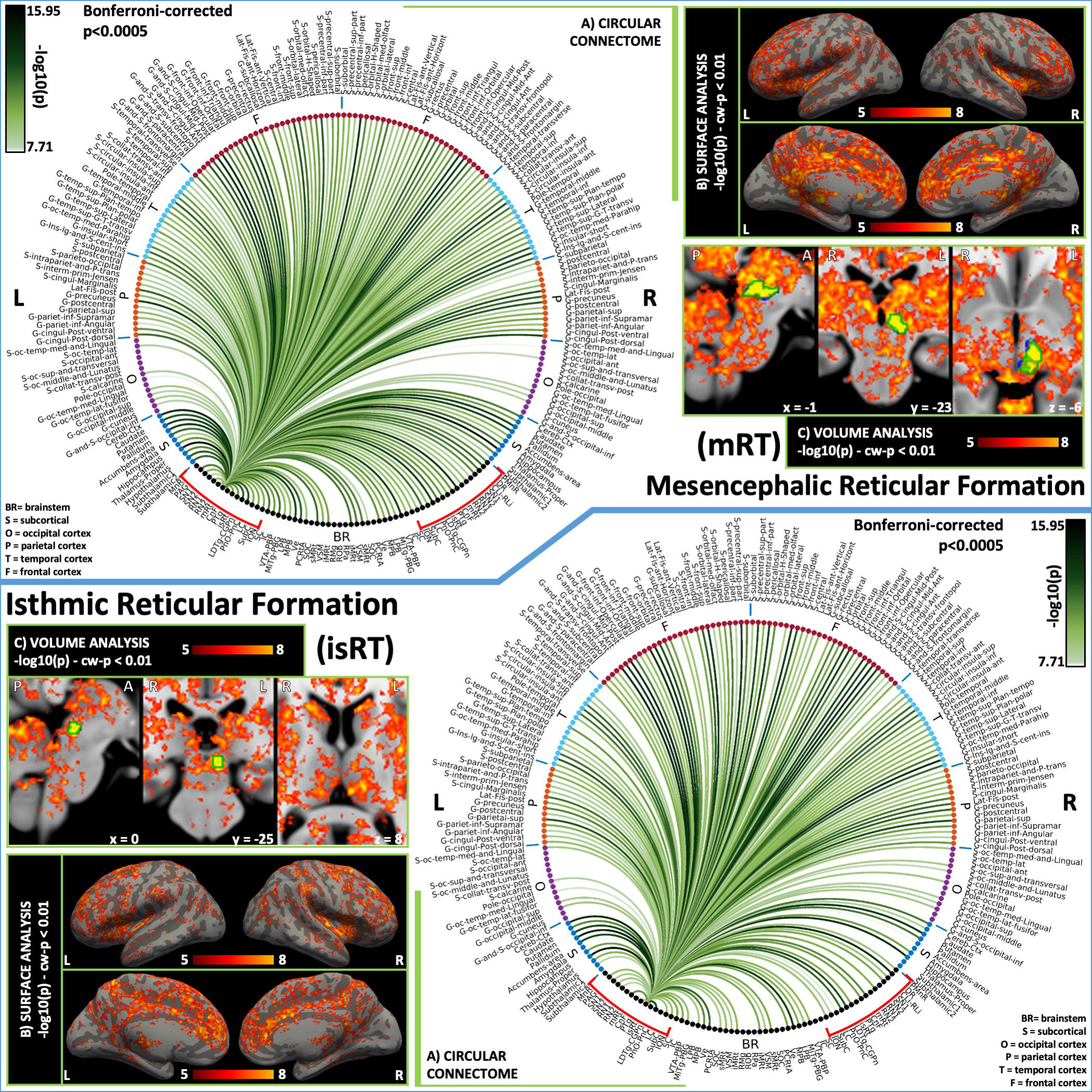
**2D circular connectome (A), voxel based connectivity maps in the cortex (B) and subcortex (C) of (top) mRt and (bottom) isRt (arousal and motor network nuclei).** mRt, and arousal-motor nucleus, showed connectivity with major arousal-motor brainstem nuclei like PAG, LDTg-CGPn, PnO-PnC, parabrachial nuclei, LC, SN1 and SN2 among others. Interestingly, the connectivity towards the frontal eye field (G-frontal-sup, G-frontal-middle, S-frontal-sup and S-frontal-middle) was stronger compared to other cortical areas in line with previous literature (Michael F. Huerta et al., 1986). isRt also showed connectivity with other brainstem nuclei involved in arousal like SN1, SN2, PnO-PnC, DR and parabrachial nuclei. **List of abbreviations: (brainstem nuclei used as seeds are marked with red brackets:** Median Raphe nucleus (MnR), Periaqueductal Gray (PAG), Substantia Nigra-subregion1 (SN1), Substantia Nigra-subregion2 (SN2), Red nucleus-subregion1 (RN1), Red Nucleus-subregion2 (RN2), Mesencephalic Reticular formation (mRt), Cuneiform (CnF), Pedunculotegmental nuclei (PTg), Isthmic Reticular formation (isRt), Laterodorsal Tegmental Nucleus-Central Gray of the rhombencephalon (LDTg-CGPn), Pontine Reticular Nucleus, Oral Part-Pontine Reticular Nucleus Caudal Part (PnO-PnC), Locus Coeruleus (LC), Subcoeruleus nucleus (SubC), Inferior Olivary Nucleus (ION), Caudal-rostral Linear Raphe (CLi-RLi), Dorsal raphe (DR), and Paramedian Raphe nucleus (PMnR). **Additional brainstem nuclei used as targets:** Superior Colliculus (SC), Inferior Colliculus (IC), Ventral Tegmental Area-Parabrachial Pigmented Nucleus (VTA-PBP), Microcellular Tegmental Nucleus-Prabigeminal nucleus (MiTg-PBG), Lateral Parabrachial Nucleus (LPB), Medial Parabrachial Nucleus (MPB), Vestibular nuclei complex (Ve), Parvocellular Reticular nucleus Alpha (PCRtA), Superior Olivary Complex (SOC), Superior Medullary Reticular formation (sMRt), Viscero-Sensory Motor nuclei complex (VSM), Inferior Medullary Reticular formation (iMRt), Raphe Magnus (RMg), Raphe Obscurus (ROb) and Raphe Pallidus (RPa).

**Figure 8.**
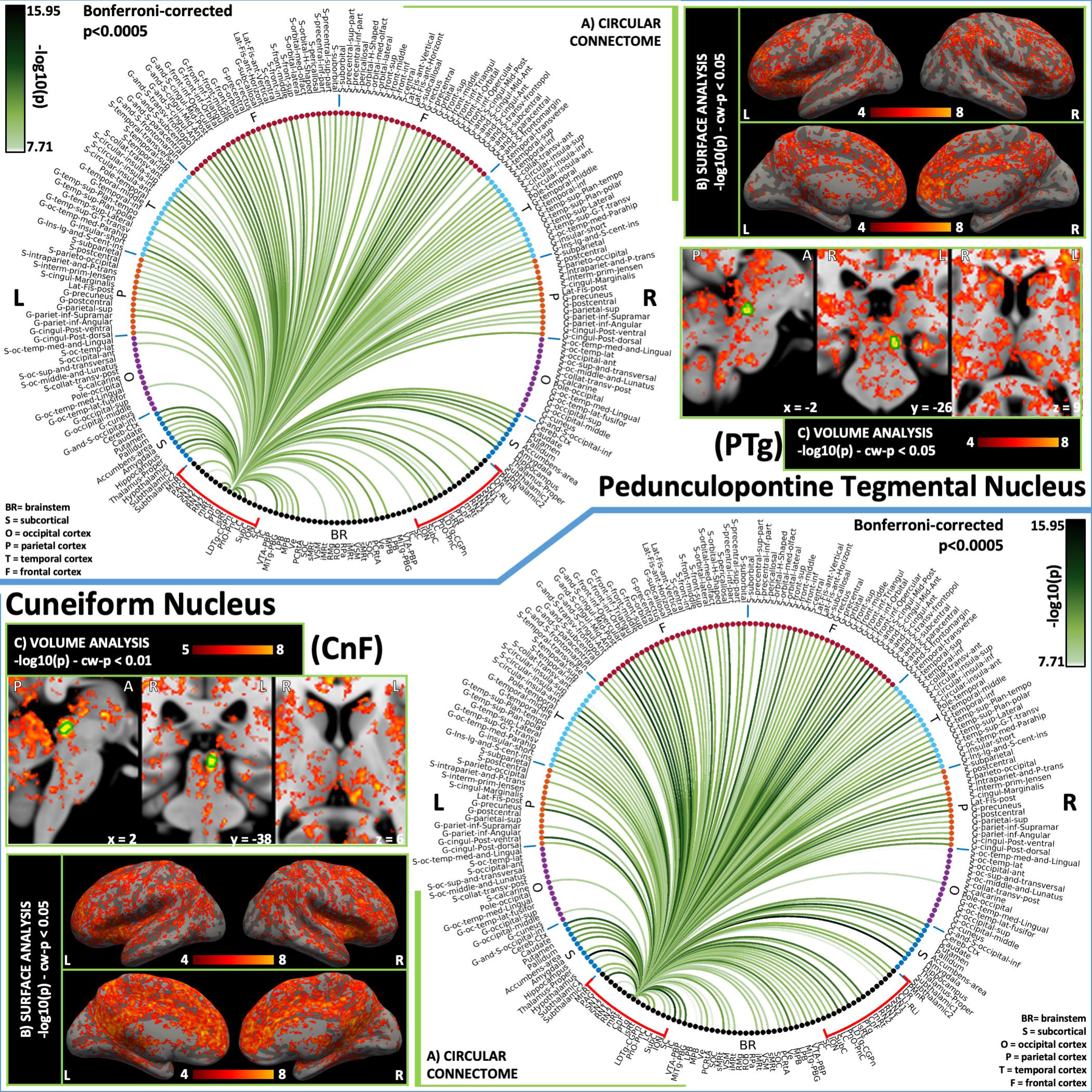
**2D circular connectome (A), voxel based connectivity maps in the cortex (B) and subcortex (C) of (top) PTg and (bottom) CnF, both arousal and motor brainstem nuclei.** PTg showed extensive connectivity to cortex, subcortex, forebrain, basal ganglia and cerebellum as expected for arousal-motor network. Brainstem nuclei showed specific connectivity to CnF, mRt, isRt, LDTG-CGPn, PnO-PnC, LPB, MPB SubC, VTA-PBP, MiTg-PBG, MPB, PAG and DR. CnF similar cortical and subcortical connectivity as PTg. With stronger connectivity with PAG, MiTg-PBG, PMnR and more contralateral brainstem nuclei connectivity compared to PTg. **List of abbreviations: (brainstem nuclei used as seeds are marked with red brackets:** Median Raphe nucleus (MnR), Periaqueductal Gray (PAG), Substantia Nigra-subregion1 (SN1), Substantia Nigra-subregion2 (SN2), Red nucleus-subregion1 (RN1), Red Nucleus-subregion2 (RN2), Mesencephalic Reticular formation (mRt), Cuneiform (CnF), Pedunculotegmental nuclei (PTg), Isthmic Reticular formation (isRt), Laterodorsal Tegmental Nucleus-Central Gray of the rhombencephalon (LDTg-CGPn), Pontine Reticular Nucleus, Oral Part-Pontine Reticular Nucleus Caudal Part (PnO-PnC), Locus Coeruleus (LC), Subcoeruleus nucleus (SubC), Inferior Olivary Nucleus (ION), Caudal-rostral Linear Raphe (CLi-RLi), Dorsal raphe (DR), and Paramedian Raphe nucleus (PMnR). **Additional brainstem nuclei used as targets:** Superior Colliculus (SC), Inferior Colliculus (IC), Ventral Tegmental Area-Parabrachial Pigmented Nucleus (VTA-PBP), Microcellular Tegmental Nucleus-Prabigeminal nucleus (MiTg-PBG), Lateral Parabrachial Nucleus (LPB), Medial Parabrachial Nucleus (MPB), Vestibular nuclei complex (Ve), Parvocellular Reticular nucleus Alpha (PCRtA), Superior Olivary Complex (SOC), Superior Medullary Reticular formation (sMRt), Viscero-Sensory Motor nuclei complex (VSM), Inferior Medullary Reticular formation (iMRt), Raphe Magnus (RMg), Raphe Obscurus (ROb) and Raphe Pallidus (RPa).

**Figure 9.**
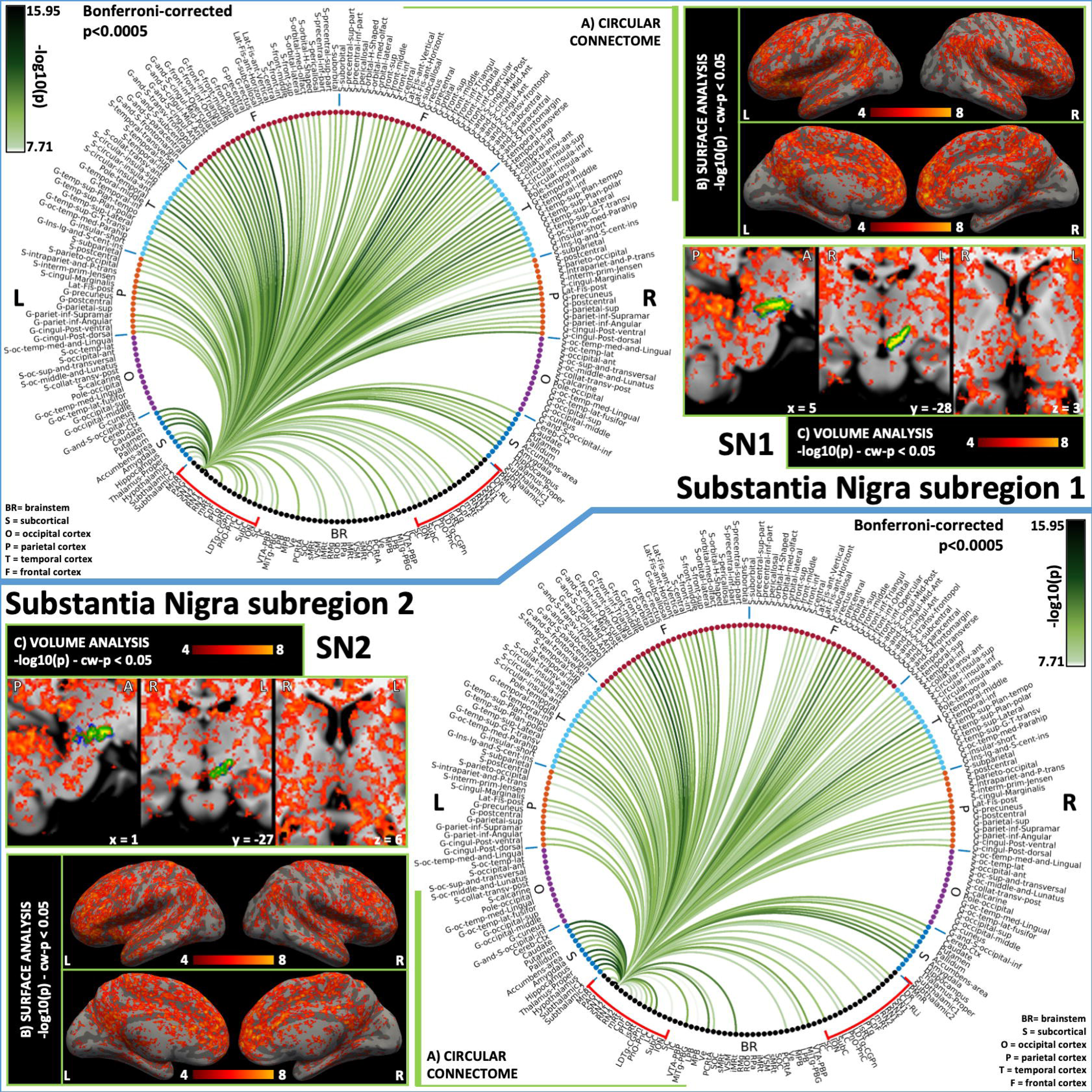
**2D circular connectome (A), voxel based connectivity maps in the cortex (B) and subcortex (C) of (top) SN1 and (bottom) SN2.** The functional connectome of SN1 and SN2 (both arousal/sleep and motor nuclei) displayed similar connectivity pattern except SN2 did not show significant connectivity with PMnR and VSM, and SN1 did not show significant connectivity with MiTg-PBG and PCRtA. Both these regions showed expected connectivity with cortex, thalamus, hypothalamus, forebrain and basal ganglia. **List of abbreviations: (brainstem nuclei used as seeds are marked with red brackets :** Median Raphe nucleus (MnR), Periaqueductal Gray (PAG), Substantia Nigra-subregion1 (SN1), Substantia Nigra-subregion2 (SN2), Red nucleus-subregion1 (RN1), Red Nucleus-subregion2 (RN2), Mesencephalic Reticular formation (mRt), Cuneiform (CnF), Pedunculotegmental nuclei (PTg), Isthmic Reticular formation (isRt), Laterodorsal Tegmental Nucleus-Central Gray of the rhombencephalon (LDTg-CGPn), Pontine Reticular Nucleus, Oral Part-Pontine Reticular Nucleus Caudal Part (PnO-PnC), Locus Coeruleus (LC), Subcoeruleus nucleus (SubC), Inferior Olivary Nucleus (ION), Caudal-rostral Linear Raphe (CLi-RLi), Dorsal raphe (DR), and Paramedian Raphe nucleus (PMnR). **Additional brainstem nuclei used as targets:** Superior Colliculus (SC), Inferior Colliculus (IC), Ventral Tegmental Area-Parabrachial Pigmented Nucleus (VTA-PBP), Microcellular Tegmental Nucleus-Prabigeminal nucleus (MiTg-PBG), Lateral Parabrachial Nucleus (LPB), Medial Parabrachial Nucleus (MPB), Vestibular nuclei complex (Ve), Parvocellular Reticular nucleus Alpha (PCRtA), Superior Olivary Complex (SOC), Superior Medullary Reticular formation (sMRt), Viscero-Sensory Motor nuclei complex (VSM), Inferior Medullary Reticular formation (iMRt), Raphe Magnus (RMg), Raphe Obscurus (ROb) and Raphe Pallidus (RPa).

The **PnO-PnC** is an arousal-motor nucleus complex; as postulated (Olszewski and Baxter, 1954), it showed functional connectivity with the thalamus, hypothalamus, basal forebrain and cortex along with other brainstem nuclei involved in arousal. It also displayed functional connectivity with cerebellum and basal ganglia along with motor brainstem nuclei, in line with (Shinoda et al., 2006). PnO has been associated with the generation of rapid eye movement during REM sleep. Indeed, previous tracers studies in cats have reported connectivity of the PnO with bilateral brainstem structures such as PTg, LC, LDTg and parabrachial nuclei (Rodrigo-Angulo et al., 2005). This is in line with our findings, with the connectome of the PnO-PnC displaying bilateral connectivity to PTg, LC, LDTg-CGPn and parabrachial nuclei (MPB, LPB). PnC has been associated with head and eye movements and previous literature report connectivity with the gigantocellular nucleus (Olszewski and Baxter, 1954), in line with our results showing functional connectivity with sMRt containing the gigantocellular nucleus.

#### Mesencephalic reticular formation (mRt) and Isthmic reticular formation (isRT)

The **mRt** is an arousal-motor nucleus involved in head movements, control of head position (Holstege and Cowie, 1989; Robinson et al., 1994) and eye movements, i.e. gaze (Leigh RJ and Zee DS, 2006). In our connectome, we found expected connectivity of mRt to the frontal eye fields (with a comparable location to G-frontal-sup, G-frontal-middle, S-frontal-sup and S-frontal-middle in our connectomes) (Michael F. Huerta et al., 1986; G. B. Stanton et al., 1988) and the supplementary eye fields (with a comparable location to S-precentral-sup-part and G-and-S-paracentral in our connectomes) (Huerta and Kaas, 1990; B. L. Shook et al., 1990), basal forebrain (Watson et al., 1974), thalamus, hypothalamus, and amygdala. We also found expected connectivity with SC and PAG (May, 2006a), medial pontine and medullary reticular formation, nucleus raphe interpositus, nucleus reticularis tegmenti pontis, LC, and ION (Holstege, 1988; Yezierski, 1988). However, we did not find connectivity with RMg.

The **isRt**, is a relatively understudied nucleus introduced in (Paxinos G et al., 2012) in a region previously pertaining to PT. Based on this location, it has been postulated in the current study to be involved in arousal and motor function. Interestingly, in line with this assumption, we found connectivity of isRt to thalamus, hypothalamus, forebrain and cortex. It also showed strong connectivity to basal ganglia, cerebellum and other arousal and motor nuclei.

#### Pedunculotegmental nucleus (PTg) and Cuneiform nucleus (CnF)

The **PTg** is a nucleus traditionally involved in modulation of REM sleep and motor functions (Olszewski and Baxter, 1954) through its thalamocortical, basal ganglia, nigral (SNr) (Kim et al., 1976; Nauta and Cole, 1978; Parent and De Bellefeuille, 1982) and cerebellar connections (Hazrati and Parent, 1992). Notably, these connections were present in our fMRI-based connectome as well, however connectivity towards SN subregions 1 and 2 was not found bilaterally. PTg has been known to be involved in arousal functions likely due to its ascending projections towards the midbrain, forebrain structures (Erro et al., 1999), and notably in the basal ganglia, which have a direct influence in the striatum and also an indirect influence through the thalamus. Recently it has been proposed that PTg plays an important role while an action becomes obsolete and it needs to be replaced with a new action, similar to the function of subthalamic nucleus at suppressing striatal output (Mena-Segovia and Bolam, 2017). Previously direct connectivity between PTg and subthalamic nucleus has been described, resulting in a positive feed-forward circuit (French and Muthusamy, 2018), nonetheless in our connectomes the link between PTg and subthalamic nucleus is sparse. Ascending projections of PTg also comprises projections to SC and IC that can be associated with modulation of attention shifts and saccadic movements. However, in our results, we lack functional connectivity of the PTg with SC and IC. The descending pathway of PTg, involved in motor inhibition, projects to PnO-PnC and nucleus gigantocellularis (Mena-Segovia and Bolam, 2017), which is in line with our findings (nucleus gigantocellularis being included in sMRt). Dense connections to motor cortex (compatible with G-precentral and S-precentral-sup-part in our connectomes), supplementary motor area (compatible with G-Front-sup in our connectomes) and frontal eye field from PTg have been reported previously (French and Muthusamy, 2018) in line with our current findings. This connectivity profile is in line with the involvement of PTg in movement control, sleep and cognition.

The **CnF** is involved in arousal and motor function (Gert Holstege, 1991) and is considered the ‘mesencephalic locomotion region’ along with PTg. We found functional connectivity of this nucleus with raphe nuclei (which modulate locomotion and nociception (Beitz, 1982)), PAG, and hypothalamus in line with previous literature (Gert Holstege, 1991). We also found bilateral connectivity of CnF with medullary reticular formation (sMRt, iMRt), in line with previous tracing studies in rodents mapping its projection towards the spinal cord (Xiang et al., 2013). In agreement with the literature (Chang et al., 2020), we also found connectivity between CnF and colliculi (SC and IC). Recently, CnF has been postulated to be a more efficacious target than PTg for treating freezing of gait with deep brain stimulation due to its less diffuse brain connectivity compared to PTg (Chang et al., 2020). This is at odds with our findings showing a similar widespread functional connectivity of both nuclei with the rest of the brain. Further investigation is needed regarding this potential target of deep brain stimulation as a treatment option.

#### Substantia nigra (SN) and its sub-divisions

The **SN** subregions (SN1, compatible with pars reticulata, and SN2, compatible with pars compacta) are involved in arousal and motor functions (generation and control of voluntary motor actions) and are largely studied in movement disorders. In line with afferents reported in the literature (Olszewski and Baxter, 1954), SN showed functional connectivity to six main areas, namely striatum, globus pallidus, the subthalamic nucleus, PTg, DR and the cerebral cortex. Based on previous studies PTg targets SN compacta (Olszewski and Baxter, 1954), and interestingly our fMRI results were in line with this connectivity profile. SN2 showed connectivity with ipsilateral PTg, as opposed to SN1, which did not show this connectivity pattern. Further, the observed links of SN to thalamus, PTg, prefrontal and motor cortex were expected based on reported efferent connections (Ilinsky et al., 1985). Finally, we did not find connection to the SC, as expected based on previous work (Ilinsky et al., 1985). Whilst SN1 and SN2 showed a similar connectivity profile they have also some differences, in general SN1 showed more connectivity to contralateral brainstem nuclei.

#### Caudal linear raphe (CLi) and inferior olivary nucleus (ION)

The **CLi**, a nucleus involved in the regulation of arousal, is expected to connect to the basal forebrain (e.g. nucleus accumbens) (Ikemoto, 2007), basal ganglia, amygdala (Imai et al., 1986), hypothalamus (Mai JK and Paxinos G, 2011; Nieuwenhuys et al., 2008b), SNc, VTA, DR and MnR (Jacobs and Azmitia, 1992). The **RLi**, related with reward, motivation and cognition, is connected to PTg, LDTg-CGPn, (Garzón et al., 1999; Oakman et al., 1995), LC, raphe nuclei, hypothalamus, nucleus accumbens and other limbic regions (Mai JK and Paxinos G, 2011). Crucially, our CLi-RLi nuclei complex displayed connectivity to the majority of aforementioned regions except for PTg, LC and MnR (yet, we found connectivity with PMnR, a region surrounding the MnR).

The **ION** has been implicated in motor learning and motor error correction through coordination and refinement of movements; also, it is known to be involved in proprioception with evidence of connectivity to cerebellum (olivocerebellar pathways), Ve, SC, RN, motor and somatosensory cortex (Barmack, 2006; Catherine J. Stoodley and Schmahmann, 2010; Voogd et al., 2013). We found similar functional connectivity in our results along with basal ganglia, augmenting its role in motor function. With regard to olivocerebellar connectivity, our results displayed higher ipsilateral connectivity between ION and ipsilateral cerebellum compared to contralateral cerebellar cortex; we also observed ipsilateral connectivity to SC and bilateral connectivity to Ve possibly to with the optokinetic signal to ION. The ION receives afferents from the prepositus hypoglossi nucleus (Olszewski and Baxter, 2014) that is comprised in the VSM; interestingly we found bilateral connectivity of ION and VSM.

#### Red nucleus (RN) and its sub-divisions

The **RN**, a motor nucleus, showed expected connectivity to motor cortex, supplementary motor cortex, rostral premotor area, the supplementary and frontal eye fields, thalamus, ION and cerebellum (Alberto Cacciola et al., 2019; Nieuwenhuys et al., 2008b). From histological studies, this nucleus has been traditionally divided into two parts: the magnocellular and the parvocellular subnucleus. The rubro-olivary projections are originated in its majority from the parvocellular part and are involved in the rubro-olivary-cerebellar-motor cortex feedback loop (Alberto Cacciola et al., 2019; Olszewski and Baxter, 2014). The atlas labels of the subregions of RN (RN1, RN2) used in the present study were defined based on different multi-contrast MRI properties of this regions (Bianciardi et al., 2015). Interestingly, our functional connectomes displayed different fMRI connectivity profiles for these subregions. Stronger ipsilateral and contralateral connectivity between ION and RN2 was founded, which might suggest that RN2 could be compatible with the parvocellular part of RN. Previous literature also reported that the parvocellular part of RN is more connected to the cerebral cortex compared to the magnocellular part (Alberto Cacciola et al., 2019); both RN1 and RN2 showed dense connectivity with the cortex, yet RN2 displayed a higher number of links. Finally, our connectomes showed that RN1 has bilateral connectivity with the cerebellum; in contrast RN2 showed unilateral connectivity with the cerebellum, while both subregions showed connectivity with PAG.

### Graph measure analysis

The human brain network is considered to have small-world network properties, which indicate that the brain is organized into both local cliques and global integration. Graph measure analysis has been used (Bullmore and Sporns, 2009; Kim et al., 2019; Nakamura et al., 2009) to explore disease induced alteration in topological parameters of the human brain functional connectivity (Li et al., 2018). Global graph analysis of our connectome showed lower brainstem seed-to-seed assortativity than cortex-to-cortex assortativity, indicating higher interconnectivity in the latter. An increase in assortativity leads to lower average hop count and robustness (Murakami et al., 2017). Thus, our results might indicate that brainstem networks might be more resilient in case of pathology, a hypothesis that needs further investigation. Synchronization, hierarchy, clustering coefficient, path length and other small world network metrics were comparable in seed-to-seed and cortex-to-cortex connectome. The cortex-to-cortex connectome showed slightly higher global and local efficiency than the brainstem seed-to-seed connectome indicating efficient information flow. In both arousal and motor circuit, we observed high inter-connectivity of cortical and sub-cortical regions. All nuclei displayed laterality index lower than 23%, with only SN2, RN1, RN2, PTg, LC and SubC getting above 10%: this indicates quite similar mirrored connectomes of left and right nuclei. RN2 showed highest and mRt the lowest laterality index. The nodal graph measures of arousal-motor network nuclei indicated their segregation into two sets of nuclei namely (i) PAG, RN2, mRt, isRt, LDTG-CGPn and DR and (ii) MnR, SN1, SN2, RN1, CnF, PTg, PnO-PnC, LC, SubC, ION, CLi-RLi and PMnR. Based on properties of random network, i.e lower clustering coefficient and a shorter shortest path length (Zhu et al., 2017), the first set of nuclei can be considered to show random network. Former set of nuclei showed above average degree centrality as compared to the other set nuclei in the network. Degree centrality reflects the number of instantaneous functional connections between a region and the rest of the brain within the entire connectivity matrix of the brain; it can assess how much a node communicates with the entire brain and integrates information across functionally segregated brain regions (Guo et al., 2016a, Guo et al., 2016b). Interestingly, nuclei with high degree centrality (PAG, RN2, mRt, isRt, LDTG-CGPn and DR) were also the most connected nuclei in the arousal circuit, with cortical, sub-cortical and other brainstem nodes. These can also be considered as ‘hubs’ in the network. Nodal efficiency, clustering coefficient, shortest path length in set (ii) of nuclei were higher than set (i) nuclei. The regions with high nodal efficiency imply pivotal roles for communications between any pair of regions within the network (Achard and Bullmore, 2007; Gong et al., 2009). Most of the nodes showed high local efficiency (except PAG, mRt, and LDTg-CGPn), indicating arousal-motor network seeds effectively shared information within their immediate local communities (seen as high connectivity with cortical and sub-cortical regions), thereby assisting effective segregated information processing in the network.

### Translatability of resting state fMRI brainstem connectome in conventional imaging

The association between 7 Tesla and 3 Tesla mean connectivity indices was high (r-value > 0.75) and did not differ across groups of regions (all targets, versus brainstem targets, versus cortical and subcortical (non-brainstem) targets). The percentage of common links between the two scanner data decreased with increasing the statistical threshold, with a steeper decrease for brainstem targets compared to cortical targets. In summary, comparison of 3 Tesla to 7 Tesla connectivity results indicated good translation of our brainstem connectome approach to conventional fMRI, especially for cortical and subcortical (non-brainstem) targets and to a lesser extent for brainstem targets. This was expected due to the small size of brainstem nuclei, their deep location in the brain, and partial volume effects with neighboring pulsatile cerebrospinal fluid, associated with a reduced fMRI sensitivity.

The reduced sensitivity for detecting brainstem connectivity at 3 Tesla compared to 7 Tesla was not explained by changes in temporal signal-to-noise ratio (tSNR) across scanners after preprocessing (Supplementary Figure 6). Indeed, the tSNR at 7 Tesla was about 50 % of the tSNR at 3 Tesla in the whole brain, 66 % in the cortex and 44 % in the brainstem (Supplementary Figure 6, Freesurfer masks in MNI space where used for this computation). Note that an about 50 % value agrees with our estimated tSNR ratio between 7 Tesla and 3 Tesla acquisitions, obtained after accounting for the B_0_, bandwidth, voxel volume and number of repetitions (albeit the computed tSNR also includes the coil sensitivity). The increased sensitivity for brainstem connectivity at 7 Tesla might be due to the increased BOLD contrast with the field strength (which counterbalances the tSNR decrease at 7 Tesla, and provides a favorable contrast-to-noise ratio at 7 Tesla compared to 3 Tesla), and the presence of reduced partial volume effects at 7 Tesla compared to 3 Tesla due to an ∼12 times reduced voxel volume employed at 7 Tesla. Partial volume effects are expected to play a bigger role in the average of BOLD signals across small regions, such as tiny brainstem nuclei, than across larger cortical and subcortical areas.

### Arousal and motor network circuit at 3 Tesla and 7 Tesla in view of existing literature

Due to the adopted connectivity metric (Pearson correlation), in the displayed diagrams we cannot easily rule out direct from indirect connectivity; nevertheless, based on literature we discuss, for some nuclei, which links potentially underlie direct (i.e. monosynaptic) or indirect (i.e. polysynaptic) connections. For instance, PAG showed connections to all brainstem nuclei in the network and cortical and sub-cortical regions. Based on literature, some PAG sub-divisions have direct connections to thalamus (for example, the lateral PAG and ventrolateral PAG innervate the medial and intralaminar thalamic nuclei (Krout and Loewy, 2000)), and hypothalamus (rostral dorsolateral PAG densely projects to hypothalamic nuclei related to defensive responses, such as the anterior hypothalamic nucleus, and dorsomedial hypothalamus); conversely, other PAG sub-regions have moderate (caudal dorsolateral PAG to anterior hypothalamic nucleus and dorsomedial hypothalamus; ventrolateral PAG to dorsomedial hypothalamus) or weak (caudal ventrolateral PAG to dorsomedial hypothalamus) projections (Cameron et al., 1995)) to the hypothalamus. The PAG also receives direct inputs from the primary auditory cortex, secondary visual cortex (Dampney, 2018), and medial prefrontal cortex (mPFC) (processing ascending pain inputs and modulating endogenous analgesia via direct projections to the PAG (Cheriyan and Sheets, 2018). Nevertheless, the observed connectivity of the PAG with the frontal cortex might also include indirect connectivity through the thalamus. Further, the PAG sends direct projections to the DR (Vertes, 1991) parabrachial nuclei (Krout et al., 1998), LC (Bajic and Proudfit, 1999) and mRt. It also showed connectivity with other brainstem nuclei of PnO-PnC (Mantyh, 1983), VTA-PBP (inhibition of the ventrolateral PAG-VTA circuit relieves the aversive state elicited during headache (Waung et al., 2019)), SN (which might be indirect connectivity as shown in rat model of Parkinson’s disease, where SN connects to retrotrapezoid nucleus via PAG (Lima et al., 2018)). Finally, PAG showed connectivity with LDTg-CGPn, which might be an indirect connection via VTA (Coimbra et al., 2017).

In line with expected direct connectivity reported in the literature, mRt showed connectivity to frontal cortex (frontal and supplementary eye field) (M.F. Huerta et al., 1986; B.L. Shook et al., 1990; G.B. Stanton et al., 1988), thalamus, hypothalamus (Olszewski and Baxter, 1954). It showed connectivity with all brainstem nuclei except MnR and SN2 in arousal network. Some of these connections are evidenced to be direct viz PAG (May, 2006b), imRt, smRt (Edwards and de Olmos, 1976), LC, ION (Olszewski and Baxter, 1954), while others might be indirect. It also showed connectivity to cerebellum in motor circuit which might be part of cerebellothalamic connections (Nieuwenhuys et al., 2008a).

CnF showed connectivity with the thalamus, hypothalamus, PAG, as expected based on direct links to these regions from previous work (G. Holstege, 1991), as well as with raphe nuclei (Noga et al., 2017; Steeves and Jordan, 1980). It showed connectivity with other brainstem nuclei of LDTg-CGPn, PnO-PnC, parabrachial, VTA-PBP which might be indicative of indirect connections.

PTg showed connectivity to basal ganglia, cerebellum and motor cortex. PTg directly connects to the basal ganglia and cerebellum, and also acts as motor and cognitive interface between the two, where the cerebellum is connected to the basal ganglia via the PTg (Mori et al., 2016). The PTg connects to the thalamus directly and also act as relay to the cerebellar cortex for basal ganglia (Mori et al., 2016).

Substantia nigra showed connectivity to striatum, which is expected to be monosynaptic (Zhang et al., 2017). It also showed connectivity to prefrontal and motor cortices, which are in line with evidence of direct cortico-nigral pathway in cats (Afifi et al., 1974), rodents (Höglinger et al., 2015), primates (Sakai, 1988) and humans (Cacciola et al., 2016). It showed connectivity to thalamus which might be its direct connectivity as part of projecting inhibitory pathways and part of “corticostriatal-basal ganglia/SN-thalamic-motor cortex loop”(Olszewski and Baxter, 1954).

Ventral tegmental area (VTA-PBP) showed connectivity to frontal cortex, hypothalamus, accumbens area and thalamus as expected and might be considered monosynaptic in nature (Beckstead et al., 1979). It also showed connectivity to DR, LPB, MPB and LC which are also its direct connections based on tracing study in rats (Beckstead et al., 1979).

The RN was functionally connected to motor cortex, cerebellum, thalamus and basal ganglia owing to its role in motor coordination (Olszewski and Baxter, 1954). Interposed nuclei of the deep cerebellum have direct projections to the RN (magnocellular part). RN (parvocellular) receives input from the deep white matter of the cerebellum and redirects input into the ION (A. Cacciola et al., 2019). The RN (parvocellular) is shown to provide important connection between the motor cortex and the cerebellum, and receives inputs directly into the inferior olivary nucleus (Ulfig and Chan, 2001), thus, RN showed direct connections to these regions. It also showed connectivity to PnO-PnC and mRt which might be indirect in nature.

PnO-PnC showed connectivity to motor cortex, cerebellum, basal ganglia and all brainstem regions. Based on literature, its connections to cerebellum, hypothalamus, SN, CnF, parabrachial, DR, LDTg-CGPn and LC can be considered direct in nature (Shammah-Lagnado et al., 1987).

mRt showed connectivity with motor cortex (role in somatic motor system (Olszewski and Baxter, 1954), cerebellum (indirect connection via pontine reticular nuclei (Strassman et al., 1986)) and basal ganglia. It showed connectivity with CnF (Korte et al., 1992), which might be direct connections. It also showed connectivity to PTg, PnO-PnC, SubC, ION, sMRt and iMRt which can be indirect connections.

ION showed connectivity to the motor cortex which could include both indirect connection via the cerebellum as well as direct connection (cortico-olivo-cerebellar projections), as investigated in cats (Saint-Cyr, 1983).

Arousal and motor circuit diagrams at 7 Tesla (Figure 14) and 3 Tesla (Supplementary Figure 5) showed similar connectivity with cortical and sub-cortical regions, yet with some differences mainly in the brainstem, as also visible from their connectivity matrices (Supplementary Figure 7). Noticeably, the 7 Tesla dataset showed greater connectivity in the brainstem than the 3 Tesla dataset. Moreover, the connectivity of PAG to thalamus was present in both 3 Tesla and 7 Tesla datasets, however, the connectivity to the hypothalamus was found in the 3 Tesla dataset only. Both 3 Tesla and 7 Tesla results showed PAG connectivity to frontal cortex. However, the expected PAG direct projections to the parabrachial nuclei (Krout et al., 1998) and LC (Bajic and Proudfit, 1999) were found at 7 Tesla but not at 3 Tesla. DR and mRt showed connectivity with PAG at both 3 Tesla and 7 Tesla. mRt showed connectivity with all brainstem nuclei except MnR and SN2 at 7 Tesla and with CnF, PnO-PnC, LPB and VTA-PBP at 3 Tesla, within the arousal network. RN showed connectivity to PnO-PnC and mRt (both at 3 Tesla and 7 Tesla) which might be indirect in nature. Thalamus connected to all brainstem nuclei except MnR at 7 Tesla and also PMnR at 3 Tesla. Striatum connected to all brainstem nuclei both at 7 Tesla and 3 Tesla.

**Figure 10.**
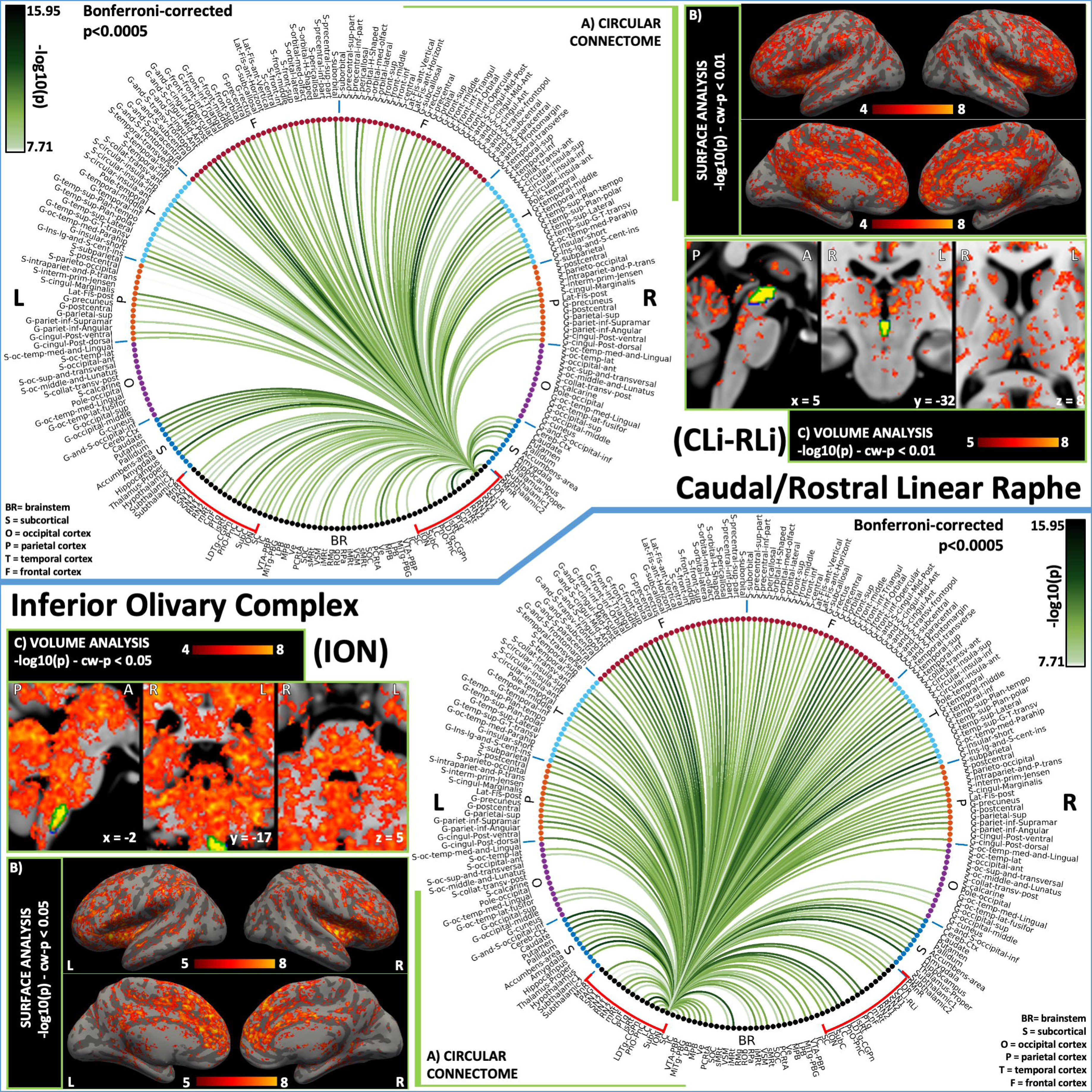
**2D circular connectome (A), voxel based connectivity maps in the cortex (B) and subcortex (C) of (top) CLi-RLi and (bottom) ION.** CLi-RLi, an arousal nucleus, showed expected connectivity with regions of frontal, temporal and parietal lobes [based on(Ikemoto, 2007)]. Subcortex also showed extensive connectivity as expected. It showed connectivity with arousal regions either ipsilaterally or contralaterally like mRt, LDTg-CGPn, PnO-PnC, thalamus and hypothalamus. ION, a motor brainstem nucleus, showed connectivity with motor cortical areas, cerebellar cortex and ipsilateral RN subregions. **List of abbreviations: (brainstem nuclei used as seeds are marked with red brackets:** Median Raphe nucleus (MnR), Periaqueductal Gray (PAG), Substantia Nigra-subregion1 (SN1), Substantia Nigra-subregion2 (SN2), Red nucleus-subregion1 (RN1), Red Nucleus-subregion2 (RN2), Mesencephalic Reticular formation (mRt), Cuneiform (CnF), Pedunculotegmental nuclei (PTg), Isthmic Reticular formation (isRt), Laterodorsal Tegmental Nucleus-Central Gray of the rhombencephalon (LDTg-CGPn), Pontine Reticular Nucleus, Oral Part-Pontine Reticular Nucleus Caudal Part (PnO-PnC), Locus Coeruleus (LC), Subcoeruleus nucleus (SubC), Inferior Olivary Nucleus (ION), Caudal-rostral Linear Raphe (CLi-RLi), Dorsal raphe (DR), and Paramedian Raphe nucleus (PMnR). **Additional brainstem nuclei used as targets:** Superior Colliculus (SC), Inferior Colliculus (IC), Ventral Tegmental Area-Parabrachial Pigmented Nucleus (VTA-PBP), Microcellular Tegmental Nucleus-Prabigeminal nucleus (MiTg-PBG), Lateral Parabrachial Nucleus (LPB), Medial Parabrachial Nucleus (MPB), Vestibular nuclei complex (Ve), Parvocellular Reticular nucleus Alpha (PCRtA), Superior Olivary Complex (SOC), Superior Medullary Reticular formation (sMRt), Viscero-Sensory Motor nuclei complex (VSM), Inferior Medullary Reticular formation (iMRt), Raphe Magnus (RMg), Raphe Obscurus (ROb) and Raphe Pallidus (RPa).

**Figure 11.**
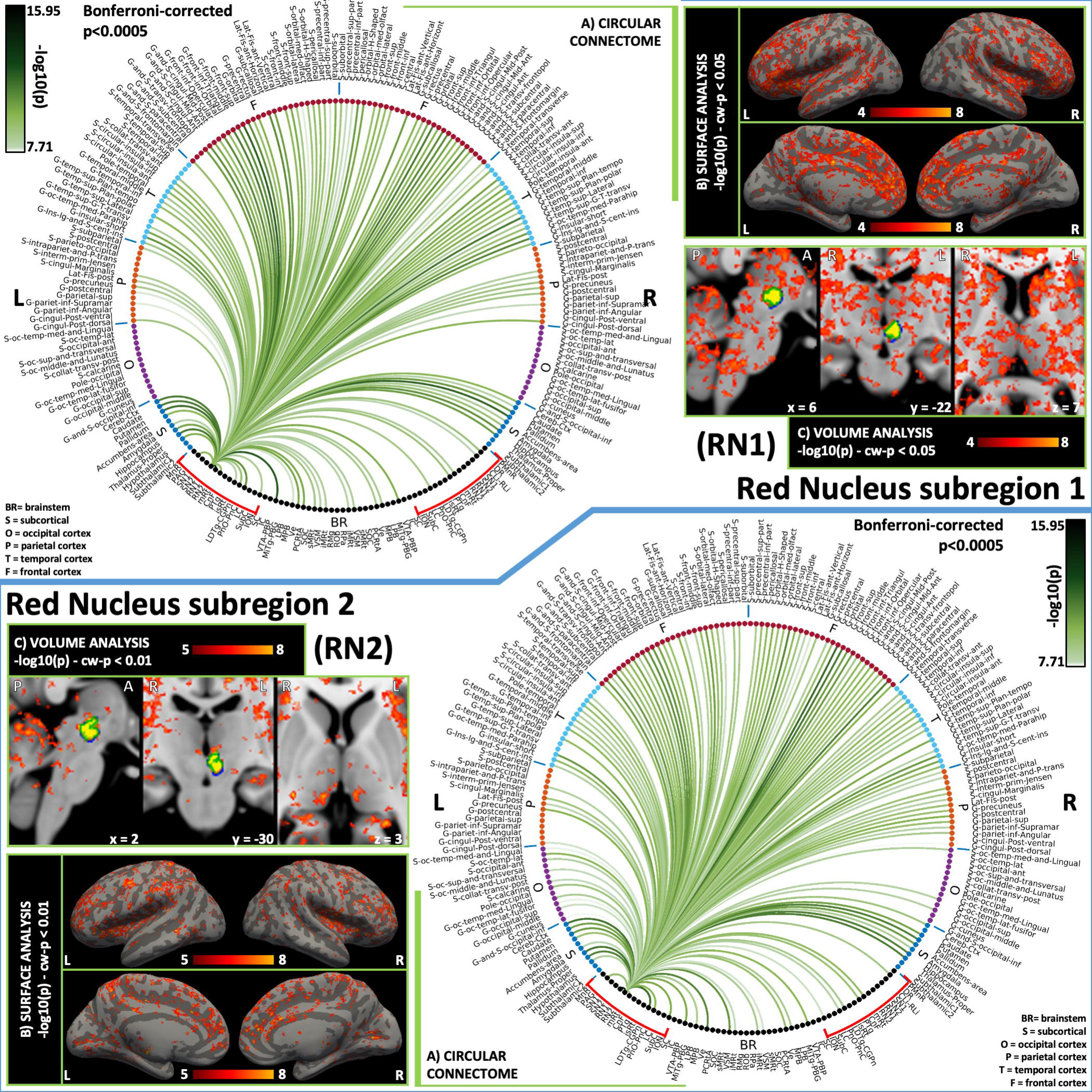
**2D circular connectome (A), voxel based connectivity maps in the cortex (B) and subcortex (C) of (top) RN1 and (bottom) RN2, which are both motor brainstem nuclei.** For RN1 we found connectivity with motor cortical areas, cerebellar cortex, ipsilateral ION and Ve among others. RN2 showed strong connectivity with ipsilateral RN1, bilateral ION, Ve, VSM, iMRt, PAG, DR, as well as frontal regions. **List of abbreviations: (brainstem nuclei used as seeds are marked with red brackets:** Median Raphe nucleus (MnR), Periaqueductal Gray (PAG), Substantia Nigra-subregion1 (SN1), Substantia Nigra-subregion2 (SN2), Red nucleus-subregion1 (RN1), Red Nucleus-subregion2 (RN2), Mesencephalic Reticular formation (mRt), Cuneiform (CnF), Pedunculotegmental nuclei (PTg), Isthmic Reticular formation (isRt), Laterodorsal Tegmental Nucleus-Central Gray of the rhombencephalon (LDTg-CGPn), Pontine Reticular Nucleus, Oral Part-Pontine Reticular Nucleus Caudal Part (PnO-PnC), Locus Coeruleus (LC), Subcoeruleus nucleus (SubC), Inferior Olivary Nucleus (ION), Caudal-rostral Linear Raphe (CLi-RLi), Dorsal raphe (DR), and Paramedian Raphe nucleus (PMnR). **Additional brainstem nuclei used as targets:** Superior Colliculus (SC), Inferior Colliculus (IC), Ventral Tegmental Area-Parabrachial Pigmented Nucleus (VTA-PBP), Microcellular Tegmental Nucleus-Prabigeminal nucleus (MiTg-PBG), Lateral Parabrachial Nucleus (LPB), Medial Parabrachial Nucleus (MPB), Vestibular nuclei complex (Ve), Parvocellular Reticular nucleus Alpha (PCRtA), Superior Olivary Complex (SOC), Superior Medullary Reticular formation (sMRt), Viscero-Sensory Motor nuclei complex (VSM), Inferior Medullary Reticular formation (iMRt), Raphe Magnus (RMg), Raphe Obscurus (ROb) and Raphe Pallidus (RPa).

**Figure 12.**
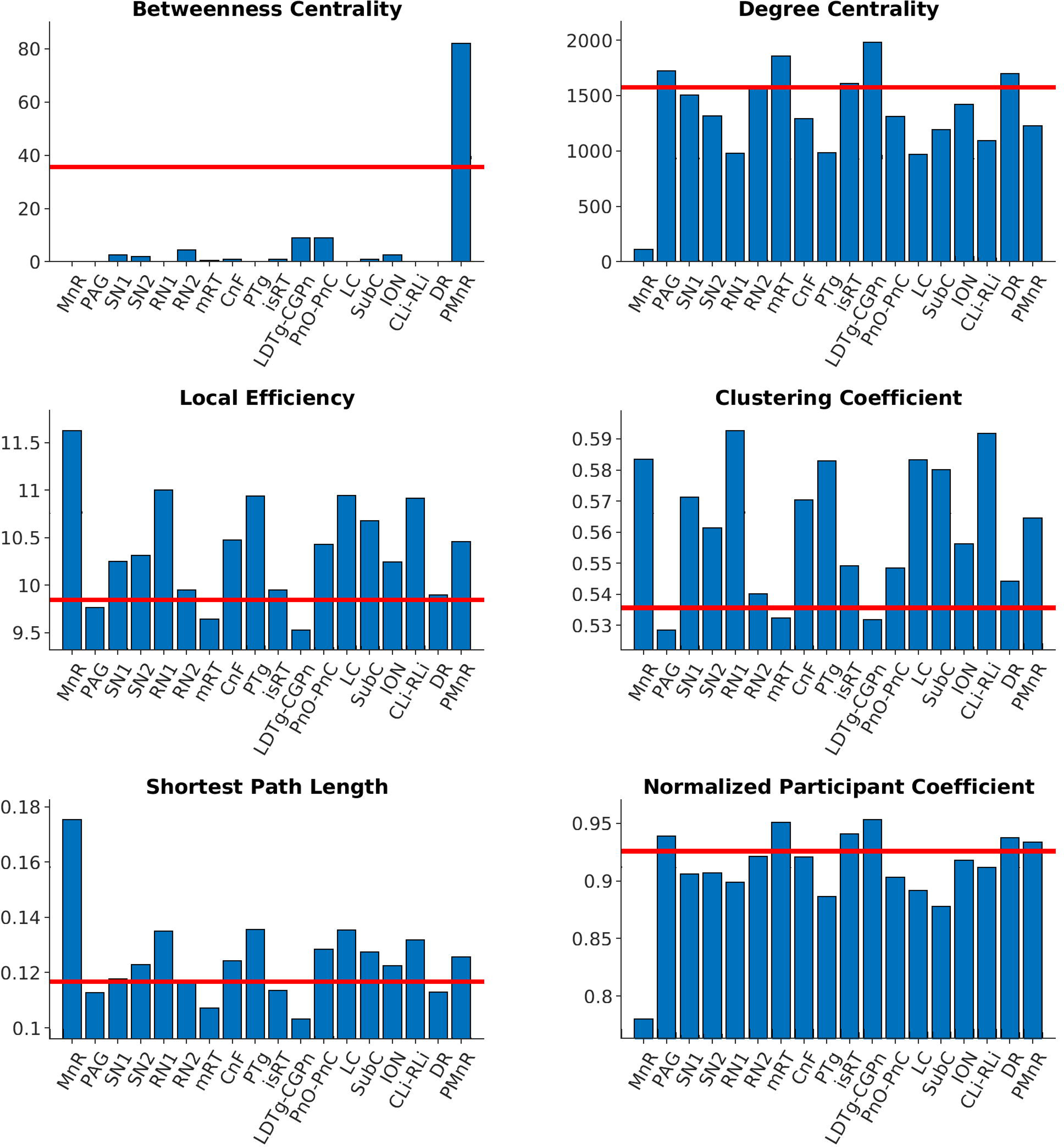
**Nodal metrics of arousal-motor network brainstem nuclei using GRETNA graph analysis (Wang et al., 2015).** Nodal metrics varied across nuclei, with PAG, mRt, isRt, LDTg-CGPn, DR showing the greatest variation (either below or above the metric average across nuclei). **List of abbreviations:** Median Raphe nucleus (MnR), Periaqueductal Gray (PAG), Substantia Nigra-subregion1 (SN1), Substantia Nigra-subregion2 (SN2), Red nucleus-subregion1 (RN1), Red Nucleus-subregion2 (RN2), Mesencephalic Reticular formation (mRt), Cuneiform (CnF), Pedunculotegmental nuclei (PTg), Isthmic Reticular formation (isRt), Laterodorsal Tegmental Nucleus-Central Gray of the rhombencephalon (LDTg-CGPn), Pontine Reticular Nucleus, Oral Part-Pontine Reticular Nucleus Caudal Part (PnO-PnC), Locus Coeruleus (LC), Subcoeruleus nucleus (SubC), Inferior Olivary Nucleus (ION), Caudal-rostral Linear Raphe (CLi-RLi), Dorsal raphe (DR), and Paramedian Raphe nucleus (PMnR).

**Figure 13.**
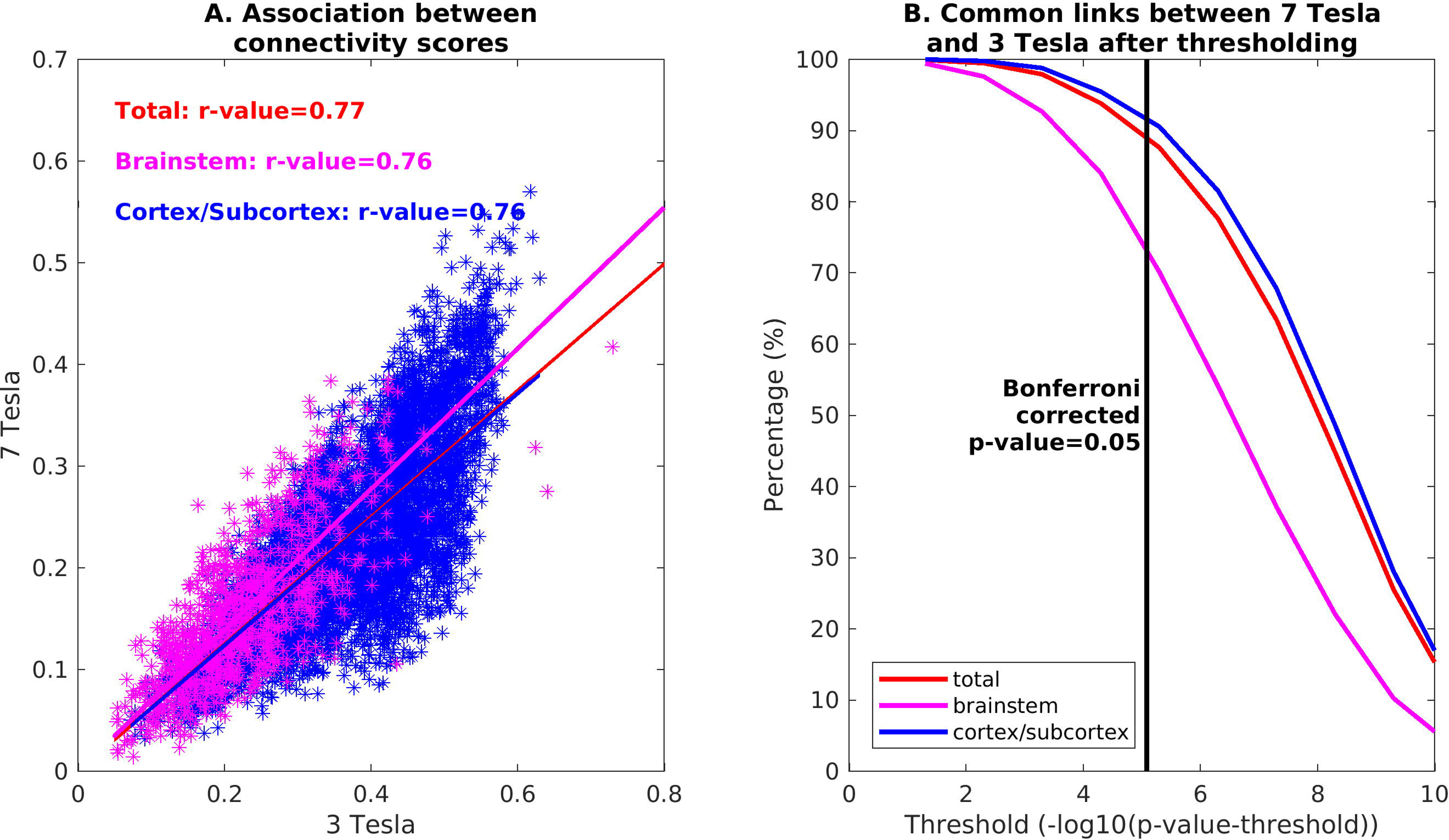
**Translatability of 7 Tesla results in a conventional dataset acquired at 3 Tesla.** (A) Association values between connectivity scores obtained at 3 Tesla and 7 Tesla for all the targets (red), brainstem only targets (magenta), and cortical/subcortical (other than brainstem) targets; (B) percentage links in common between 7 Tesla vs 3 Tesla results found in the whole brain, brainstem and cortex/subcortex.

**Figure 14.**
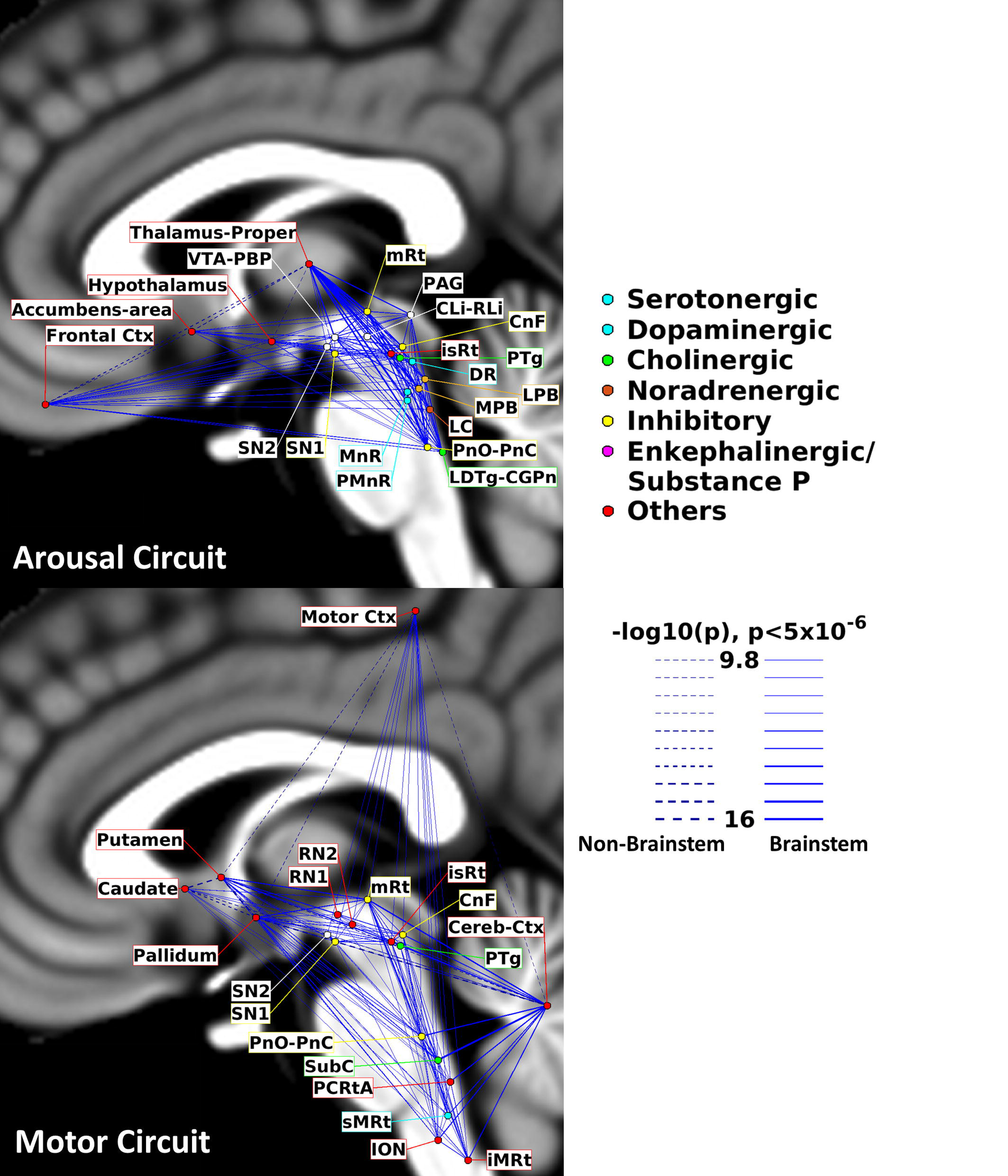
**Circuit diagram of the arousal (top) and motor (bottom) network derived from resting-state 7 Tesla fMRI functional connectivity.** The diagram of the arousal network displayed high node interconnectivity. Interestingly, several brainstem arousal nuclei displayed connectivity to the thalamus, hypothalamus, basal forebrain and frontal cortex, as expected for arousal regions. The diagram of the motor network matched expected connectivity of motor regions, including the motor cortex, basal ganglia and cerebellum. Note that the link thickness was varied based on the statistical significance of the connectivity strength. For the arousal circuit, the brainstem nodes were color-coded based on the main neurotransmitter employed.

### Diagrams of the arousal and motor networks

The diagram of the arousal network displayed high node interconnectivity. Interestingly, several brainstem arousal nuclei displayed connectivity to the thalamus, hypothalamus, basal forebrain and frontal cortex, as expect for arousal regions (Parvizi and Damasio, 2001; Saper et al., 2001, 2010; Olszewski and Baxter, 1954). The diagram of the motor network matched expected connectivity of motor regions, including the motor cortex, basal ganglia and cerebellum (Merel et al., 2019).

### Limitations

The brainstem nuclei functional connectome was developed using optimized acquisition and analysis methods for the brainstem, including high spatial resolution imaging at ultra high-field, and ad-hoc coregistration and physiological noise correction methods. Yet, limitations of current functional connectivity methods might yield residual confounds to the achieved brainstem nuclei connectome. These include the possible presence of spurious indirect connections due to the use of Pearson’s correlation coefficient and to residual physiological noise. Moreover, in spite of the high degree of confidence achieved in the image distortion correction and coregistration, residual spatial distortions and mismatch might have introduced partial volume effects in the obtained brainstem nuclei connectome. Yet, we expect these effects to be more pronounced in voxel-based analysis and to be mitigated in region-based connectivity analysis.

The choice of the threshold to be applied to connectivity matrices is nontrivial: on one side, thresholding is important to control for the effects of subject-specific spurious correlations at the group level. On the other side, the threshold, either corrected or uncorrected for multiple comparisons, is often arbitrary, and can be defined in several ways. Thresholding at the subject level, for example, is an approach that suffers from strong dependency on differences in connectivity density across subjects. In this study, we choose to perform the thresholding on the group level statistics. In literature, recent alternative approaches can be found, such as proportionally thresholding at the single subject level (van den Heuvel et al., 2017), that ensure equal network density across datasets, setting a subject-specific threshold at the subject level. Similar alternatives are interesting, and should be tested in future studies.

In this work we used TRs of 2.5 s (for 7 Tesla fMRI) and 3.5 s (for 3 Tesla MRI), which were minimum TRs for the chosen number of slices and spatial resolution. We also employed a band-pass filter with cut-off frequencies of 0.01 Hz and 0.1 Hz. Based on conventional fMRI connectivity analyses, we focused on evaluating signal fluctuations occurring over five to tens of seconds. Previous studies (Lee et al., 2013) suggest these cut-off frequencies improve the temporal signal-to-noise-ratio by broadly reducing noise related signal fluctuations. Moreover, a low-pass filtering cut-off of 0.1 Hz allows to use a TR of up to 5 s, which corresponds to 0.1 Hz Nyquist frequency. There are similar 3 Tesla fMRI studies with longer TRs (for example 6s, Horovitz et al., 2008). The use of long TRs might add chances of aliasing cardiac artefacts on the same frequencies of the BOLD signal (Huotari et al., 2019); however, most recent physiological noise correction algorithms, such as RETROICOR, model these artefacts including their aliasing effects. Also, recent studies suggest use of shorter TRs (aided by the use of simultaneous-multi-slice imaging methods) and low spatial resolutions, to investigate faster BOLD signal fluctuations and dynamic changes in functional connectivity in health and disease (Sahib et al., 2018). With the aim of evaluating the translatability of our connectome to clinical settings, we employed standard acquisition parameters, thus at 3 Tesla, for instance, we did not use simultaneous-multi-slice imaging methods.

We used a custom-built 64 channel head coil for 3 Tesla data acquisition as opposed to widely used 32 channel commercial coils. Based on Keil 2013 (Keil et al., 2013), in the brainstem, at 3 Tesla, the sensitivity of the custom-built 64 channel coil did not vary from that of a 32 channel coil, while the former provided some sensitivity increase in the outer cortex (∼30 %, see (Keil et al., 2013)). Thus, for brainstem-to-brainstem connectivity, the use of a 64 channel coil at 3 Tesla does not limit the generalizability of the 3 Tesla findings to more widely used 32 channel coils; yet, a non negligible increase in sensitivity in the cortex, might have affected the brainstem-to-cortical connectivity results.

Literature indicates resting-state fMRI studies have been done with either eyes closed or eyes open condition, which has been shown to affect connectivity results (Agcaoglu et al., 2019; Bianciardi et al., 2009; Costumero et al., 2020; Patriat et al., 2013). For instance, auditory and sensorimotor networks showed increased connectivity during the eyes open condition compared to the eyes closed one (Agcaoglu et al., 2019; Patriat et al., 2013). Another study showed increased connectivity of occipital areas with eyes closed condition (Bianciardi et al., 2009; Costumero et al., 2020), while (Agcaoglu et al., 2019; Patriat et al., 2013) showed similar results with eyes open condition. Our current findings showed lower connectivity of seeds to occipital regions. Based on above evidences, we recognize our connectivity results are based on widely used eyes-closed state and might be influenced by the adopted condition. Future studies might investigate time-varying connectivity measures also in relation to physiological autonomic and arousal oscillations recorded by peripheral probes.

### Conclusions

Brainstem nuclei have been shown to play a pivotal role in many functions, such as sleep, arousal, wakefulness, autonomic, motor, and nociception. (Bianciardi et al., 2016; Merel et al., 2019; Saper et al., 2001; Satpute et al., 2013). These nuclei work in a coordinated fashion, so it becomes imperative to map their connectome to fully understand these processes. Further, graph analysis has received much attention in connectome research (Tomasi and Volkow, 2011; van den Heuvel and Sporns, 2013) to study integrative processing (Heuvel et al., 2012) and related pathological conditions (Saper et al., 2010). Hub regions are central in the connectome, route information exchange, and achieve integration of otherwise segregated processes (de Reus and van den Heuvel, 2014; van den Heuvel and Sporns, 2013). In the present work, we provided a connectome of arousal-motor brainstem nuclei along with the diagrams of the resulting arousal and motor resting state functional networks in living humans, to provide a comprehensive yet segregated view on these processes. This work was based on recently developed high-resolution 7 Tesla MRI based probabilistic brainstem atlas in living humans (Bianciardi et al., 2018, 2015; García-Gomar et al., 2019; Singh et al., 2019). Further, we demonstrated the translatability of our connectome in conventional imaging by comparing 3 Tesla data connectome results to 7 Tesla results. We speculate that these results might provide a baseline for future studies of brainstem connectivity in heath and disease.

## Supporting information

https://figshare.com/s/22cfc0708ab8477c5018

## Acknowledgements

This work was funded by grants from the National Institutes of Health (NIBIB-K01EB019474, NIDCD-R21DC015888, NIA-R01AG063982), from the MGH Claflin Award, and Harvard Mind Brain Behavior.

## List of abbreviations of brainstem nuclei used as seeds

(MnR): Median Raphe nucleus
(PAG): Periaqueductal Gray
(SN1): Substantia Nigra-subregion1
(SN2): Substantia Nigra-subregion2
(RN1): Red nucleus-subregion1
(RN2): Red Nucleus-subregion2
(mRt): Mesencephalic Reticular formation
(CnF): Cuneiform
(PTg): Pedunculotegmental nuclei
(isRt): Isthmic Reticular formation
(LDTg-CGPn): Laterodorsal Tegmental Nucleus- Central Gray of the rhombencephalon
(PnO-PnC): Pontine Reticular Nucleus Oral Part- Pontine Reticular Nucleus Caudal Part
(LC): Locus Coeruleus
(SubC): Subcoeruleus nucleus
(ION): Inferior Olivary Nucleus
(CLi-RLi): Caudal-rostral Linear Raphe
(DR): Dorsal raphe
(PMnR): Paramedian Raphe nucleus

**List of abbreviations of additional brainstem nuclei used as targets**
(SC): Superior Colliculus
(IC): Inferior Colliculus
(VTA-PBP): Ventral Tegmental Area-Parabrachial Pigmented Nucleus
(MiTg-PBG): Microcellular Tegmental Nucleus-Prabigeminal nucleus
(LPB): Lateral Parabrachial Nucleus
(MPB): Medial Parabrachial Nucleus
(Ve): Vestibular nuclei complex
(PCRtA): Parvocellular Reticular nucleus Alpha
(SOC): Superior Olivary Complex
(sMRt): Superior Medullary Reticular formation
(VSM): Viscero-Sensory Motor nuclei complex
(iMRt): Inferior Medullary Reticular formation
(RMg): Raphe Magnus
(ROb): Raphe Obscurus
(RPa): Raphe Pallidus

## References

Achard, S., Bullmore, E., 2007. Efficiency and Cost of Economical Brain Functional Networks. PLOS Computational Biology 3, e17. https://doi.org/10.1371/journal.pcbi.0030017

Afifi, A.K., Bahuth, N.B., Kaelber, W.W., Mikhael, E., Nassar, S., 1974. The cortico-nigral fibre tract. An experimental Fink-Heimer study in cats. J Anat 118, 469–476.

Agcaoglu, O., Wilson, T.W., Wang, Y., Stephen, J., Calhoun, V.D., 2019. Resting state connectivity differences in eyes open versus eyes closed conditions. Human Brain Mapping 40, 2488–2498. https://doi.org/10.1002/hbm.24539

Amunts, K., Kedo, O., Kindler, M., Pieperhoff, P., Mohlberg, H., Shah, N.J., Habel, U., Schneider, F., Zilles, K., 2005. Cytoarchitectonic mapping of the human amygdala, hippocampal region and entorhinal cortex: intersubject variability and probability maps. Anat Embryol (Berl) 210, 343–352. https://doi.org/10.1007/s00429-005-0025-5

Aquino, K.M., Fulcher, B.D., Parkes, L., Sabaroedin, K., Fornito, A., 2020. Identifying and removing widespread signal deflections from fMRI data: Rethinking the global signal regression problem. Neuroimage 212, 116614. https://doi.org/10.1016/j.neuroimage.2020.116614

Arnsten, A.F., Goldman-Rakic, P.S., 1984. Selective prefrontal cortical projections to the region of the locus coeruleus and raphe nuclei in the rhesus monkey. Brain Res 306, 9–18. https://doi.org/10.1016/0006-8993(84)90351-2

Aston-Jones, G., Shipley, M.T., Chouvet, G., Ennis, M., van Bockstaele, E., Pieribone, V., Shiekhattar, R., Akaoka, H., Drolet, G., Astier, B., Charléty, P., Valentino, R.J., Williams, J.T., 1991. Afferent regulation of locus coeruleus neurons: anatomy, physiology and pharmacology, in: Progress in Brain Research. Elsevier, pp. 47–75. https://doi.org/10.1016/S0079-6123(08)63799-1

Auer, D.P., 2008. Spontaneous low-frequency blood oxygenation level-dependent fluctuations and functional connectivity analysis of the “resting” brain. Magn Reson Imaging 26, 1055–1064. https://doi.org/10.1016/j.mri.2008.05.008

Bajic, D., Proudfit, H.K., 1999. Projections of neurons in the periaqueductal gray to pontine and medullary catecholamine cell groups involved in the modulation of nociception. J Comp Neurol 405, 359–379.

Balaban, C.D., 2004. Projections from the parabrachial nucleus to the vestibular nuclei: potential substrates for autonomic and limbic influences on vestibular responses. Brain Research 996, 126–137. https://doi.org/10.1016/j.brainres.2003.10.026

Bär, K.-J., de la Cruz, F., Schumann, A., Koehler, S., Sauer, H., Critchley, H., Wagner, G., 2016. Functional connectivity and network analysis of midbrain and brainstem nuclei. Neuroimage 134, 53–63. https://doi.org/10.1016/j.neuroimage.2016.03.071

Barmack, N.H., 2006. Inferior olive and oculomotor system, in: Progress in Brain Research. Elsevier, pp. 269–291. https://doi.org/10.1016/S0079-6123(05)51009-4

Beckstead, R.M., Domesick, V.B., Nauta, W.J., 1979. Efferent connections of the substantia nigra and ventral tegmental area in the rat. Brain Res 175, 191–217. https://doi.org/10.1016/0006-8993(79)91001-1

Behzadi, G., Kalén, P., Parvopassu, F., Wiklund, L., 1990. Afferents to the median raphe nucleus of the rat: retrograde cholera toxin and wheat germ conjugated horseradish peroxidase tracing, and selective D-[3H]aspartate labelling of possible excitatory amino acid inputs. Neuroscience 37, 77–100. https://doi.org/10.1016/0306-4522(90)90194-9

Beitz, A.J., 1982. The nuclei of origin of brain stem enkephalin and substance P projections to the rodent nucleus raphe magnus. Neuroscience 7, 2753–2768. https://doi.org/10.1016/0306-4522(82)90098-7

Beliveau, V., Svarer, C., Frokjaer, V.G., Knudsen, G.M., Greve, D.N., Fisher, P.M., 2015. Functional connectivity of the dorsal and median raphe nuclei at rest. Neuroimage 116, 187–195. https://doi.org/10.1016/j.neuroimage.2015.04.065

Bianciardi, M., Fukunaga, M., van Gelderen, P., Horovitz, S.G., de Zwart, J.A., Duyn, J.H., 2009. Modulation of spontaneous fMRI activity in human visual cortex by behavioral state. NeuroImage 45, 160–168. https://doi.org/10.1016/j.neuroimage.2008.10.034

Bianciardi, M., Strong, C., Toschi, N., Edlow, B.L., Fischl, B., Brown, E.N., Rosen, B.R., Wald, L.L., 2018. A probabilistic template of human mesopontine tegmental nuclei from in vivo 7T MRI. Neuroimage 170, 222–230. https://doi.org/10.1016/j.neuroimage.2017.04.070

Bianciardi, M., Toschi, N., Edlow, B.L., Eichner, C., Setsompop, K., Polimeni, J.R., Brown, E.N., Kinney, H.C., Rosen, B.R., Wald, L.L., 2015. Toward an In Vivo Neuroimaging Template of Human Brainstem Nuclei of the Ascending Arousal, Autonomic, and Motor Systems. Brain Connect 5, 597–607. https://doi.org/10.1089/brain.2015.0347

Bianciardi, M., Toschi, N., Eichner, C., Polimeni, J.R., Setsompop, K., Brown, E.N., Hämäläinen, M.S., Rosen, B.R., Wald, L.L., 2016. In vivo functional connectome of human brainstem nuclei of the ascending arousal, autonomic, and motor systems by high spatial resolution 7-Tesla fMRI. MAGMA 29, 451–462. https://doi.org/10.1007/s10334-016-0546-3

Birn, R.M., Smith, M.A., Jones, T.B., Bandettini, P.A., 2008. The respiration response function: The temporal dynamics of fMRI signal fluctuations related to changes in respiration. NeuroImage 40, 644–654. https://doi.org/10.1016/j.neuroimage.2007.11.059

Biswal, B., Yetkin, F.Z., Haughton, V.M., Hyde, J.S., 1995. Functional connectivity in the motor cortex of resting human brain using echo-planar MRI. Magn Reson Med 34, 537–541. https://doi.org/10.1002/mrm.1910340409

Bobillier, P., Seguin, S., Petitjean, F., Salvert, D., Touret, M., Jouvet, M., 1976. The raphe nuclei of the cat brain stem: a topographical atlas of their efferent projections as revealed by autoradiography. Brain Res 113, 449–486. https://doi.org/10.1016/0006-8993(76)90050-0

Boeve, B.F., Silber, M.H., Saper, C.B., Ferman, T.J., Dickson, D.W., Parisi, J.E., Benarroch, E.E., Ahlskog, J.E., Smith, G.E., Caselli, R.C., Tippman-Peikert, M., Olson, E.J., Lin, S.-C., Young, T., Wszolek, Z., Schenck, C.H., Mahowald, M.W., Castillo, P.R., Del Tredici, K., Braak, H., 2007. Pathophysiology of REM sleep behaviour disorder and relevance to neurodegenerative disease. Brain 130, 2770–2788. https://doi.org/10.1093/brain/awm056

Bryant, R.A., Harvey, A.G., Guthrie, R.M., Moulds, M.L., 2000. A prospective study of psychophysiological arousal, acute stress disorder, and posttraumatic stress disorder. J Abnorm Psychol 109, 341–344.

Bullmore, E., Sporns, O., 2009. Complex brain networks: graph theoretical analysis of structural and functional systems. Nature Reviews Neuroscience 10, 186–198. https://doi.org/10.1038/nrn2575

Cacciola, A., Milardi, D., Anastasi, G.P., Basile, G.A., Ciolli, P., Irrera, M., Cutroneo, G., Bruschetta, D., Rizzo, G., Mondello, S., Bramanti, P., Quartarone, A., 2016. A Direct Cortico-Nigral Pathway as Revealed by Constrained Spherical Deconvolution Tractography in Humans. Front Hum Neurosci 10, 374. https://doi.org/10.3389/fnhum.2016.00374

Cacciola, Alberto, Milardi, D., Basile, G.A., Bertino, S., Calamuneri, A., Chillemi, G., Paladina, G., Impellizzeri, F., Trimarchi, F., Anastasi, G., Bramanti, A., Rizzo, G., 2019. The cortico-rubral and cerebello-rubral pathways are topographically organized within the human red nucleus. Sci Rep 9, 12117. https://doi.org/10.1038/s41598-019-48164-7

Cacciola, A., Milardi, D., Basile, G.A., Bertino, S., Calamuneri, A., Chillemi, G., Paladina, G., Impellizzeri, F., Trimarchi, F., Anastasi, G., Bramanti, A., Rizzo, G., 2019. The cortico-rubral and cerebello-rubral pathways are topographically organized within the human red nucleus. Sci Rep 9, 12117. https://doi.org/10.1038/s41598-019-48164-7

Cameron, A.A., Khan, I.A., Westlund, K.N., Cliffer, K.D., Willis, W.D., 1995. The efferent projections of the periaqueductal gray in the rat: a Phaseolus vulgaris-leucoagglutinin study. I. Ascending projections. J Comp Neurol 351, 568–584. https://doi.org/10.1002/cne.903510407

Carlsson, A., 1995. Neurocircuitries and neurotransmitter interactions in schizophrenia. Int Clin Psychopharmacol 10 Suppl 3, 21–28.

Chai, C.Y., Lin, R.H., Lin, A.M., Pan, C.M., Lee, E.H., Kuo, J.S., 1988. Pressor responses from electrical or glutamate stimulations of the dorsal or ventrolateral medulla. Am J Physiol 255, R709–717. https://doi.org/10.1152/ajpregu.1988.255.5.R709

Chang, C., Cunningham, J.P., Glover, G.H., 2009. Influence of heart rate on the BOLD signal: The cardiac response function. NeuroImage 44, 857–869. https://doi.org/10.1016/j.neuroimage.2008.09.029

Chang, S.J., Cajigas, I., Opris, I., Guest, J.D., Noga, B.R., 2020. Dissecting Brainstem Locomotor Circuits: Converging Evidence for Cuneiform Nucleus Stimulation. Front. Syst. Neurosci. 14, 64. https://doi.org/10.3389/fnsys.2020.00064

Cheriyan, J., Sheets, P.L., 2018. Altered Excitability and Local Connectivity of mPFC-PAG Neurons in a Mouse Model of Neuropathic Pain. J Neurosci 38, 4829–4839. https://doi.org/10.1523/JNEUROSCI.2731-17.2018

Coimbra, B., Soares-Cunha, C., Borges, S., Vasconcelos, N.A., Sousa, N., Rodrigues, A.J., 2017. Impairments in laterodorsal tegmentum to VTA projections underlie glucocorticoid-triggered reward deficits. eLife 6, e25843. https://doi.org/10.7554/eLife.25843

Coimbra, B., Soares-Cunha, C., Vasconcelos, N.A.P., Domingues, A.V., Borges, S., Sousa, N., Rodrigues, A.J., 2019. Role of laterodorsal tegmentum projections to nucleus accumbens in reward-related behaviors. Nat Commun 10, 4138. https://doi.org/10.1038/s41467-019-11557-3

Cordes, D., Haughton, V.M., Arfanakis, K., Carew, J.D., Turski, P.A., Moritz, C.H., Quigley, M.A., Meyerand, M.E., 2001. Frequencies contributing to functional connectivity in the cerebral cortex in “resting-state” data. AJNR Am J Neuroradiol 22, 1326–1333.

Costumero, V., Bueichekú, E., Adrián-Ventura, J., Ávila, C., 2020. Opening or closing eyes at rest modulates the functional connectivity of V1 with default and salience networks. Scientific Reports 10, 9137. https://doi.org/10.1038/s41598-020-66100-y

Coulombe, M.-A., Erpelding, N., Kucyi, A., Davis, K.D., 2016. Intrinsic functional connectivity of periaqueductal gray subregions in humans. Hum Brain Mapp 37, 1514–1530. https://doi.org/10.1002/hbm.23117

Dampney, R., 2018. Emotion and the Cardiovascular System: Postulated Role of Inputs From the Medial Prefrontal Cortex to the Dorsolateral Periaqueductal Gray. Frontiers in Neuroscience 12, 343. https://doi.org/10.3389/fnins.2018.00343

Datta, S., Curró Dossi, R., Paré, D., Oakson, G., Steriade, M., 1991. Substantia nigra reticulata neurons during sleep-waking states: relation with ponto-geniculo-occipital waves. Brain Res 566, 344–347. https://doi.org/10.1016/0006-8993(91)91723-e

Datta, S., Siwek, D.F., Patterson, E.H., Cipolloni, P.B., 1998. Localization of pontine PGO wave generation sites and their anatomical projections in the rat. Synapse 30, 409–423. https://doi.org/10.1002/(SICI)1098-2396(199812)30:4<409::AID-SYN8>3.0.CO;2-#

Dautan, D., Hacioğlu Bay, H., Bolam, J.P., Gerdjikov, T.V., Mena-Segovia, J., 2016. Extrinsic Sources of Cholinergic Innervation of the Striatal Complex: A Whole-Brain Mapping Analysis. Front. Neuroanat. 10. https://doi.org/10.3389/fnana.2016.00001

de Lacalle, S., Saper, C.B., 2000. Calcitonin gene-related peptide-like immunoreactivity marks putative visceral sensory pathways in human brain. Neuroscience 100, 115–130. https://doi.org/10.1016/S0306-4522(00)00245-1

de Reus, M.A., van den Heuvel, M.P., 2014. Simulated rich club lesioning in brain networks: a scaffold for communication and integration? Front. Hum. Neurosci. 8. https://doi.org/10.3389/fnhum.2014.00647

Demirci, O., Stevens, M.C., Andreasen, N.C., Michael, A., Liu, J., White, T., Pearlson, G.D., Clark, V.P., Calhoun, V.D., 2009. Investigation of relationships between fMRI brain networks in the spectral domain using ICA and Granger causality reveals distinct differences between schizophrenia patients and healthy controls. Neuroimage 46, 419–431. https://doi.org/10.1016/j.neuroimage.2009.02.014

Destrieux, C., Fischl, B., Dale, A., Halgren, E., 2010. Automatic parcellation of human cortical gyri and sulci using standard anatomical nomenclature. Neuroimage 53, 1–15. https://doi.org/10.1016/j.neuroimage.2010.06.010

Dobbs, L.K., Mark, G.P., 2012. Acetylcholine from the mesopontine tegmental nuclei differentially affects methamphetamine induced locomotor activity and neurotransmitter levels in the mesolimbic pathway. Behavioural Brain Research 226, 224–234. https://doi.org/10.1016/j.bbr.2011.09.022

Edwards, S.B., de Olmos, J.S., 1976. Autoradiographic studies of the projections of the midbrain reticular formation: ascending projections of nucleus cuneiformis. J Comp Neurol 165, 417–431. https://doi.org/10.1002/cne.901650403

Elvsåshagen, T., Bahrami, S., van der Meer, D., Agartz, I., Alnæs, D., Barch, D.M., Baur-Streubel, R., Bertolino, A., Beyer, M.K., Blasi, G., Borgwardt, S., Boye, B., Buitelaar, J., Bøen, E., Celius, E.G., Cervenka, S., Conzelmann, A., Coynel, D., Di Carlo, P., Djurovic, S., Eisenacher, S., Espeseth, T., Fatouros-Bergman, H., Flyckt, L., Franke, B., Frei, O., Gelao, B., Harbo, H.F., Hartman, C.A., Håberg, A., Heslenfeld, D., Hoekstra, P.J., Høgestøl, E.A., Jonassen, R., Jönsson, E.G., Kirsch, P., Kłoszewska, I., Lagerberg, T.V., Landrø, N.I., Hellard, S.L., Lesch, K.-P., Maglanoc, L.A., Malt, U.F., Mecocci, P., Melle, I., Meyer-Lindenberg, A., Moberget, T., Nordvik, J.E., Nyberg, L., Connell, K.S.O., Oosterlaan, J., Papalino, M., Papassotiropoulos, A., Pauli, P., Pergola, G., Persson, K., Quervain D. de, Reif, A., Rokicki, J., Rooij D. van, Shadrin, A.A., Schmidt, A., Schwarz, E., Selbæk, G., Soininen, H., Sowa, P., Steen, V.M., Tsolaki, M., Vellas, B., Wang, L., Westman, E., Ziegler, G.C., Zink, M., Andreassen, O.A., Westlye, L.T., Kaufmann, T., 2020. The genetic architecture of human brainstem structures and their involvement in common brain disorders. Nature Communications 11, 4016. https://doi.org/10.1038/s41467-020-17376-1

Erro, E., Lanciego, J.L., Giménez-Amaya, J.M., 1999. Relationships between thalamostriatal neurons and pedunculopontine projections to the thalamus: a neuroanatomical tract-tracing study in the rat. Experimental Brain Research 127, 162–170. https://doi.org/10.1007/s002210050786

Eser, R.A., Ehrenberg, A.J., Petersen, C., Dunlop, S., Mejia, M.B., Suemoto, C.K., Walsh, C.M., Rajana, H., Oh, J., Theofilas, P., Seeley, W.W., Miller, B.L., Neylan, T.C., Heinsen, H., Grinberg, L.T., 2018. Selective Vulnerability of Brainstem Nuclei in Distinct Tauopathies: A Postmortem Study. Journal of Neuropathology & Experimental Neurology 77, 149– 161. https://doi.org/10.1093/jnen/nlx113

Frackowiak, R.S.J., Ashburner, J.T., Penny, W.D., Zeki, S., 2004. Human Brain Function, Second Edition, 2nd edition. ed. Academic Press, Amsterdam ; Boston.

French, I.T., Muthusamy, K.A., 2018. A Review of the Pedunculopontine Nucleus in Parkinson’s Disease. Front. Aging Neurosci. 10, 99. https://doi.org/10.3389/fnagi.2018.00099

Garcia-Gomar MG, Singh K, Bianciardi M, 2021. Probabilistic structural atlas and connectome of brainstem nuclei involved in arousal and sleep by 7 Tesla MRI in living humans. ISMRM 29th Annual Meeting & Exhibition.

Garcia-Gomar, M.G., Singh, K., Bianciardi, M., 2021. Probabilistic structural atlas and connectome of brainstem nuclei involved in arousal and sleep by 7 Tesla MRI in living humans, in: ISMRM 29th Annual Meeting & Exhibition.

García-Gomar, M.G., Strong, C., Toschi, N., Singh, K., Rosen, B.R., Wald, L.L., Bianciardi, M., 2019a. In vivo Probabilistic Structural Atlas of the Inferior and Superior Colliculi, Medial and Lateral Geniculate Nuclei and Superior Olivary Complex in Humans Based on 7 Tesla MRI. Front Neurosci 13, 764. https://doi.org/10.3389/fnins.2019.00764

García-Gomar, M.G., Strong, C., Toschi, N., Singh, K., Rosen, B.R., Wald, L.L., Bianciardi, M., 2019b. In vivo Probabilistic Structural Atlas of the Inferior and Superior Colliculi, Medial and Lateral Geniculate Nuclei and Superior Olivary Complex in Humans Based on 7 Tesla MRI. Front Neurosci 13, 764. https://doi.org/10.3389/fnins.2019.00764

Garzón, M., Vaughan, R.A., Uhl, G.R., Kuhar, M.J., Pickel, V.M., 1999. Cholinergic axon terminals in the ventral tegmental area target a subpopulation of neurons expressing low levels of the dopamine transporter. J Comp Neurol 410, 197–210. https://doi.org/10.1002/(sici)1096-9861(19990726)410:2<197::aid-cne3>3.0.co;2-d

Glover, G.H., Li, T.Q., Ress, D., 2000. Image-based method for retrospective correction of physiological motion effects in fMRI: RETROICOR. Magn Reson Med 44, 162–167. https://doi.org/10.1002/1522-2594(200007)44:1<162::aid-mrm23>3.0.co;2-e

Goldberg, J.M., Wilson, V.J., Cullen, K.E., Angelaki, D.E., Broussard, D.M., Buttner-Ennever, J., Fukushima, K., Minor, L.B., 2012. The Vestibular System: A Sixth Sense. Oxford University Press. https://doi.org/10.1093/acprof:oso/9780195167085.001.0001

Gong, G., He, Y., Concha, L., Lebel, C., Gross, D.W., Evans, A.C., Beaulieu, C., 2009. Mapping anatomical connectivity patterns of human cerebral cortex using in vivo diffusion tensor imaging tractography. Cereb Cortex 19, 524–536. https://doi.org/10.1093/cercor/bhn102

Gould, E., Woolf, N.J., Butcher, L.L., 1989. Cholinergic projections to the substantia nigra from the pedunculopontine and laterodorsal tegmental nuclei. Neuroscience 28, 611–623. https://doi.org/10.1016/0306-4522(89)90008-0

Grabner, G., Janke, A.L., Budge, M.M., Smith, D., Pruessner, J., Collins, D.L., 2006. Symmetric atlasing and model based segmentation: an application to the hippocampus in older adults. Med Image Comput Comput Assist Interv 9, 58–66. https://doi.org/10.1007/11866763_8

Haller, J., Tóth, M., Halász, J., 2005. The activation of raphe serotonergic neurons in normal and hypoarousal-driven aggression: a double labeling study in rats. Behav Brain Res 161, 88– 94. https://doi.org/10.1016/j.bbr.2005.01.006

Halliday, G., Reyes, S., Double, K., 2012. Chapter 13 - Substantia Nigra, Ventral Tegmental Area, and Retrorubral Fields, in: Mai, J.K., Paxinos, G. (Eds.), The Human Nervous System (Third Edition). Academic Press, San Diego, pp. 439–455. https://doi.org/10.1016/B978-0-12-374236-0.10013-6

Hazrati, L.-N., Parent, A., 1992. Projection from the deep cerebellar nuclei to the pedunculopontine nucleus in the squirrel monkey. Brain Research 585, 267–271. https://doi.org/10.1016/0006-8993(92)91216-2

Heuvel, M., Kahn, R., Goñi, J., Sporns, O., 2012. High-cost, high-capacity backbone for global brain communication. Proceedings of the National Academy of Sciences of the United States of America 109, 11372–7. https://doi.org/10.1073/pnas.1203593109

Höglinger, G.U., Alvarez-Fischer, D., Arias-Carrión, O., Djufri, M., Windolph, A., Keber, U., Borta, A., Ries, V., Schwarting, R.K.W., Scheller, D., Oertel, W.H., 2015. A new dopaminergic nigro-olfactory projection. Acta Neuropathol 130, 333–348. https://doi.org/10.1007/s00401-015-1451-y

Holstege, Gert, 1991. Chapter 14 Descending motor pathways and the spinal motor system: Limbic and non-limbic components, in: Progress in Brain Research. Elsevier, pp. 307– 421. https://doi.org/10.1016/S0079-6123(08)63057-5

Holstege, G., 1991. Descending motor pathways and the spinal motor system: limbic and non-limbic components. Prog Brain Res 87, 307–421. https://doi.org/10.1016/s0079-6123(08)63057-5

Holstege, G., 1989. Anatomical study of the final common pathway for vocalization in the cat. J Comp Neurol 284, 242–252. https://doi.org/10.1002/cne.902840208

Holstege, G., 1988. Brainstem-spinal cord projections in the cat, related to control of head and axial movements. Rev Oculomot Res 2, 431–470.

Holstege, G., Cowie, R.J., 1989. Projections from the rostral mesencephalic reticular formation to the spinal cord: An HRP and autoradiographical tracing study in the cat. Exp Brain Res 75. https://doi.org/10.1007/BF00247933

Hornung, J.-P., 2003. The human raphe nuclei and the serotonergic system. J Chem Neuroanat 26, 331–343. https://doi.org/10.1016/j.jchemneu.2003.10.002

Horovitz, S.G., Fukunaga, M., de Zwart, J.A., van Gelderen, P., Fulton, S.C., Balkin, T.J., Duyn, J.H., 2008. Low frequency BOLD fluctuations during resting wakefulness and light sleep: A simultaneous EEG-fMRI study. Human Brain Mapping 29, 671–682. https://doi.org/10.1002/hbm.20428

Huerta, M.F., Kaas, J.H., 1990. Supplementary eye field as defined by intracortical microstimulation: Connections in macaques. J. Comp. Neurol. 293, 299–330. https://doi.org/10.1002/cne.902930211

Huerta, Michael F., Krubitzer, L.A., Kaas, J.H., 1986. Frontal eye field as defined by intracortical microstimulation in squirrel monkeys, owl monkeys, and macaque monkeys: I. Subcortical connections. J. Comp. Neurol. 253, 415–439. https://doi.org/10.1002/cne.902530402

Huerta, M.F., Krubitzer, L.A., Kaas, J.H., 1986. Frontal eye field as defined by intracortical microstimulation in squirrel monkeys, owl monkeys, and macaque monkeys: I. Subcortical connections. J. Comp. Neurol 253, 415–439. https://doi.org/10.1002/cne.902530402

Huotari, N., Raitamaa, L., Helakari, H., Kananen, J., Raatikainen, V., Rasila, A., Tuovinen, T., Kantola, J., Borchardt, V., Kiviniemi, V.J., Korhonen, V.O., 2019. Sampling Rate Effects on Resting State fMRI Metrics. Frontiers in Neuroscience 13. https://doi.org/10.3389/fnins.2019.00279

Ikemoto, S., 2007. Dopamine reward circuitry: two projection systems from the ventral midbrain to the nucleus accumbens-olfactory tubercle complex. Brain Res Rev 56, 27– 78. https://doi.org/10.1016/j.brainresrev.2007.05.004

Ilinsky, I.A., Jouandet, M.L., Goldman-Rakic, P.S., 1985. Organization of the nigrothalamocortical system in the rhesus monkey. J. Comp. Neurol. 236, 315–330. https://doi.org/10.1002/cne.902360304

Illing, R.B., Kraus, K.S., Michler, S.A., 2000. Plasticity of the superior olivary complex. Microsc Res Tech 51, 364–381. https://doi.org/10.1002/1097-0029(20001115)51:4<364::AID-JEMT6>3.0.CO;2-E

Imai, H., Steindler, D.A., Kitai, S.T., 1986. The organization of divergent axonal projections from the midbrain raphe nuclei in the rat. J. Comp. Neurol. 243, 363–380. https://doi.org/10.1002/cne.902430307

Indovina, I., Bosco, G., Riccelli, R., Maffei, V., Lacquaniti, F., Passamonti, L., Toschi, N., 2020. Structural connectome and connectivity lateralization of the multimodal vestibular cortical network. NeuroImage 222, 117247. https://doi.org/10.1016/j.neuroimage.2020.117247

Jacobs, B.L., Azmitia, E.C., 1992. Structure and function of the brain serotonin system. Physiological Reviews 72, 165–229. https://doi.org/10.1152/physrev.1992.72.1.165

Kalén, P., Karlson, M., Wiklund, L., 1985. Possible excitatory amino acid afferents to nucleus raphe dorsalis of the rat investigated with retrograde wheat germ agglutinin and d-[3H]aspartate tracing. Brain Research 360, 285–297. https://doi.org/10.1016/0006-8993(85)91244-2

Karlsson, K.Æ., Gall, A.J., Mohns, E.J., Seelke, A.M.H., Blumberg, M.S., 2005. The Neural Substrates of Infant Sleep in Rats. PLoS Biol 3, e143. https://doi.org/10.1371/journal.pbio.0030143

Kaur, S., Pedersen, N.P., Yokota, S., Hur, E.E., Fuller, P.M., Lazarus, M., Chamberlin, N.L., Saper, C.B., 2013a. Glutamatergic signaling from the parabrachial nucleus plays a critical role in hypercapnic arousal. J Neurosci 33, 7627–7640. https://doi.org/10.1523/JNEUROSCI.0173-13.2013

Kaur, S., Pedersen, N.P., Yokota, S., Hur, E.E., Fuller, P.M., Lazarus, M., Chamberlin, N.L., Saper, C.B., 2013b. Glutamatergic signaling from the parabrachial nucleus plays a critical role in hypercapnic arousal. J. Neurosci. 33, 7627–7640. https://doi.org/10.1523/JNEUROSCI.0173-13.2013

Keil, B., Blau, J.N., Biber, S., Hoecht, P., Tountcheva, V., Setsompop, K., Triantafyllou, C., Wald, L.L., 2013. A 64-channel 3T array coil for accelerated brain MRI. Magn Reson Med 70, 248–258. https://doi.org/10.1002/mrm.24427

Keil, B., Triantafyllou, C., Hamm, M., Wald, L., 2010. Design optimization of a 32-channel head coil at 7 T; Presented at the In: Proceedings of the Annual Meeting of the International Society for Magnetic Resonance in Medicine; Stockholm. 2010. p. 1493.

Kim, J., Ghadery, C., Cho, S.S., Mihaescu, A., Christopher, L., Valli, M., Houle, S., Strafella, A.P., 2019. Network Patterns of Beta-Amyloid Deposition in Parkinson’s Disease. Mol Neurobiol 56, 7731–7740. https://doi.org/10.1007/s12035-019-1625-z

Kim, R., Nakano, K., Jayaraman, A., Carpenter, M.B., 1976. Projections of the globus pallidus and adjacent structures: An autoradiographic study in the monkey. J. Comp. Neurol. 169, 263–289. https://doi.org/10.1002/cne.901690302

Korte, S.M., Jaarsma, D., Luiten, P.G., Bohus, B., 1992. Mesencephalic cuneiform nucleus and its ascending and descending projections serve stress-related cardiovascular responses in the rat. J Auton Nerv Syst 41, 157–176. https://doi.org/10.1016/0165-1838(92)90137-6

Krout, K.E., Jansen, A.S., Loewy, A.D., 1998. Periaqueductal gray matter projection to the parabrachial nucleus in rat. J Comp Neurol 401, 437–454.

Krout, K.E., Loewy, A.D., 2000. Periaqueductal gray matter projections to midline and intralaminar thalamic nuclei of the rat. J Comp Neurol 424, 111–141. https://doi.org/10.1002/1096-9861(20000814)424:1<111::aid-cne9>3.0.co;2-3

Lee, J., Groh, J.M., 2012. Auditory signals evolve from hybrid-to eye-centered coordinates in the primate superior colliculus. J Neurophysiol 108, 227–242. https://doi.org/10.1152/jn.00706.2011

Lee, M.H., Smyser, C.D., Shimony, J.S., 2013. Resting-State fMRI: A Review of Methods and Clinical Applications. American Journal of Neuroradiology 34, 1866–1872. https://doi.org/10.3174/ajnr.A3263

Leigh RJ, Zee DS, 2006. The Neurology of Eye Movements, Fourth Edition. ed. New York: Oxford University Press.

Li, H., Fang, S., Goni, J., Contreras, J.A., Liang, Y., Cai, C., West, J.D., Risacher, S.L., Wang, Y., Sporns, O., Saykin, A.J., Shen, L., 2015. Integrated Visualization of Human Brain Connectome Data. Brain Inform Health (2015) 9250, 295–305. https://doi.org/10.1007/978-3-319-23344-4_29

Li, W., Yang, C., Wu, S., Nie, Y., Zhang, X., Lu, M., Chu, T., Shi, F., 2018. Alterations of Graphic Properties and Related Cognitive Functioning Changes in Mild Alzheimer’s Disease Revealed by Individual Morphological Brain Network. Front. Neurosci. 12. https://doi.org/10.3389/fnins.2018.00927

Lima, J.C., Oliveira, L.M., Botelho, M.T., Moreira, T.S., Takakura, A.C., 2018. The involvement of the pathway connecting the substantia nigra, the periaqueductal gray matter and the retrotrapezoid nucleus in breathing control in a rat model of Parkinson’s disease. Experimental Neurology 302, 46–56. https://doi.org/10.1016/j.expneurol.2018.01.003

Lima, M.M.S., Andersen, M.L., Reksidler, A.B., Vital, M.A.B.F., Tufik, S., 2007. The role of the substantia nigra pars compacta in regulating sleep patterns in rats. PLoS One 2, e513. https://doi.org/10.1371/journal.pone.0000513

Lima, M.M.S., Reksidler, A.B., Vital, M.A.B.F., 2008. The dopaminergic dilema: Sleep or wake? Implications in Parkinson’s disease. Bioscience Hypotheses 1, 9–13. https://doi.org/10.1016/j.bihy.2008.01.010

Liu, D., Dan, Y., 2019. A Motor Theory of Sleep-Wake Control: Arousal-Action Circuit. Annu Rev Neurosci 42, 27–46. https://doi.org/10.1146/annurev-neuro-080317-061813

Lowe, M.J., Mock, B.J., Sorenson, J.A., 1998. Functional connectivity in single and multislice echoplanar imaging using resting-state fluctuations. Neuroimage 7, 119–132. https://doi.org/10.1006/nimg.1997.0315

Luo, Y.-J., Li, Y.-D., Wang, L., Yang, S.-R., Yuan, X.-S., Wang, J., Cherasse, Y., Lazarus, M., Chen, J.-F., Qu, W.-M., Huang, Z.-L., 2018. Nucleus accumbens controls wakefulness by a subpopulation of neurons expressing dopamine D1 receptors. Nat Commun 9, 1576. https://doi.org/10.1038/s41467-018-03889-3

Mai JK, Paxinos G, 2011. The Human Nervous System, 3rd ed. ed. Academic Press.

Mantyh, P.W., 1983. Connections of midbrain periaqueductal gray in the monkey. II. Descending efferent projections. J Neurophysiol 49, 582–594. https://doi.org/10.1152/jn.1983.49.3.582

Mark, G.P., Shabani, S., Dobbs, L.K., Hansen, S.T., 2011. Cholinergic modulation of mesolimbic dopamine function and reward. Physiology & Behavior 104, 76–81. https://doi.org/10.1016/j.physbeh.2011.04.052

Matelli, M., Luppino, G., Geyer, S., Zilles, K., 2004. CHAPTER 26 - MOTOR CORTEX, in: Paxinos, G., Mai, J.K. (Eds.), The Human Nervous System (Second Edition). Academic Press, San Diego, pp. 973–996. https://doi.org/10.1016/B978-012547626-3/50027-2

May, P.J., 2006a. The mammalian superior colliculus: laminar structure and connections. Prog. Brain Res. 151, 321–378. https://doi.org/10.1016/S0079-6123(05)51011-2

May, P.J., 2006b. The mammalian superior colliculus: laminar structure and connections. Prog Brain Res 151, 321–378. https://doi.org/10.1016/S0079-6123(05)51011-2

Mena-Segovia, J., Bolam, J.P., 2017. Rethinking the Pedunculopontine Nucleus: From Cellular Organization to Function. Neuron 94, 7–18. https://doi.org/10.1016/j.neuron.2017.02.027

Merel, J., Botvinick, M., Wayne, G., 2019. Hierarchical motor control in mammals and machines. Nat Commun 10, 5489. https://doi.org/10.1038/s41467-019-13239-6

Miano, S., Donfrancesco, R., Bruni, O., Ferri, R., Galiffa, S., Pagani, J., Montemitro, E., Kheirandish, L., Gozal, D., Pia Villa, M., 2006. NREM sleep instability is reduced in children with attention-deficit/hyperactivity disorder. Sleep 29, 797–803.

Middleton, F.A., Strick, P.L., 2000. Basal ganglia and cerebellar loops: motor and cognitive circuits. Brain Research Reviews 31, 236–250. https://doi.org/10.1016/S0165-0173(99)00040-5

Mori, F., Okada, K., Nomura, T., Kobayashi, Y., 2016. The Pedunculopontine Tegmental Nucleus as a Motor and Cognitive Interface between the Cerebellum and Basal Ganglia. Front Neuroanat 10, 109. https://doi.org/10.3389/fnana.2016.00109

Moruzzi, G., Magoun, H.W., 1949. Brain stem reticular formation and activation of the EEG. Electroencephalogr Clin Neurophysiol 1, 455–473.

Murakami, M., Ishikura, S., Kominami, D., Shimokawa, T., Murata, M., 2017. Robustness and efficiency in interconnected networks with changes in network assortativity. Appl Netw Sci 2, 1–21. https://doi.org/10.1007/s41109-017-0025-4

Murphy, K., Birn, R.M., Handwerker, D.A., Jones, T.B., Bandettini, P.A., 2009. The impact of global signal regression on resting state correlations: are anti-correlated networks introduced? Neuroimage 44, 893–905. https://doi.org/10.1016/j.neuroimage.2008.09.036

Mutolo, D., 2017. Brainstem mechanisms underlying the cough reflex and its regulation. Respir Physiol Neurobiol 243, 60–76. https://doi.org/10.1016/j.resp.2017.05.008

Nakamura, T., Hillary, F.G., Biswal, B.B., 2009. Resting Network Plasticity Following Brain Injury. PLOS ONE 4, e8220. https://doi.org/10.1371/journal.pone.0008220

Nauta, H.J.W., Cole, M., 1978. Efferent projections of the subthalamic nucleus: An autoradiographic study in monkey and cat. J. Comp. Neurol. 180, 1–16. https://doi.org/10.1002/cne.901800102

Nieuwenhuys, R., Voogd, J., Huijzen, C. van, 2008a. The Human Central Nervous System: A Synopsis and Atlas, 4th ed. Steinkopff-Verlag Heidelberg. https://doi.org/10.1007/978-3-540-34686-9

Nieuwenhuys, R., Voogd, J., Huijzen, C. van, 2008b. The Human Central Nervous System A Synopsis and Atlas.

Noga, B.R., Turkson, R.P., Xie, S., Taberner, A., Pinzon, A., Hentall, I.D., 2017. Monoamine Release in the Cat Lumbar Spinal Cord during Fictive Locomotion Evoked by the Mesencephalic Locomotor Region. Front Neural Circuits 11, 59. https://doi.org/10.3389/fncir.2017.00059

Oakman, S., Faris, P., Kerr, P., Cozzari, C., Hartman, B., 1995. Distribution of pontomesencephalic cholinergic neurons projecting to substantia nigra differs significantly from those projecting to ventral tegmental area. J. Neurosci. 15, 5859– 5869. https://doi.org/10.1523/JNEUROSCI.15-09-05859.1995

Oh, S.W., Harris, J.A., Ng, L., Winslow, B., Cain, N., Mihalas, S., Wang, Q., Lau, C., Kuan, L., Henry, A.M., Mortrud, M.T., Ouellette, B., Nguyen, T.N., Sorensen, S.A., Slaughterbeck, C.R., Wakeman, W., Li, Y., Feng, D., Ho, A., Nicholas, E., Hirokawa, K.E., Bohn, P., Joines, K.M., Peng, H., Hawrylycz, M.J., Phillips, J.W., Hohmann, J.G., Wohnoutka, P., Gerfen, C.R., Koch, C., Bernard, A., Dang, C., Jones, A.R., Zeng, H., 2014. A mesoscale connectome of the mouse brain. Nature 508, 207–214. https://doi.org/10.1038/nature13186

Olszewski, J., Baxter, D., 2014. Cytoarchitecture of the Human Brain Stem, 3rd ed. Karger, Philadelphia, PA: Lippincott.

Olszewski, J., Baxter, D., 1954. Cytoarchitecture of the human brain stem. Cytoarchitecture of the human brain stem.

Omelchenko, N., Sesack, S.R., 2005. Laterodorsal tegmental projections to identified cell populations in the rat ventral tegmental area. J. Comp. Neurol. 483, 217–235. https://doi.org/10.1002/cne.20417

Parent, A., De Bellefeuille, L., 1982. Organization of efferent projections from the internal segment of globus pallidus in primate as revealed by flourescence retrograde labeling method. Brain Research 245, 201–213. https://doi.org/10.1016/0006-8993(82)90802-2

Parvizi, J., Damasio, A., 2001. Consciousness and the brainstem. Cognition 79, 135–160. https://doi.org/10.1016/s0010-0277(00)00127-x

Patriat, R., Molloy, E.K., Meier, T.B., Kirk, G.R., Nair, V.A., Meyerand, M.E., Prabhakaran, V., Birn, R.M., 2013. The effect of resting condition on resting-state fMRI reliability and consistency: A comparison between resting with eyes open, closed, and fixated. NeuroImage 78, 463–473. https://doi.org/10.1016/j.neuroimage.2013.04.013

Pauli, W.M., Nili, A.N., Tyszka, J.M., 2018. A high-resolution probabilistic in vivo atlas of human subcortical brain nuclei. Sci Data 5, 180063. https://doi.org/10.1038/sdata.2018.63

Paxinos G, Xu-Feng H, Sengul G, Watson C, 2012. Organization of brainstem nuclei. In: The Human Nervous System. Elsevier.

Pfaff, D.W., Martin, E.M., Faber, D., 2012. Origins of arousal: roles for medullary reticular neurons. Trends Neurosci 35, 468–476. https://doi.org/10.1016/j.tins.2012.04.008

Poller, W.C., Bernard, R., Derst, C., Weiss, T., Madai, V.I., Veh, R.W., 2011. Lateral habenular neurons projecting to reward-processing monoaminergic nuclei express hyperpolarization-activated cyclic nucleotid-gated cation channels. Neuroscience 193, 205–216. https://doi.org/10.1016/j.neuroscience.2011.07.013

Redinbaugh, M.J., Phillips, J.M., Kambi, N.A., Mohanta, S., Andryk, S., Dooley, G.L., Afrasiabi, M., Raz, A., Saalmann, Y.B., 2020. Thalamus Modulates Consciousness via Layer-Specific Control of Cortex. Neuron 106, 66–75.e12. https://doi.org/10.1016/j.neuron.2020.01.005

Robinson, F.R., Phillips, J.O., Fuchs, A.F., 1994. Coordination of gaze shifts in primates: Brainstem inputs to neck and extraocular motoneuron pools. J. Comp. Neurol. 346, 43– 62. https://doi.org/10.1002/cne.903460104

Rodrigo-Angulo, M.L., Rodr guez-Veiga, E., Reinoso-Su rez, F., 2005. A quantitative study of the brainstem cholinergic projections to the ventral part of the oral pontine reticular nucleus (REM sleep induction site) in the cat. Exp Brain Res 160, 334–343. https://doi.org/10.1007/s00221-004-2015-x

Rosen, B.Q., Halgren, E., 2021. A Whole-Cortex Probabilistic Diffusion Tractography Connectome. eNeuro 8, ENEURO.0416-20.2020. https://doi.org/10.1523/ENEURO.0416-20.2020

Sahib, A.K., Erb, M., Marquetand, J., Martin, P., Elshahabi, A., Klamer, S., Vulliemoz, S., Scheffler, K., Ethofer, T., Focke, N.K., 2018. Evaluating the impact of fast-fMRI on dynamic functional connectivity in an event-based paradigm. PLOS ONE 13, e0190480. https://doi.org/10.1371/journal.pone.0190480

Saint-Cyr, J.A., 1983. The projection from the motor cortex to the inferior olive in the cat. An experimental study using axonal transport techniques. Neuroscience 10, 667–684. https://doi.org/10.1016/0306-4522(83)90209-9

Sakai, S.T., 1988. Corticonigral projections from area 6 in the raccoon. Exp Brain Res 73, 498– 504. https://doi.org/10.1007/BF00406607

Salvador, R., Suckling, J., Coleman, M.R., Pickard, J.D., Menon, D., Bullmore, E., 2005. Neurophysiological architecture of functional magnetic resonance images of human brain. Cereb Cortex 15, 1332–1342. https://doi.org/10.1093/cercor/bhi016

Saper, C.B., Chou, T.C., Scammell, T.E., 2001. The sleep switch: hypothalamic control of sleep and wakefulness. Trends Neurosci 24, 726–731. https://doi.org/10.1016/s0166-2236(00)02002-6

Saper, C.B., Fuller, P.M., Pedersen, N.P., Lu, J., Scammell, T.E., 2010. Sleep state switching. Neuron 68, 1023–1042. https://doi.org/10.1016/j.neuron.2010.11.032

Saper, C.B., Scammell, T.E., Lu, J., 2005. Hypothalamic regulation of sleep and circadian rhythms. Nature 437, 1257–1263. https://doi.org/10.1038/nature04284

Saper, C.B., Stornetta, R.L., 2015. Central Autonomic System, in: The Rat Nervous System. Elsevier, pp. 629–673. https://doi.org/10.1016/B978-0-12-374245-2.00023-1

Sapin, E., Lapray, D., Bérod, A., Goutagny, R., Léger, L., Ravassard, P., Clément, O., Hanriot, L., Fort, P., Luppi, P.-H., 2009. Localization of the Brainstem GABAergic Neurons Controlling Paradoxical (REM) Sleep. PLoS ONE 4, e4272. https://doi.org/10.1371/journal.pone.0004272

Sara, S.J., 2009. The locus coeruleus and noradrenergic modulation of cognition. Nat Rev Neurosci 10, 211–223. https://doi.org/10.1038/nrn2573

Satoh, K., Fibiger, H.C., 1986. Cholinergic neurons of the laterodorsal tegmental nucleus: Efferent and afferent connections. J. Comp. Neurol. 253, 277–302. https://doi.org/10.1002/cne.902530302

Satpute, A.B., Wager, T.D., Cohen-Adad, J., Bianciardi, M., Choi, J.-K., Buhle, J.T., Wald, L.L., Barrett, L.F., 2013. Identification of discrete functional subregions of the human periaqueductal gray. Proc. Natl. Acad. Sci. U.S.A. 110, 17101–17106. https://doi.org/10.1073/pnas.1306095110

Shammah-Lagnado, S.J., Negrão, N., Silva, B.A., Ricardo, J.A., 1987. Afferent connections of the nuclei reticularis pontis oralis and caudalis: a horseradish peroxidase study in the rat. Neuroscience 20, 961–989. https://doi.org/10.1016/0306-4522(87)90256-9

Shinoda, Y., Sugiuchi, Y., Izawa, Y., Hata, Y., 2006. Long descending motor tract axons and their control of neck and axial muscles, in: Progress in Brain Research. Elsevier, pp. 527–563. https://doi.org/10.1016/S0079-6123(05)51017-3

Shook, B. L., Schlag-Rey, M., Schlag, J., 1990. Primate supplementary eye field: I. Comparative aspects of mesencephalic and pontine connections. J. Comp. Neurol. 301, 618–642. https://doi.org/10.1002/cne.903010410

Shook, B.L., Schlag-Rey, M., Schlag, J., 1990. Primate supplementary eye field: I. Comparative aspects of mesencephalic and pontine connections. J. Comp. Neurol 301, 618–642. https://doi.org/10.1002/cne.903010410

Simon, C., Hayar, A., Garcia-Rill, E., 2012. Developmental changes in glutamatergic fast synaptic neurotransmission in the dorsal subcoeruleus nucleus. Sleep 35, 407–417. https://doi.org/10.5665/sleep.1706

Singh, K., García-Gomar, M.G., Bianciardi, M., 2021. Probabilistic Atlas of the Mesencephalic Reticular Formation, Isthmic Reticular Formation, Microcellular Tegmental Nucleus, Ventral Tegmental Area Nucleus Complex, and Caudal-Rostral Linear Raphe Nucleus Complex in Living Humans from 7 Tesla Magnetic Resonance Imaging. Brain Connect. https://doi.org/10.1089/brain.2020.0975

Singh, K., Indovina, I., Augustinack, J.C., Nestor, K., García-Gomar, M.G., Staab, J.P., Bianciardi, M., 2019. Probabilistic Template of the Lateral Parabrachial Nucleus, Medial Parabrachial Nucleus, Vestibular Nuclei Complex, and Medullary Viscero-Sensory-Motor Nuclei Complex in Living Humans From 7 Tesla MRI. Front Neurosci 13, 1425. https://doi.org/10.3389/fnins.2019.01425

Solt, K., Van Dort, C.J., Chemali, J.J., Taylor, N.E., Kenny, J.D., Brown, E.N., 2014. Electrical stimulation of the ventral tegmental area induces reanimation from general anesthesia. Anesthesiology 121, 311–319. https://doi.org/10.1097/ALN.0000000000000117

Stanton, G. B., Goldberg, M.E., Bruce, C.J., 1988. Frontal eye field efferents in the macaque monkey: II. Topography of terminal fields in midbrain and pons. J. Comp. Neurol. 271, 493–506. https://doi.org/10.1002/cne.902710403

Stanton, G.B., Goldberg, M.E., Bruce, C.J., 1988. Frontal eye field efferents in the macaque monkey: II. Topography of terminal fields in midbrain and pons. J. Comp. Neurol 271, 493–506. https://doi.org/10.1002/cne.902710403

Steeves, J.D., Jordan, L.M., 1980. Localization of a descending pathway in the spinal cord which is necessary for controlled treadmill locomotion. Neurosci Lett 20, 283–288. https://doi.org/10.1016/0304-3940(80)90161-5

Stoodley, C.J., Schmahmann, J.D., 2010. Evidence for topographic organization in the cerebellum of motor control versus cognitive and affective processing. Cortex 46, 831– 844. https://doi.org/10.1016/j.cortex.2009.11.008

Stoodley, Catherine J., Schmahmann, J.D., 2010. Evidence for topographic organization in the cerebellum of motor control versus cognitive and affective processing. Cortex 46, 831– 844. https://doi.org/10.1016/j.cortex.2009.11.008

Strassman, A., Highstein, S.M., McCrea, R.A., 1986. Anatomy and physiology of saccadic burst neurons in the alert squirrel monkey. II. Inhibitory burst neurons. J Comp Neurol 249, 358–380. https://doi.org/10.1002/cne.902490304

Szymusiak, R., McGinty, D., 2008. Hypothalamic regulation of sleep and arousal. Ann N Y Acad Sci 1129, 275–286. https://doi.org/10.1196/annals.1417.027

Thomas, C.G., Harshman, R.A., Menon, R.S., 2002. Noise reduction in BOLD-based fMRI using component analysis. Neuroimage 17, 1521–1537. https://doi.org/10.1006/nimg.2002.1200

Tomasi, D., Volkow, N.D., 2011. Functional connectivity hubs in the human brain. Neuroimage 57, 908–917. https://doi.org/10.1016/j.neuroimage.2011.05.024

Uddin, L.Q., Kelly, A.M., Biswal, B.B., Castellanos, F.X., Milham, M.P., 2009. Functional connectivity of default mode network components: correlation, anticorrelation, and causality. Hum Brain Mapp 30, 625–637. https://doi.org/10.1002/hbm.20531

Ulfig, N., Chan, W.Y., 2001. Differential expression of calcium-binding proteins in the red nucleus of the developing and adult human brain. Anat Embryol (Berl) 203, 95–108.

van den Heuvel, M.P., de Lange, S.C., Zalesky, A., Seguin, C., Yeo, B.T.T., Schmidt, R., 2017. Proportional thresholding in resting-state fMRI functional connectivity networks and consequences for patient-control connectome studies: Issues and recommendations. Neuroimage 152, 437–449. https://doi.org/10.1016/j.neuroimage.2017.02.005

van den Heuvel, M.P., Sporns, O., 2013. Network hubs in the human brain. Trends Cogn Sci 17, 683–696. https://doi.org/10.1016/j.tics.2013.09.012

Vertes, R.P., 1991. A PHA-L analysis of ascending projections of the dorsal raphe nucleus in the rat. J Comp Neurol 313, 643–668. https://doi.org/10.1002/cne.903130409

Voogd, J., Shinoda, Y., Ruigrok, T.J.H., Sugihara, I., 2013. Cerebellar Nuclei and the Inferior Olivary Nuclei: Organization and Connections, in: Manto, M., Schmahmann, J.D., Rossi, F., Gruol, D.L., Koibuchi, N. (Eds.), Handbook of the Cerebellum and Cerebellar Disorders. Springer Netherlands, Dordrecht, pp. 377–436. https://doi.org/10.1007/978-94-007-1333-8_19

Wang, J., Wang, X., Xia, M., Liao, X., Evans, A., He, Y., 2015. GRETNA: a graph theoretical network analysis toolbox for imaging connectomics. Front Hum Neurosci 9, 386. https://doi.org/10.3389/fnhum.2015.00386

Watson, R.T., Heilman, K.M., Miller, B.D., King, F.A., 1974. Neglect after mesencephalic reticular formation lesions. Neurology 24, 294–294. https://doi.org/10.1212/WNL.24.3.294

Waung, M.W., Margolis, E.B., Charbit, A.R., Fields, H.L., 2019. A Midbrain Circuit that Mediates Headache Aversiveness in Rats. Cell Rep 28, 2739–2747.https://doi.org/10.1016/j.celrep.2019.08.009

Weber, F., Hoang Do, J.P., Chung, S., Beier, K.T., Bikov, M., Saffari Doost, M., Dan, Y., 2018. Regulation of REM and Non-REM Sleep by Periaqueductal GABAergic Neurons. Nat Commun 9, 354. https://doi.org/10.1038/s41467-017-02765-w

Woolf, N.J., Butcher, L.L., 1986. Cholinergic systems in the rat brain: III. Projections from the pontomesencephalic tegmentum to the thalamus, tectum, basal ganglia, and basal forebrain. Brain Research Bulletin 16, 603–637. https://doi.org/10.1016/0361-9230(86)90134-6

Xi, Z., Luning, W., 2009. REM sleep behavior disorder in a patient with pontine stroke. Sleep Medicine 10, 143–146. https://doi.org/10.1016/j.sleep.2007.12.002

Xiang, H.-B., Zhu, W.-Z., Guan, X.-H., Ye, D.-W., 2013. The cuneiform nucleus may be involved in the regulation of skeletal muscle tone by motor pathway: a virally mediated trans-synaptic tracing study in surgically sympathectomized mice. Brain 136, e251–e251. https://doi.org/10.1093/brain/awt123

Yasui, Y., Saper, C.B., Cechetto, D.F., 1989. Calcitonin gene-related peptide immunoreactivity in the visceral sensory cortex, thalamus, and related pathways in the rat. The Journal of Comparative Neurology 290, 487–501. https://doi.org/10.1002/cne.902900404

Yezierski, R.P., 1988. Spinomesencephalic tract: Projections from the lumbosacral spinal cord of the rat, cat, and monkey. J. Comp. Neurol. 267, 131–146. https://doi.org/10.1002/cne.902670109

Zhang, Y., Larcher, K.M.-H., Misic, B., Dagher, A., 2017. Anatomical and functional organization of the human substantia nigra and its connections. eLife 6, e26653. https://doi.org/10.7554/eLife.26653

Zhong, Y., Wang, H., Lu, G., Zhang, Z., Jiao, Q., Liu, Y., 2009. Detecting functional connectivity in fMRI using PCA and regression analysis. Brain Topogr 22, 134–144. https://doi.org/10.1007/s10548-009-0095-4

Zhu, H., Qiu, C., Meng, Y., Yuan, M., Zhang, Y., Ren, Z., Li, Y., Huang, X., Gong, Q., Lui, S., Zhang, W., 2017. Altered Topological Properties of Brain Networks in Social Anxiety Disorder: A Resting-state Functional MRI Study. Sci Rep 7, 43089. https://doi.org/10.1038/srep43089

