## Supplementary material for "Functional connectome of arousal and motor brainstem nuclei in living humans by 7 Tesla resting-state fMRI": https://figshare.com/s/22cfc0708ab8477c5018

**Supplementary figures and text:**

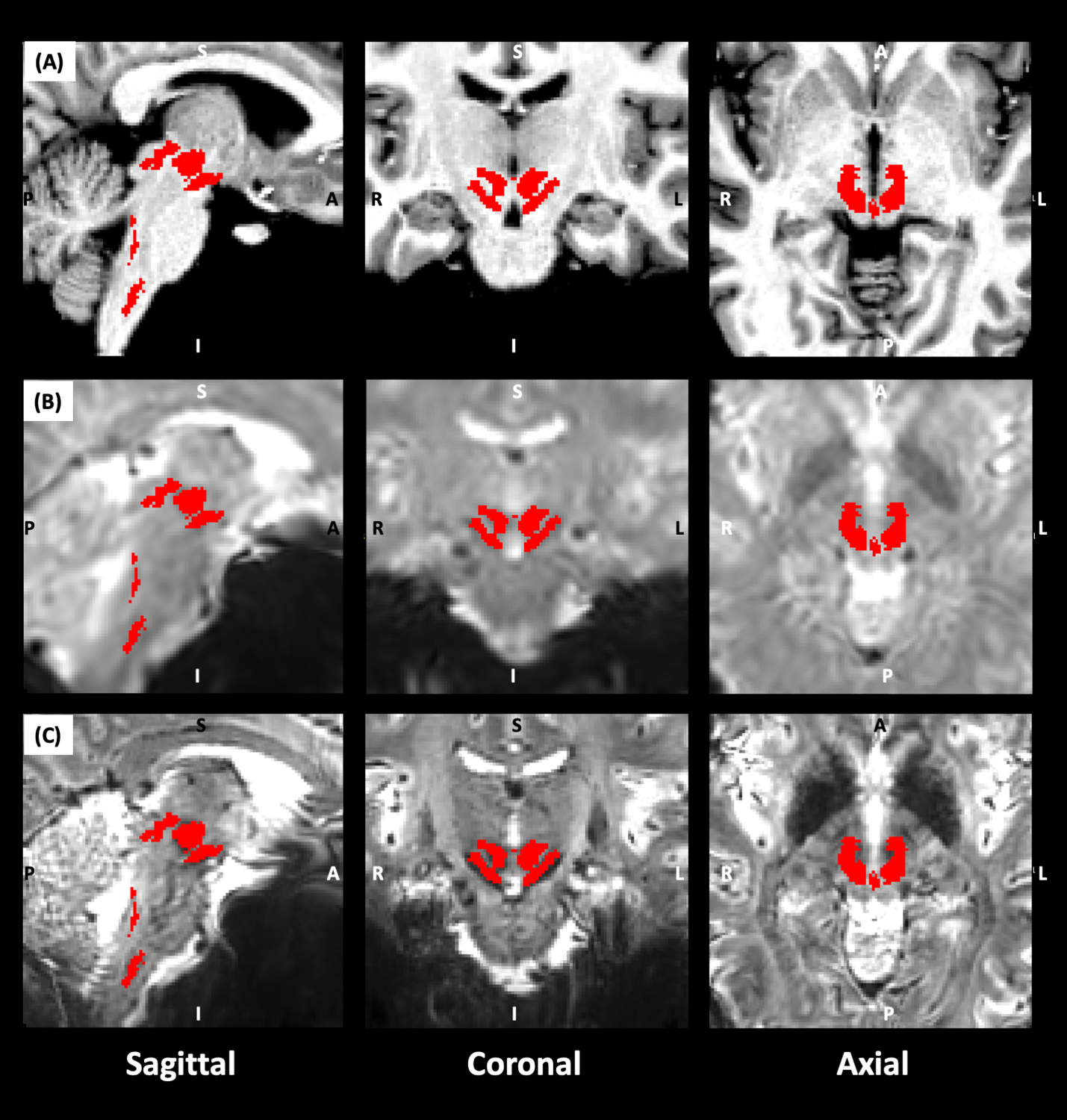

**Supplementary Figure 1.** For an example dataset, we show the alignment of all brainstem seeds (red) on the (A) T_1_–weighted image, (B) 3 Tesla and (C) 7 Tesla pre-processed fMRI.

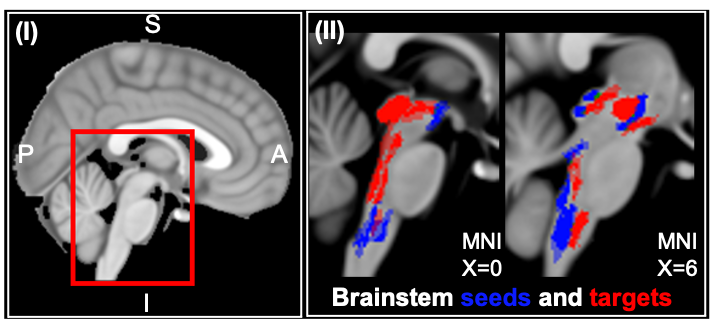

**Supplementary Figure 2.** We show (I) MNI template of brain with inset showing brainstem enlarged in (II) to show brainstem seeds and targets studied in current study.

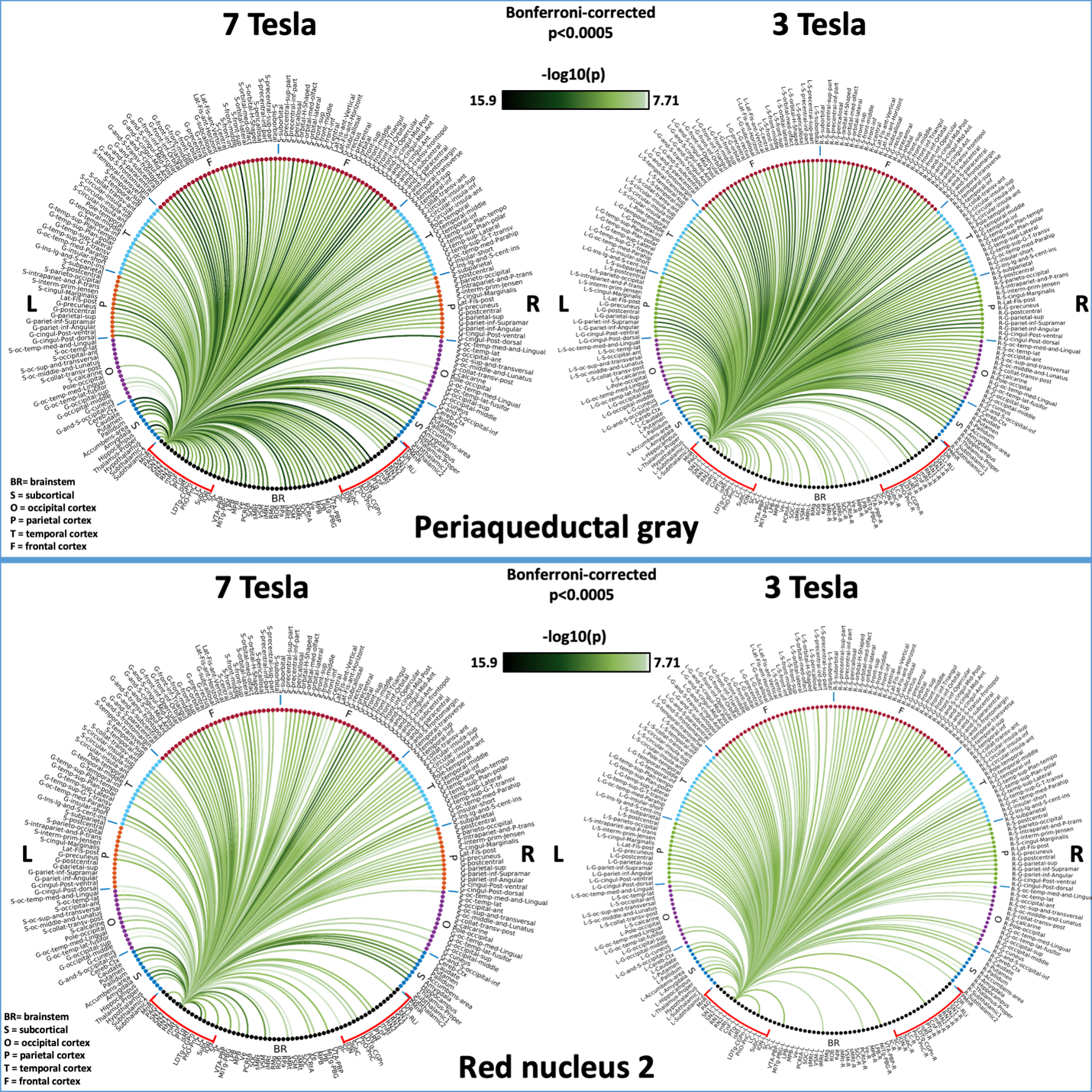

**Supplementary Figure 3:** We show 7 Tesla and 3 Tesla 2D connectome of two representative nuclei (PAG, from the arousal system, and RN1, from the motor system) at p < 0.0005 Bonferroni corrected. We see denser connectivity in the brainstem at 7 Tesla as compared to 3 Tesla.

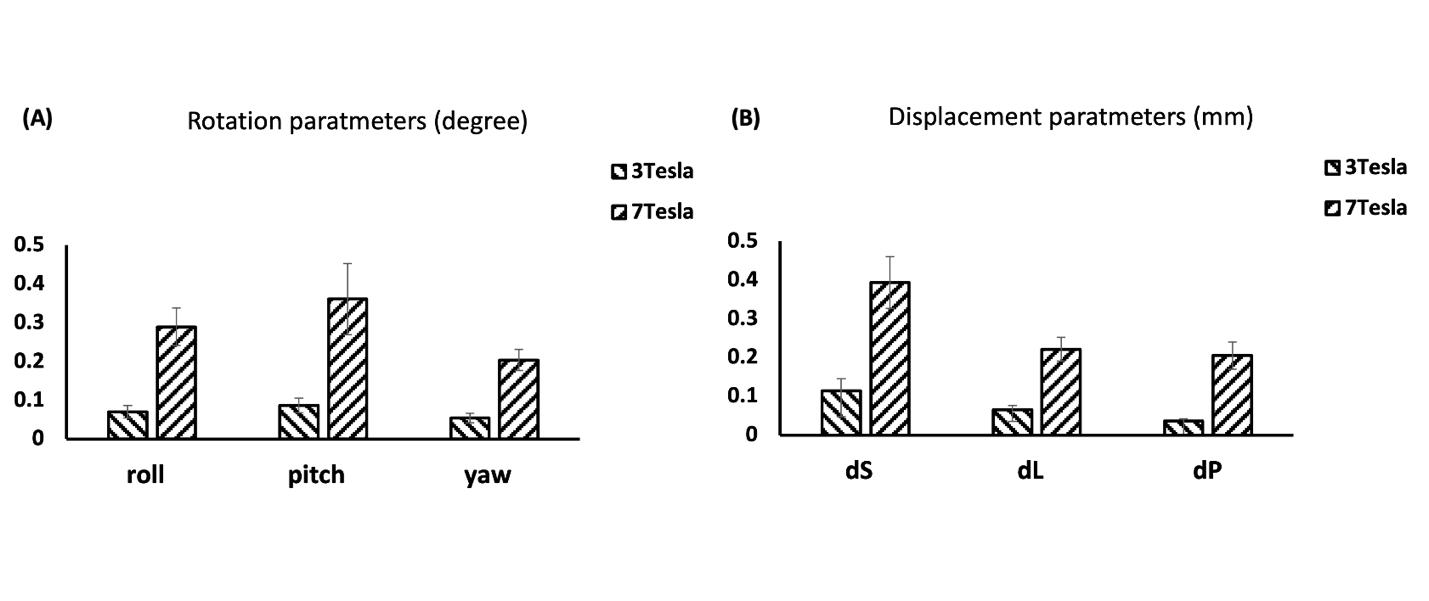

**Supplementary Figure 4.** We show the head motion metric parameters of (A) rotation (degree) and (B) displacement (mm) for 3 Tesla and 7 Tesla fMRI datasets. For each subject, the root mean square value of the motion parameters was computed, and the mean ± s.e. across subjects was displayed.

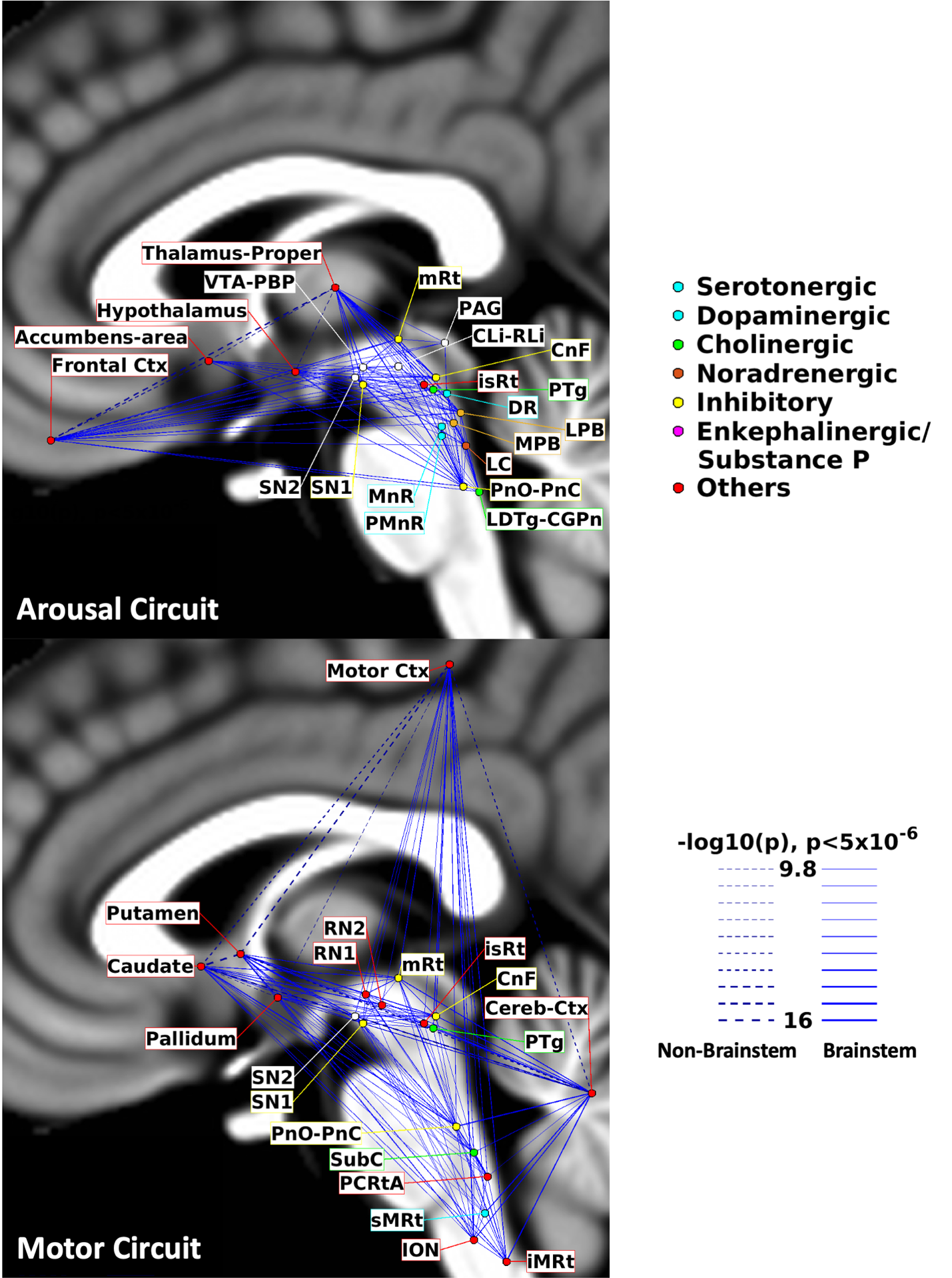

**Supplementary Figure 5. Circuit diagram of the arousal (top) and motor (bottom) network derived from resting-state 3 Tesla fMRI.** The diagram of the arousal network displayed high node interconnectivity. Interestingly, several brainstem arousal nuclei displayed connectivity to the thalamus, hypothalamus, basal forebrain and frontal cortex, as expected for arousal regions. The diagram of the motor network matched expected connectivity of motor regions, including the motor cortex, basal ganglia and cerebellum. Note that the link thickness was varied based on the statistical significance of the connectivity strength. For the arousal circuit, the brainstem nodes were color-coded based on the main neurotransmitter employed. Compared to the 7 Tesla diagrams, several links of the 3 Tesla diagrams were missing or found at lower connectivity strength, especially in the brainstem.

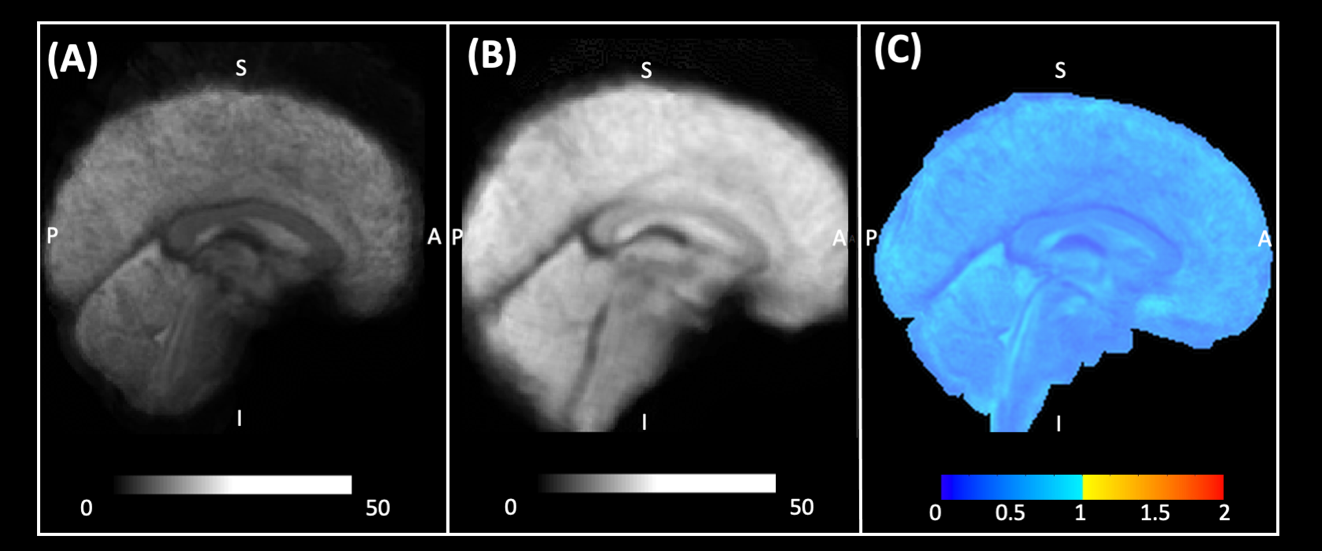

**Supplementary Figure 6:** We show the tSNR image of (A) 7 Tesla fMRI data, (B) 3 Tesla fMRI data, computed after preprocessing in each subject, and averaged across subjects; in (C) we show the ratio of the group average tSNR at 7 Tesla divided by the group average tSNR at 3 Tesla.

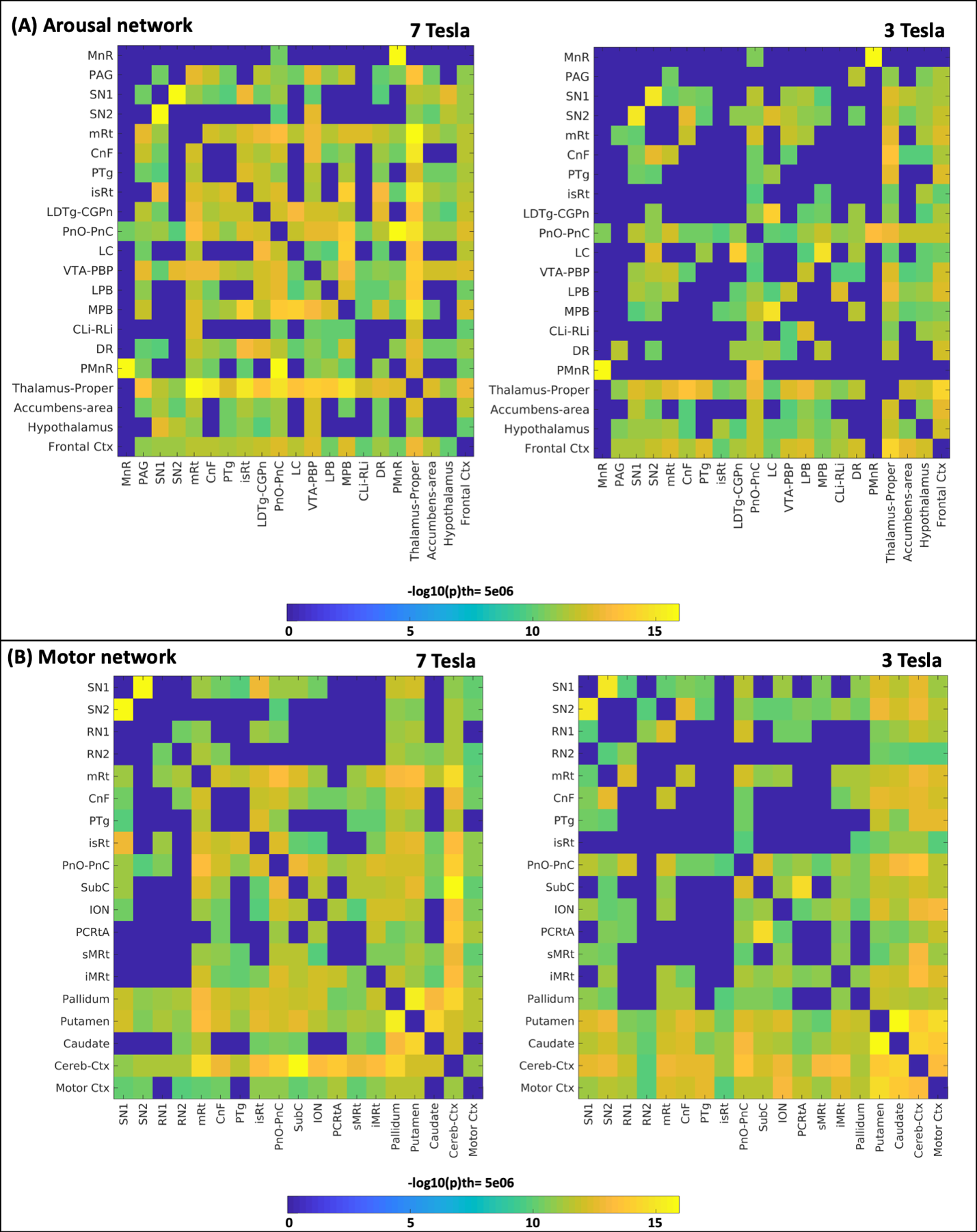

**Supplementary Figure 7** showing the connectivity matrix of arousal and motor nodes of the circuit diagrams of Figure 14 and Supplementary Figure 5, respectively from 7 Tesla and 3 Tesla dataset. The 7 Tesla dataset showed higher connectivity in the brainstem as compared to 3 Tesla. Arousal network nodes showed high connectivity with frontal cortex and related sub-cortical regions both at 7 Tesla and 3 Tesla. Motor network nodes showed high connectivity with motor cortex and striatum regions both at 7 Tesla and 3 Tesla. Note that these matrices were obtained as described in manuscript (section 2.3: Diagram generation), by averaging the connectivity strengths between left and right hemispheres.
